# Nitrogen metabolism profiling reveals cell state-specific pyrimidine synthesis pathway choice

**DOI:** 10.1101/2025.07.23.666448

**Authors:** Milan R. Savani, Bailey C. Smith, Wen Gu, Yi Xiao, Gerard Baquer, Bingbing Li, Skyler S. Oken, Namya Manoj, Lauren G. Zacharias, Vinesh T. Puliyappadamba, Sylwia A. Stopka, Michael S. Regan, Michael M. Levitt, Charles K. Edgar, William H. Hicks, Soummitra Anand, Tracey Shipman, Misty S. Martin-Sandoval, Rainah Winston, João S. Patrício, Xandria Johnson, Trevor S. Tippetts, Diana D. Shi, Andrew Lemoff, Timothy E. Richardson, Pascal O. Zinn, Ashley Solmonson, Thomas P. Mathews, Nathalie Y.R. Agar, Ralph J. DeBerardinis, Kalil G. Abdullah, Samuel K. McBrayer

**Author notes:** Correspondence (K.G.A.), (S.K.M.).

## Abstract

Conventional stable isotope tracing assays track one or several metabolites. However, cells use an array of nutrients to sustain nitrogen metabolic pathways. This incongruency hampers a system level understanding of cellular nitrogen metabolism. Therefore, we created a platform to simultaneously trace 30 nitrogen isotope-labeled metabolites. This platform revealed that while primitive cells engage both de novo and salvage pyrimidine synthesis pathways, differentiated cells nearly exclusively salvage uridine despite expressing de novo pathway enzymes. This link between cell state and pyrimidine synthesis routes persisted in physiological contexts, including primary murine and human tissues and tumor xenografts. Mechanistically, we found that Ser1900 phosphorylation of CAD, the first enzyme of the de novo pathway, was enriched in primitive cells and that mimicking this modification in differentiated cells abrogated their preference for pyrimidine salvage. Collectively, we establish a method for nitrogen metabolism profiling and define a mechanism of cell state-specific pyrimidine synthesis pathway choice.

## INTRODUCTION

Over the past two decades, technical advances have broadened understanding of the influence of metabolism on physiology and disease. Metabolism can be probed by steady-state metabolite profiling or stable isotope tracing. Steady-state metabolite profiling measures static metabolite pool sizes while stable isotope tracing captures dynamic metabolic activities^1^. In tracing experiments, a stable isotope (typically ^13^C or ^15^N) is incorporated into a nutrient and its metabolic fate is then tracked throughout biochemical networks. Metabolites are then quantified by magnetic resonance spectroscopy or mass spectrometry. High magnetic field instruments and new chemical probes, including hyperpolarized probes with increased signal, have created new opportunities to survey metabolism in vivo^2^. Likewise, development of high resolution mass spectrometers has expanded the number of metabolites that can be quantified and enhanced discrimination between isotopes of different atoms^3,4^.

The impact of these advances, however, has not manifested evenly across biochemical pathways. For instance, in vivo tracing studies involving nutrients labeled with ^13^C isotopes have changed our understanding of central carbon metabolism in organs and tumors^5–9^. In contrast, the mechanisms used by cells and tissues to incorporate nitrogen into the metabolome remain obscure^10^. This difference is largely attributable to the greater diversity of nutrients cells use to support nitrogen metabolism compared to carbon metabolism^10,11^. Stable isotope tracing evaluating carbon metabolism yields broad coverage by tracing few or single ^13^C-labeled nutrients (such as glucose, glutamine, lactate, acetate, and free fatty acids)^4^. To similarly cover nitrogen metabolic pathways, many ^15^N-labeled tracers would be required. Because current experimental paradigms are not tailored for multiplexed stable isotope tracing, our understanding of cellular nitrogen metabolism programs is poor relative to carbon metabolism.

To address this limitation, we sought to develop a method for parallelized stable isotope tracing. We hypothesized that we could generate a system level portrait of nitrogen metabolism programs using a library of medium preparations to simultaneously trace dozens of ^15^N-labeled nutrients. Moreover, we posited that we could leverage human plasma-like medium (HPLM) to maximize the physiologic relevance of the findings generated by this method^12,13^. Our previous research using stable isotope tracing to comparatively assess substrate preferences for nucleotide biosynthesis in glioma cells provided a template for this effort^14^. We thus endeavored to create a nitrogen metabolism profiling platform and companion computational pipeline that could reveal fundamental insights into the function and regulation of biochemical pathways involving nitrogenous metabolites.

## RESULTS

### Development and validation of a nitrogen metabolism profiling platform

Cells use an array of nitrogen-containing nutrients to produce metabolites including nucleotides^15^, amino acids^11^, heme^16^, creatine^17^, amino sugars, nucleotide sugars^18^, glutathione^19^, and polyamines^20^. To comprehensively map contributions of individual nutrients to nitrogen metabolic pathways, we first created a library of 30 HPLM stocks containing stable isotope tracers (Figure 1A). In each, we replaced a single, unlabeled nitrogenous metabolite with its ^15^N isotope-labeled counterpart. We included tracers corresponding to the most abundant nitrogenous compounds in human plasma: all 20 proteinogenic amino acids, non-proteinogenic amino acids (taurine), urea cycle intermediates (ornithine and citrulline), nucleotide precursors (hypoxanthine and uridine), ions (nitrate and ammonium), and end products of nitrogen metabolism (urea and uric acid) (Table S1). We also generated tracer-free HPLM as a negative control. We then could perform parallelized stable isotope tracing assays in paired cultures to elucidate the impact of individual variables (such as drug treatments, genetic mutations, cell states) on nitrogen metabolism. After analyzing metabolite extracts using high-resolution liquid-chromatography mass spectrometry^21,22^, we built a computational pipeline to quantify and deconvolute the resulting rich dataset. This pipeline produces quantitative labeling data for each tracer-metabolite pair (30 ^15^N-labeled nutrients each traced into >100 nitrogen-containing metabolite pools) and leverages these data to generate: 1) Sankey diagrams^23^ describing global nitrogen metabolism, 2) a rank-ordered list of differences in tracer metabolism between cultures (reflected by “differential labeling score”, and 3) steady-state metabolite levels in each condition.

**Figure 1.**
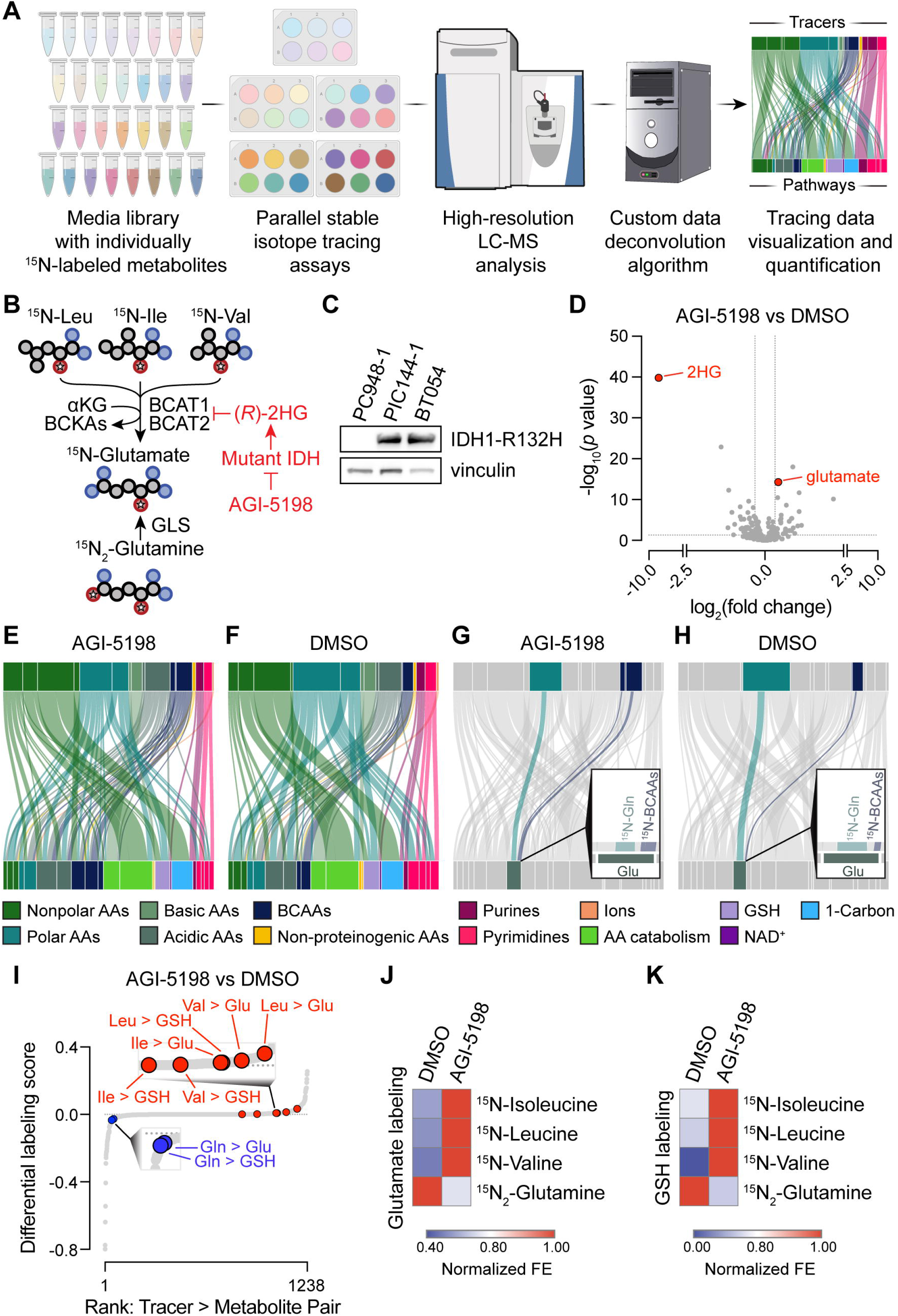
Development and validation of a nitrogen metabolism profiling platform. **(A)** Schema depicting nitrogen metabolism profiling platform. **(B)** Schema of branched-chain amino acid aminotransferase (BCAT)- or glutaminase (GLS)-dependent nitrogen transfer. αKG = alpha-ketoglutarate. BCKA = branched chain ketoacids. **(C)** Representative immunoblot of IDH1-R132H expression in BT054 and murine primary glioma cell lines derived from IDH1 WT (PC: *Pik3ca^mut^*, *Trp53^mut^*, and *Atrx^mut^*) or IDH1-R132H mutant (PIC: *Idh1^R^*^132^*^H^*, *Pik3ca^mut^*, *Trp53^mut^*, and *Atrx^mut^*) glioma stem-like cell lines. **(D-K)** Nitrogen metabolism profiling platform in BT054s treated with 3µM AGI-5198 or DMSO for 72 hours, followed by tracing for 18 hours. (*n* = 1 per tracer per treatment). (D) Volcano plot of metabolite levels (*n* = 30). 2HG = 2-hydroxyglutarate. (E-H) Sankey diagrams depicting labeling from ^15^N-labeled tracers (top bars) to representative intermediates of metabolic pathways (bottom bars) in (E, G) AGI-5198 or (F, H) DMSO conditions. Width of connecting lines indicates the fractional enrichment. Insets in (G-H) highlight branched chain amino acid (BCAA) or glutamine contribution to glutamate. AA = amino acid. GSH = reduced glutathione. NAD^+^ = nicotinamide adenine dinucleotide. **(I)** Waterfall plot ranking differential labeling scores. **(J-K)** Heatmap of label from indicated tracer (y-axis) in (J) glutamate or (K) GSH in each treatment (x-axis). Fractional enrichments normalized to condition with maximal labeling for each tracer. Two-tailed *p* values were determined by unpaired *t*-test. See also Figure S1.

After preparing the HPLM library, we confirmed selective labeling of the metabolite pool of interest in each stock (Figure S1A). Isotope enrichment was not observed in tracer-free medium, indicating specificity of these patterns. We did not detect cysteine (likely due to spontaneous oxidation to cystine^24^) nor quantify ammonia, nitrate, or urea [due to their low mass-to-charge (m/z) values]. We also found minimal variance across medium stocks in cognate metabolite pool size for each tracer (Figure S1B). We next sought to validate our platform by attempting to “rediscover” an established mechanism of nitrogen metabolism dysregulation. Previously, we reported that (*R*)-2-hydroxyglutarate [(*R*)-2HG], the product of mutant IDH oncoproteins, competitively inhibits branched chain amino acid transaminase (BCAT)-dependent glutamate synthesis^25^ (Figure 1B). To compensate, IDH-mutant tumor cells activate glutaminase (GLS)-dependent glutamate synthesis. Treatment of these cells with a mutant IDH inhibitor (such as AGI-5198^26^) causes (*R*)-2HG levels to fall, derepressing BCAT enzymes and reestablishing a balance between branched chain amino acids and glutamine as nitrogen donors to glutamate. Because glutamate is incorporated into glutathione (GSH), similar relationships between amino acid substrate preference and (*R*)-2HG abundance also apply to GSH synthesis. Using BT054, IDH1-mutant glioma stem-like cells (GSCs) derived from a patient with Grade 3 oligodendroglioma^27^, we tested whether nitrogen metabolism profiling could uncover these metabolic interactions in an unbiased manner.

We first confirmed endogenous expression of the IDH1-R132H mutant enzyme in BT054 cells, benchmarking against levels in isogenic murine GSCs from genetically engineered mouse models of IDH-mutant and IDH-wildtype astrocytoma^14,28^ (Figure 1C). Next, we treated BT054 with dimethyl sulfoxide (DMSO) vehicle or AGI-5198 to create (*R*)-2HG^high^ and (*R*)-2HG^low^ conditions, respectively, prior to culturing them in our HPLM library for 18 hours. We confirmed successful labeling of intracellular metabolite pools in each tracer-containing medium stock (Figure S1C) and expected effects of AGI-1598 on intracellular metabolite pool sizes (i.e., 2HG depletion and glutamate upregulation, Figure 1D and Table S2). Finally, we employed our computational pipeline to systematically assess how (*R*)-2HG accumulation affects nitrogen metabolism (Figure S1D). We constructed Sankey diagrams to provide a system level view in which tracers and metabolites representing individual biochemical pathways (Table S1) at the top and bottom of these plots, respectively, are connected by lines of widths correlating with labeling between them (Figures 1E and 1F). Both ^15^N-labeled tracers and metabolic pathways were grouped and color-coded by function. We then highlighted contributions of ^15^N_2_-glutamine and ^15^N-BCAA tracers to glutamate (Figures 1G and 1H). Consistent with our prior hypothesis-driven work^25^ (Figure 1B), nitrogen metabolism profiling revealed that (*R*)-2HG accumulation diminishes BCAA-dependent glutamate production and stimulates compensatory glutamine-dependent glutamate synthesis (Figure 1H). Under (*R*)-2HG^low^ conditions, BCAT enzymes are derepressed and BCAAs and glutamine contribute similarly to glutamate synthesis (Figure 1G)^25^.

Other platform outputs also captured these (*R*)-2HG-dependent differences in glutamate metabolism. Differential labeling scores reflecting metabolism of ^15^N-BCAAs to glutamate and glutathione were elevated in AGI-5198-treated cells whereas those for ^15^N_2_-glutamine-dependent labeling of these metabolites were decreased (Figure 1I, full dataset in Table S2). Differential labeling scores reflected marked differences in BCAA and glutamine metabolism between the two treatment conditions (Figures 1J and 1K). Steady-state metabolomics analysis demonstrated increased glutamate and glutathione levels as (*R*)-2HG declined (Figures S1E-G), consistent with differences in BCAT activity. We performed low-throughput stable isotope tracing experiments to validate these findings, showing that AGI-5198 treatment increases BCAA labeling of glutamate and glutathione pools (Figures S1H and S1I). These data validate platform performance and suggest that it may be useful for unbiased discovery of novel nitrogen metabolism programs.

### Nitrogen metabolism profiling reveals a link between pyrimidine synthesis pathway preference and cell state

Following development and validation of our platform, we sought to apply this tool to a fundamental question: how do nitrogen metabolic programs differ between malignant and nonmalignant cells? We therefore profiled BT054 GSCs and NHA immortalized human astrocytes^29^. These lines differ in both malignancy and differentiation state: BT054s are primitive and transformed whereas NHAs are differentiated and non-tumorigenic. We first optimized plating to avoid media nutrient depletion (Figure S2A) and confirmed intracellular tracer accumulation (Figure S2B). Myriad differences in nitrogen metabolism programs existed between these lines (Figures 2A and 2B). One difference entailed substrate preference for pyrimidine nucleotide synthesis (Figures 2C-E, full dataset in Table S3). NHA cells nearly exclusively used uridine to sustain uridine monophosphate (UMP), uridine diphosphate (UDP), and uridine triphosphate (UTP) pyrimidine pools while BT054 cells predominantly used glutamine. Pyrimidine nucleotides can be produced by de novo synthesis (for which glutamine, aspartate, and bicarbonate serve as substrates) or uridine-dependent salvage^15^ (Figure 2F). Therefore, NHA cells rely principally on pyrimidine salvage and BT054 cells engage both de novo and salvage pathways. This interpretation was supported by steady-state metabolomics revealing that de novo pathway intermediates and pyrimidine nucleotides were elevated in BT054 relative to NHA cells whereas uridine was diminished (Figures S2C and S2E-K and Table S3). Importantly, enhanced de novo pyrimidine nucleotide synthesis could not be explained by the demands of rapid proliferation because NHA divided more quickly than BT054^14^ (Figure S2D).

**Figure 2.**
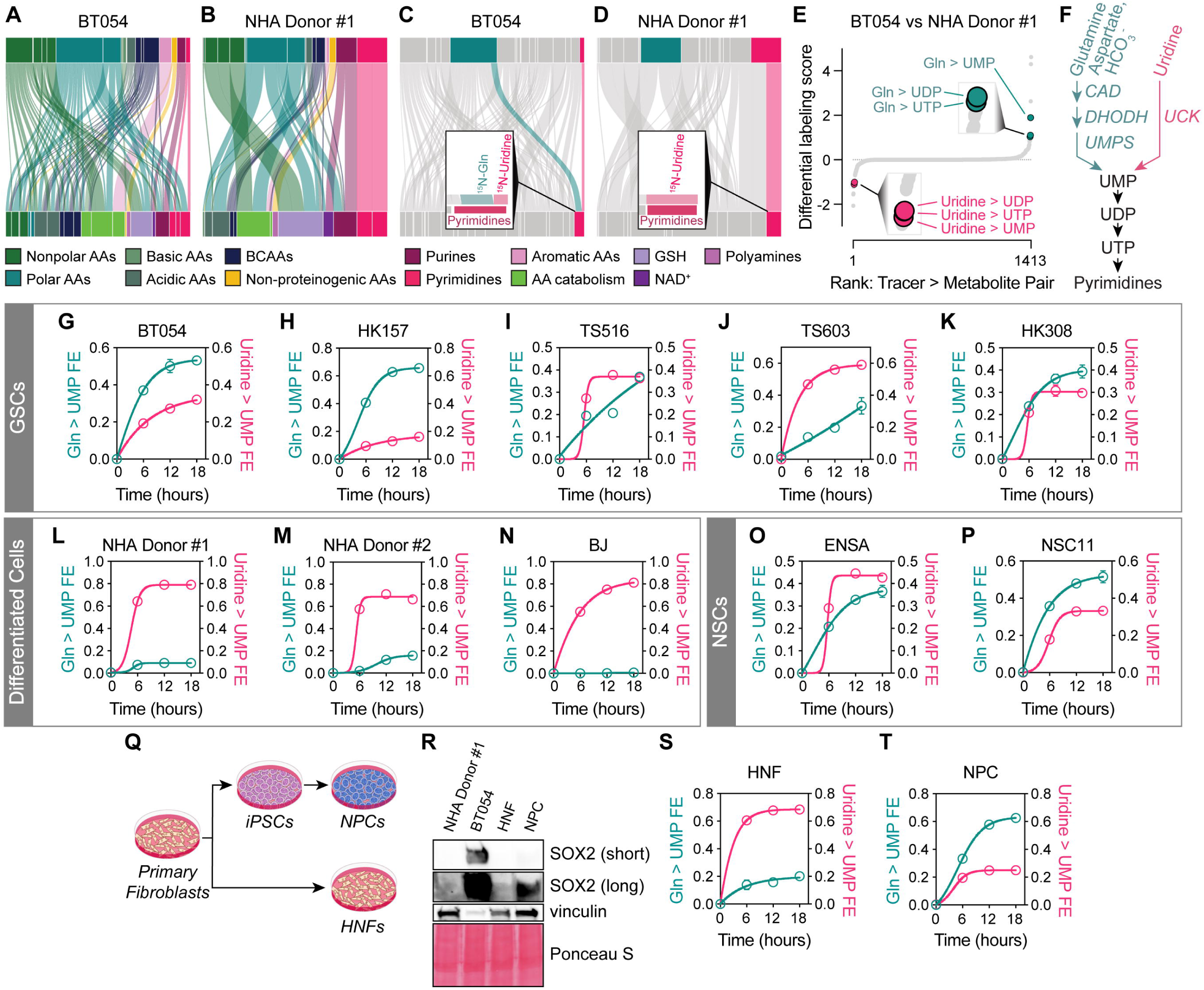
Nitrogen metabolism profiling reveals link between pyrimidine synthesis pathway choice and differentiation state. **(A-E)** Nitrogen metabolism profiling in BT054 and NHA Donor #1 traced for 18 hours (*n* = 1 per tracer per line). (A-D) Sankey diagrams depicting labeling from ^15^N-labeled tracers (top bars) to representative intermediates of metabolic pathways (bottom bars) in (A, C) BT054 or (B, D) NHA. Insets in (C-D) highlight glutamine and uridine contribution to pyrimidines. AA = amino acid. BCAAs = branched chain amino acids. GSH = reduced glutathione. NAD^+^ = nicotinamide adenine dinucleotide. (E) Waterfall plot ranking differential labeling scores. UMP = uridine monophosphate. UDP = uridine diphosphate. UTP = uridine triphosphate. **(F)** Schema of pyrimidine synthesis pathways. **(G-P)** Amide-^15^N-glutamine and ^15^N_2_-uridine tracing to UMP in (G-K) glioma stem-like cell (GSC), (L-N) differentiated, or (O-P) neural stem cell (NSC) lines (*n* = 3 for all except HK308, for which *n* = 2). FE = fractional enrichment. **(Q)** Schema of neural progenitor cell (NPC) derivation from primary human fibroblasts. HNFs = human normal fibroblasts. **(R)** Representative immunoblot for SOX2 in NHA Donor #1, BT054, HNF, and NPC. **(S-T)** Amide-^15^N-glutamine and ^15^N_2_-uridine tracing to UMP in (S) HNF and (T) NPC. (*n* = 3). All amide-^15^N-glutamine to UMP labeling data represent UMP M+1 FE normalized to glutamine M+1 FE. All ^15^N_2_-uridine to UMP labeling data represent UMP M+2 FE. For all panels, data are means ± SEM. See also Figure S2.

De novo pyrimidine synthesis is aberrantly activated in a spectrum of human cancers^30^. Accordingly, this pathway is being investigated as a tumor-selective vulnerability in several contexts^14,31–40^. However, the determinants of pyrimidine synthesis pathway choice in individual cells are not well defined. We profiled BT054 and NHA under identical nutrient conditions, implying that pyrimidine synthesis pathway preference is cell-autonomous. To investigate this idea, we evaluated de novo and salvage pyrimidine synthesis pathways in a panel of cell lines by quantifying UMP labeling from amide-^15^N-glutamine and ^15^N_2_-uridine tracers. GSCs derived from both IDH-wildtype glioblastomas (GBMs) and IDH-mutant gliomas constitutively activated de novo pyrimidine synthesis (Figures 2G-2K). Differentiated, non-transformed cells (including immortalized astrocytes derived from two different donors and BJ fibroblasts), however, had minimal de novo and robust salvage pathway flux (Figures 2L-2N). To ask if these differences associated with differentiation state or malignancy, we also analyzed two non-transformed yet primitive neural stem cell lines. These lines displayed comparable de novo pyrimidine synthesis activity to GSCs (Figures 2O and 2P), suggesting that de novo pathway activation is principally associated with differentiation state rather than malignancy.

To mitigate confounding effects of genetic differences between lines, we created isogenic primitive and differentiated cell cultures then compared their pyrimidine synthesis programs. We expanded differentiated human normal fibroblasts (HNFs) or reprogrammed them to induced pluripotent stem cells (iPSCs) and then to primitive neural progenitor cells (NPCs) (Figure 2Q). We confirmed differentiation state by immunoblotting for the primitive cell marker SOX2^41^ (Figure 2R). NPCs recapitulated the preference for de novo pyrimidine synthesis observed in GSCs and NSCs while HNFs, like astrocytes and other fibroblasts, predominantly relied on pyrimidine salvage (Figures 2S and 2T). Differential pyrimidine synthesis programs in GSCs, differentiated cells, and NSCs/NPCs were not explained by tracer accumulation (Figures S2L). Moreover, neither growth factor/serum content of culture media nor adherent/suspension growth conditions (factors that distinguish differentiated and primitive cell cultures) affected pyrimidine synthesis programs (Figures S2M-P). Taken together, our findings show that differentiation state is a key cell-intrinsic determinant of pyrimidine synthesis pathway choice.

### Cellular differentiation states are associated with distinct pyrimidine synthesis programs in primary brain tissue

We next asked if findings from our platform are relevant to primary brain tissue metabolism. Gliomas are enriched in primitive cell populations^42–44^ while adult brain parenchyma largely comprises terminally differentiated neural cells^45^. We exploited this distinction to ask if cell state-specific differences in pyrimidine synthesis are observed in vivo. We randomized naïve mice or mice bearing MGG152 orthotopic patient-derived xenografts of IDH-mutant glioma to continuous infusions of amide-^15^N-glutamine, ^15^N_2_-uridine, or saline and performed conventional and spatial metabolomics on brain tissues (Figure 3A). Approximately 40% of plasma glutamine and uridine pools were labeled in tracer-infused mice, leading to ∼20-25% and ∼15% labeling of glutamine and uridine, respectively, in non-neoplastic brain tissue and glioma xenografts (Figures 3B-3E). Labeling was specific because isotope enrichment was negligible in saline-infused mice. Glutamine infusion did not elevate plasma or brain glutamine levels (Figures S3A and S3B). Although uridine infusion increased plasma uridine, levels in brain tissue were not affected (Figures S3C and S3D). Therefore, our experimental approach did not artificially alter substrate availability.

**Figure 3.**
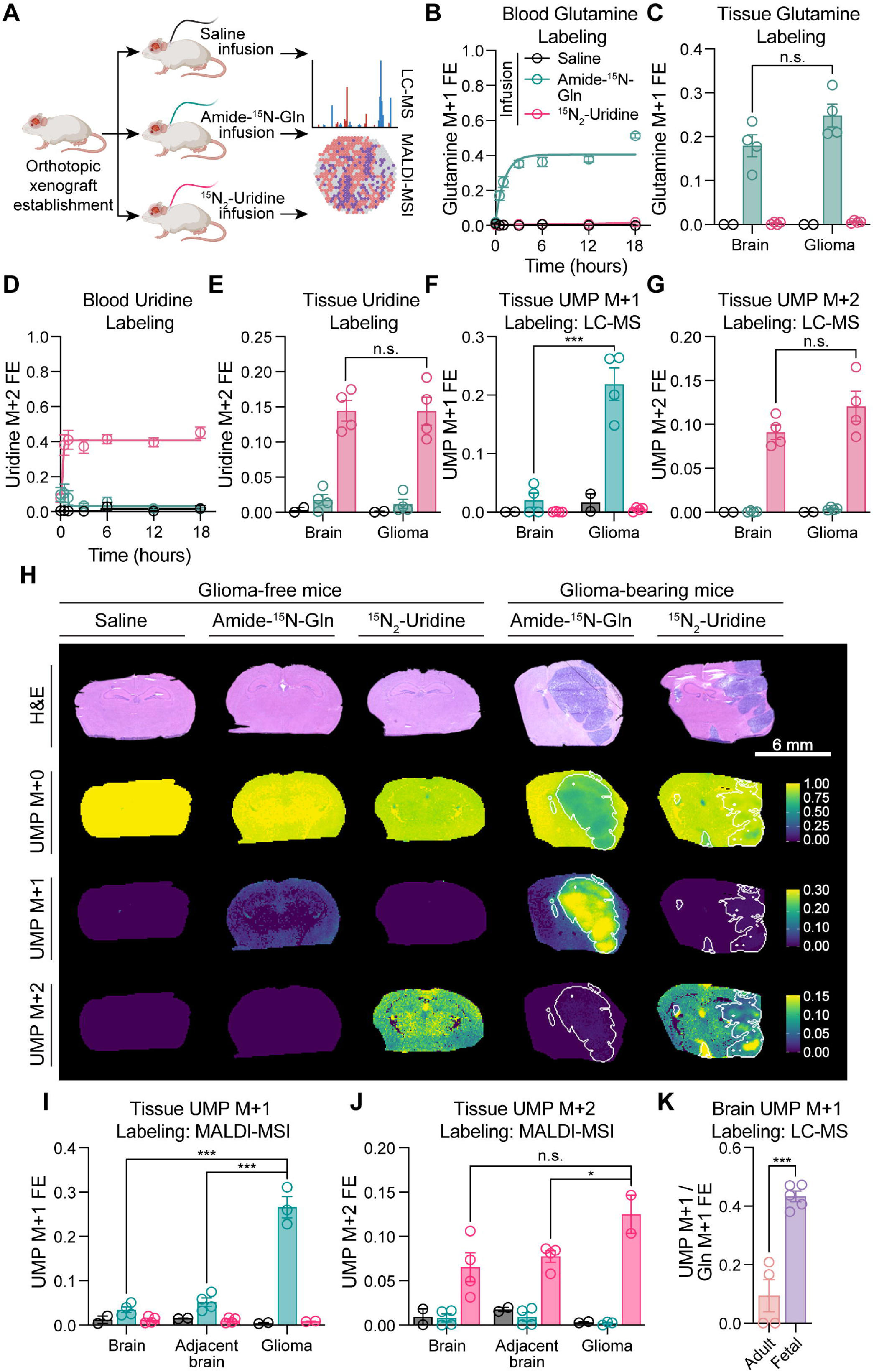
Tissues enriched in primitive cells display constitutive de novo pyrimidine synthesis in vivo. **(A)** Schema depicting infusion with saline, amide-^15^N-glutamine, or ^15^N_2_-uridine for 18 hours in mice bearing MGG152 glioma orthotopic xenografts or non-tumor-bearing mice. **(B-C)** Glutamine M+1 FE in (B) blood or (C) tissue (*n* = 2 for saline, *n* = 4 for amide-^15^N-glutamine and ^15^N_2_-uridine). **(D-E)** Uridine M+2 FE in (D) blood or (E) tissue (*n* = 2 for saline, *n* = 4 for amide-^15^N-glutamine and ^15^N_2_-uridine). **(F-G)** UMP (F) M+1 or (G) M+2 FE in normal brain or glioma xenograft tissues (*n* = 2 for saline, *n* = 4 for amide-^15^N-glutamine and ^15^N_2_-uridine). **(H)** Representative brain sections from mice bearing MGG152 xenografts. Top: hematoxylin and eosin (H&E) stain. UMP M+0 (second row), M+1 (third row), or M+2 (bottom) FE determined by matrix-assisted laser desorption ionization mass spectrometry imaging (MALDI-MSI). Scale bar = 6 mm. **(I-J)** MALDI-MSI-based quantification of UMP (I) M+1 or (J) M+2 FE in indicated tissues (*n* = 2 for tissues from saline-infused mice and for xenograft gliomas from ^15^N_2_-uridine-infused mice; *n* = 3 for xenograft gliomas from amide-^15^N-glutamine-infused mice; *n* = 4 for all others). **(K)** UMP M+1 FE relative to glutamine M+1 FE in adult mouse brains infused for 18 hours or fetal mouse brains from dams infused for 5 hours (*n* = 4 for adult and *n* = 5 for fetal). n.s. = not significant, \**p* < 0.05, \*\*\**p* < 0.001. Two-tailed *p* values were determined by unpaired *t*-test. For all panels, data are means ± SEM. See also Figure S3.

Although glutamine tracer accumulation was similar in normal brain and glioma, robust glutamine-dependent labeling of UMP was only present in glioma (Figure 3F) while uridine-dependent labeling of UMP was similar in normal brain and glioma (Figure 3G). Glutamine labeled metabolites other than UMP in normal brain, including CDP-ethanolamine (Figures S3F and S3G). Isotope enrichment in the M+1 isotopologue of CDP-ethanolamine occurs through two enzymatic activities: 1) synthesis of CTP from glutamine and UTP by CTP synthase, and 2) synthesis of CDP-ethanolamine from CTP and phosphoethanolamine by CTP-phosphoethanolamine cytidylyltransferase (Figures S3E, S3H, and S3I). Evidence of glutamine metabolism by CTP synthase (and other enzymes) suggests that global repression of glutamine catabolism did not cause low UMP labeling by glutamine. Rather, these data reflect specific inactivation of de novo pyrimidine synthesis.

We also performed spatial metabolomics analysis of brain parenchyma from tracer-infused naïve or glioma-bearing mice using matrix-assisted laser desorption ionization mass spectrometry imaging (MALDI-MSI). Normal brain tissue analyzed by MALDI-MSI from naïve mice and tumor-adjacent brain tissues from glioma-bearing mice also showed lower levels of de novo pyrimidine synthesis than glioma xenografts (Figures 3H and 3I). Uridine-dependent pyrimidine salvage was again constitutively active in all tissues (Figures 3H and 3J). Collectively, these data support a tight coupling of differentiation state to de novo pyrimidine synthesis regulation.

We next compared de novo pyrimidine synthesis in adult and fetal brain tissues. Fetal brain contains more undifferentiated cells, including neural stem cells, oligodendrocyte precursor cells, and neuroblasts, than adult tissue^45^. Consistent with this large population of primitive cells, fetal brain tissue recapitulated the elevated levels of de novo pyrimidine synthesis observed in glioma xenografts (Figure 3K).

We next asked if pyrimidine synthesis pathway choice observed in mouse brain and orthotopic xenografts extended to primary human brain samples. We first analyzed steady-state metabolite profiling data from a collection of 50 human brain specimens. The de novo pyrimidine synthesis pathway intermediate carbamoyl aspartate was increased in both high grade gliomas (HGGs) and lower grade gliomas (LGGs) relative to non-malignant brain tissue, consistent with pathway activation (Figure 4A). In contrast, uridine did not differ between these tissues (Figure 4B). We previously developed an approach to culture primary brain tissue explants using a customized formulation of HPLM that preserves viability and key biological features of resident cell populations^46^. Leveraging this technique, we sought to assess pyrimidine synthesis in human brain tissues using stable isotope tracing. We generated organoid explants from non-malignant brain parenchyma and glioma using stereotactic neurosurgical navigation to target specific brain regions. We then immediately cultured these tissues in HPLM containing amide-^15^N-glutamine, ^15^N_2_-uridine, or no tracer (Figure 4C). We collected 2 metastasis-adjacent non-malignant brain or 5 glioma samples from patients who underwent surgical tumor resection (Figure 4D, Table S4). We also collected non-malignant brain parenchyma from 3 patients without oncologic disease who underwent ventriculoperitoneal shunt placement. After establishing explants, we stained remaining tissue with hematoxylin and eosin (H&E) and a board-certified neuropathologist (T.E.R.) confirmed presence or absence of tumor cells (Figure 4E).

**Figure 4.**
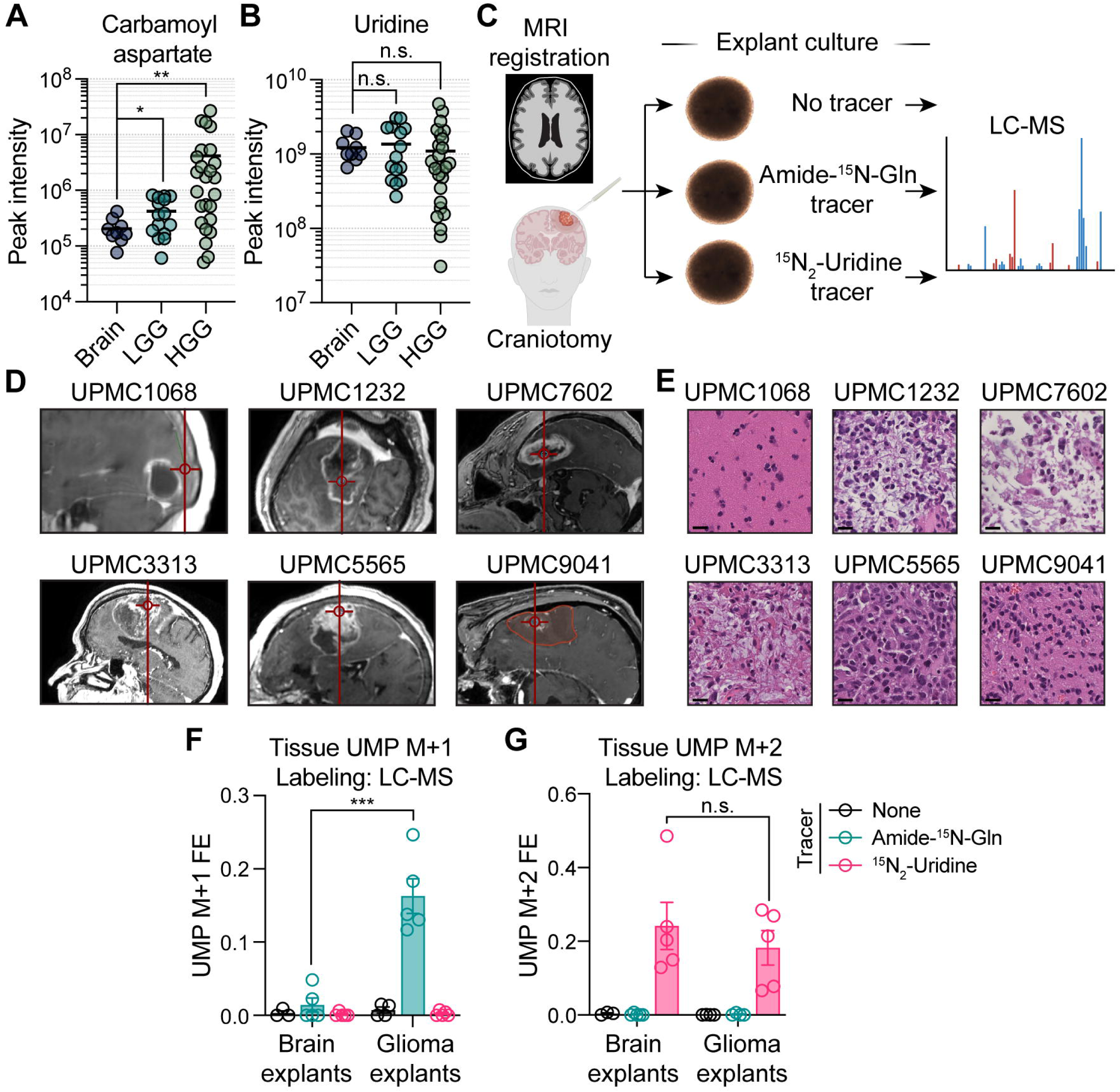
Human gliomas display constitutive de novo pyrimidine synthesis ex vivo. **(A-B)** Peak intensity of (A) carbamoyl aspartate and (B) uridine in normal brain (*n* = 9), index resections of lower-grade glioma (LGG, grades 2 and 3, *n* = 15), and index resections of high-grade glioma (HGG, grade 4, *n* = 26) from human surgical specimens. **(C)** Schema depicting stable isotope tracing in organoid explant cultures. **(D)** Representative intraoperative stereotactic neuronavigation of sample collection site from non-neoplastic neocortex (UPMC1068) or glioblastoma (all others). **(E)** H&E stains of tissues from (D). Scale bar = 20µm. **(F-G)** 18-hour stable isotope tracing of explants with amide-^15^N-glutamine (*n* = 5 per cohort), ^15^N_2_-uridine (*n* = 5 per cohort), or no tracer (*n* = 3 for brain cohort, *n* = 4 for glioma cohort). UMP (F) M+1 and (G) M+2 FE values are shown. n.s. = not significant, \**p* < 0.05, \*\**p* < 0.01, \*\*\**p* < 0.001. Two-tailed *p* values were determined by unpaired *t*-test. For all panels, data are means ± SEM. See also Figure S4.

Labeling in explants mirrored xenografts: glioma but not non-malignant brain explants demonstrated constitutive de novo pyrimidine synthesis whereas all explants displayed robust uridine salvage (Figures 4F and 4G). Like cell culture experiments (Figure S2L), glutamine tracer accumulation was higher in non-malignant brain than glioma explants, whereas uridine accumulation was similar (Figures S4A and S4B). These data suggest our assay may underestimate differences in de novo pathway activity between glioma and nonmalignant brain due to larger unlabeled glutamine pools in the former. Collectively, our results show that de novo pyrimidine synthesis pathway activation is a conserved feature of primitive cell populations in primary brain tissues.

### De novo pyrimidine synthesis is repressed in a uridine-dependent manner in differentiated cells

As we pursued mechanism(s) by which preference for pyrimidine salvage is established in differentiated cells, we considered a seemingly paradoxical finding. Although NHA cells show a strong preference for pyrimidine salvage (Figure 2L) and rapidly consume uridine relative to GSCs (Figure 5A), they do not display proliferation defects during routine culture when uridine is depleted. To ask how pyrimidine synthesis is regulated under standard culture conditions, we collected conditioned media and cells from NHA cultures during a 24-hour period in which medium was replenished in 6-hour intervals. We observed a reciprocal, cyclical relationship between intracellular carbamoyl aspartate and media uridine levels (Figure 5B, upper panel). As NHAs depleted media uridine, intracellular carbamoyl aspartate levels rose. Intracellular carbamoyl aspartate fell sharply, however, when media was replenished and uridine availability was restored. These metabolic shifts did not affect cell proliferation, as cell numbers steadily increased during this experiment (Figure 5B, lower panel). Pyrimidine synthesis pathway switching occurred without cyclical effects on UMP, UTP, or other metabolites downstream of de novo and salvage pathway convergence (Figures 5C and S5A-S5C). We therefore hypothesized that differentiated cells repress de novo pyrimidine synthesis in response to uridine availability.

**Figure 5.**
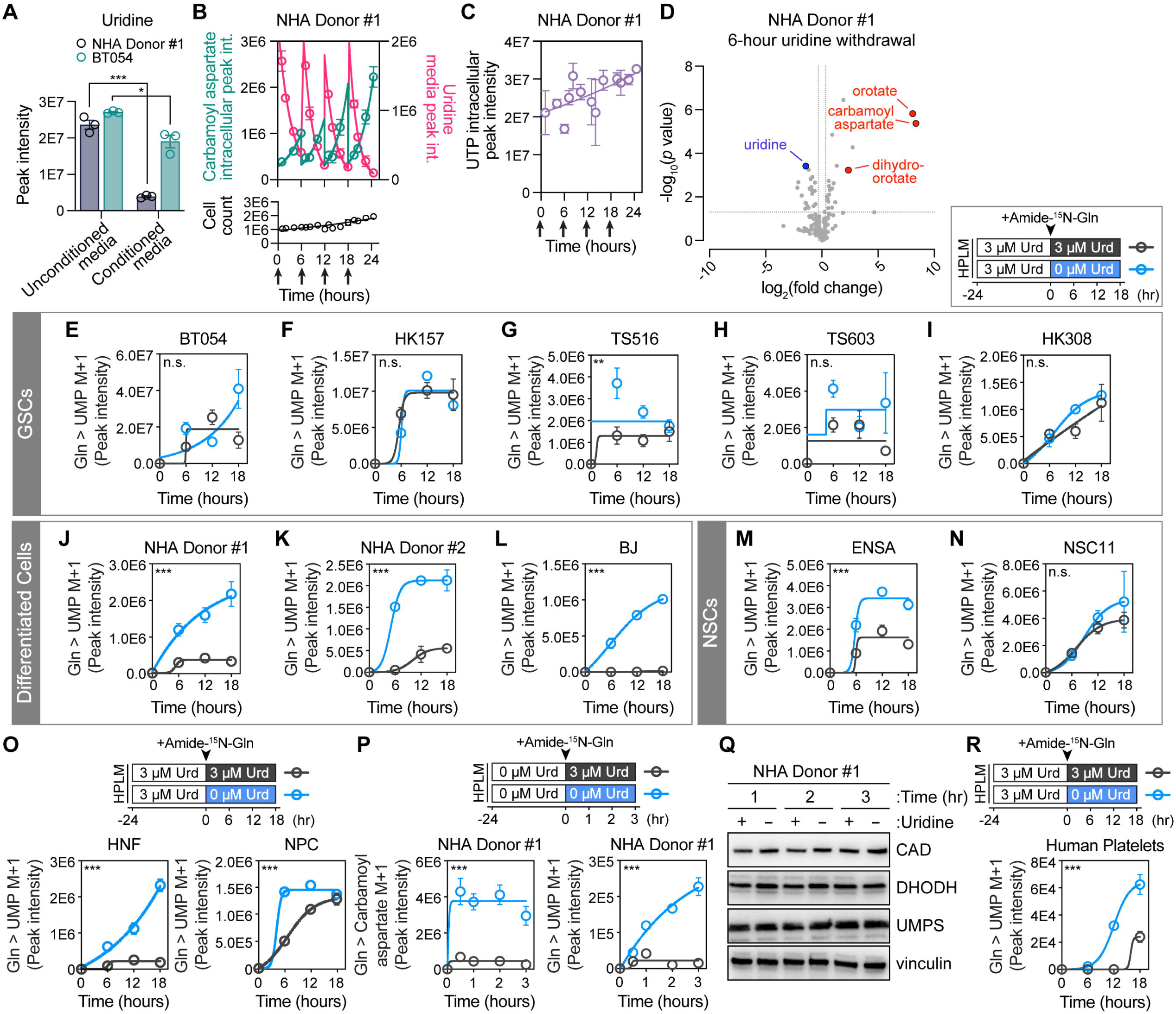
Uridine deprivation acutely activates de novo pyrimidine synthesis in differentiated cells through a post-transcriptional mechanism. **(A)** Peak intensity of uridine in unconditioned and conditioned media from 18-hour cultures of NHA Donor #1 or BT054 cells (*n* = 3 per group). **(B-C)** Quantification of (B, top panel) intracellular carbamoyl aspartate and media uridine, (B, bottom panel) cell count, and (C) intracellular UTP in NHA Donor #1 cells cultured in DMEM with 3µM uridine. Media changes indicated by arrows. Curves are fit by nonlinear regression for (B) a 6-hour window after each media change or (C) full 24-hour experiment (*n* = 3). Int. = intensity. **(D)** Volcano plot of metabolites in NHA Donor #1 cells cultured in 0µM or 3µM uridine for 6 hours after 24 hours of culture with 3µM uridine (*n* = 6 per group). **(E-N)** Amide-^15^N-glutamine stable isotope tracing in (E-I) GSC, (J-L) differentiated, or (M-N) NSC lines cultured in 0µM or 3µM uridine after 24 hours of culture with 3µM uridine (*n* = 3 for all except HK308, for which *n* = 2). Urd = uridine. **(O)** Amide-^15^N-glutamine stable isotope tracing in HNF or NPC cells cultured in 0µM or 3µM uridine after 24 hours of culture with 3µM uridine (*n* = 3 per group). **(P)** Amide-^15^N-glutamine stable isotope tracing in NHA Donor #1 cells cultured in 0µM or 3µM uridine after 24 hours of culture with 0µM uridine (*n* = 3 per group). **(Q)** Representative immunoblots for CAD, DHODH, and UMPS in NHA Donor #1 cells cultured in 0µM or 3µM uridine after 24 hours of culture with 3µM uridine. **(R)** Amide-^15^N-glutamine stable isotope tracing in human donor platelets cultured in 0µM or 3µM uridine after 24 hours of culture with 3µM uridine (*n* = 3). \**p* < 0.05, \*\**p* < 0.01, \*\*\**p* < 0.001 Two-tailed *p* values were determined by unpaired *t*-test. *t*-tests in (E-P and R) compare area under the curve values for 0µM uridine and 3µM uridine conditions. For all panels, data are means ± SEM. See also Figure S5.

To test this idea, we evaluated de novo pyrimidine synthesis pathway activity under uridine replete and depleted conditions. In NHAs deprived of uridine, de novo pyrimidine synthesis intermediates rose sharply in a time-dependent manner (Figures 5D and S5D). We next preconditioned cells in HPLM with 3 μM uridine for 24 hours before replacing media with one of two medium formulations: 1) HPLM with amide-^15^N-glutamine and 3 μM uridine, or 2) HPLM with amide-^15^N-glutamine and without uridine. We then assessed UMP labeling over 18 hours (Figures 5E-5N). Immortalized astrocytes and fibroblasts rapidly activated de novo pyrimidine synthesis upon uridine withdrawal (Figures 5J-5L). Conversely, de novo pyrimidine synthesis in most GSCs and NSCs was unaffected by uridine deprivation (Figures 5E-5I and 5M-5N). De novo pyrimidine synthesis did increase in TS516 GSCs and ENSA NSCs upon uridine deprivation, but these responses were more subtle than those of differentiated cell lines. We also evaluated isogenic fibroblasts and iPSC-derived NPCs in a similar assay (Figure 5O). Although both cultures activated de novo pyrimidine synthesis upon uridine withdrawal, differentiated fibroblasts responded more robustly than primitive NPCs. Taken together, our data show that uridine availability represses de novo pyrimidine synthesis in differentiated, but not primitive, cells (Figures S5E and S5F).

We sought to characterize the mechanistic link between uridine availability and de novo pyrimidine synthesis activity. We first considered whether low expression of de novo pyrimidine synthesis enzymes in differentiated cells may restrict pathway activity, as prior work has implicated expression as a driver of de novo pyrimidine synthesis in tumor cells^35^. Indeed, protein levels of the three enzymes that comprise the de novo synthesis pathway (CAD, DHODH, and UMPS) are higher in human GBMs than non-malignant brain^47^ (Figures S5G-S5I). However, the salvage pathway enzyme UCK2 (one of two kinases that phosphorylate uridine to generate UMP) was also upregulated in GBMs (Figures S5J-S5L). We then evaluated RNA sequencing data^28,46^ from our GSCs and NHAs and found that NHAs generally did not express de novo pathway enzymes at lower levels nor salvage pathway enzymes at higher levels than GSCs (Figure S5M). Therefore, basal differences in gene expression cannot solely explain de novo pyrimidine synthesis repression in differentiated cells under nutrient rich conditions.

We next evaluated kinetics of the uridine deprivation response in NHA cells at earlier timepoints (Figure S5N). Increases in carbamoyl aspartate labeling from amide-^15^N-glutamine were evident 1 hour following uridine depletion, with stark differences at 3 hours. Likewise, uridine replenishment rapidly inhibited de novo synthesis in uridine-starved differentiated cells (Figure 5P). Amide-^15^N-glutamine labeling of both carbamoyl aspartate and UMP was suppressed 30 minutes after uridine replenishment. Given these rapid kinetics, we suspected involvement of a post-transcriptional mechanism. We therefore evaluated levels of de novo pathway enzymes in NHA cells and BJ fibroblasts following uridine withdrawal. After validating specificity of CAD, DHODH, and UMPS antibodies (Figure S5O), we subsequently found no changes in these proteins following short- (Figure 5Q) or long-term (Figures S5P-S5R) uridine starvation. We next assessed cultured human platelets, which lack nuclei and do not display transcription^48^ but do harbor de novo pyrimidine synthesis enzymes^49^. Platelets activated de novo pyrimidine synthesis after uridine withdrawal (Figure 5R), indicative of a post-transcriptional mechanism.

We considered whether changes in intracellular metabolite levels stimulate de novo pyrimidine synthesis upon uridine withdrawal. The only metabolites consistently altered in NHA cells following uridine withdrawal (Figure S5D) included uridine and de novo pyrimidine synthesis intermediates. De novo pathway substrate levels were not altered upon uridine withdrawal. We also evaluated metabolites involved in allosteric control of pyrimidine synthesis: UTP, which represses CAD^50–52^, and uric acid, which inhibits UMPS^12^. At timepoints when uridine deprivation stimulated de novo pyrimidine synthesis (Figures S5S and S5V) we observed no differences in UTP (Figures S5T and S5W) or uric acid (Figures S5U and S5X).

Activating phosphorylation of CAD on Ser 1859 downstream of the insulin/PI3K/mTORC1 signaling axis is an important post-translational mechanism of pyrimidine synthesis regulation^53,54^. To ask if uridine availability changes this modification, we first validated antibodies against S6 phospho-S240, S6K phospho-T389 (both downstream targets of mTORC1 signaling^53,54^) and CAD phospho-S1859 (Figure S5Y). Neither NHAs nor BJ fibroblasts increased mTORC1 signaling markers during short- or long-term uridine starvation (Figures S5Z-S5CC). In the same conditions, CAD phospho-S1859 levels did not change (Figures S5DD-S5GG), suggesting that uridine governs de novo pyrimidine synthesis activity in differentiated cells independently of mTORC1. Therefore, uridine-dependent repression of de novo pyrimidine synthesis in differentiated cells is mediated by a post-transcriptional process that is independent of established regulatory mechanisms.

### Constitutive de novo pyrimidine synthesis in primitive cells is associated with CAD phosphorylation at Ser 1900

We next asked a related question: by what mechanism do primitive cells uncouple de novo pyrimidine synthesis activity from uridine availability? Because expression of pyrimidine metabolic genes was not markedly different between differentiated and primitive cells (Figure S5M), we reasoned that constitutive de novo pyrimidine synthesis in the latter may be driven by post-translational modification. Therefore, we assessed proteomic and phosphoproteomic analyses of human GBM and nonmalignant brain tissues^47^. We first used linear regression to evaluate correlations between the primitive cell marker SOX2^41^ and phosphorylation marks on pyrimidine synthesis enzymes in GBM tissues (Figure 6A). Two modifications positively correlated with SOX2 in these tissues: phosphorylation of S61 on UPP1 and S1900 on CAD. We also found similar correlations with the primitive neural cell marker, OLIG2^55^ (Figure 6B). CAD S1900 is located near S1859, an established target of activating phosphorylation downstream of mTORC1 signaling^53,54^. To understand how CAD phosphorylation at S1900 may impact enzyme function, we used AlphaFold^56^ to evaluate the S1900 residue (Figure 6C). S1900 and S1859 are both predicted to reside on a large, unstructured, interdomain loop. Although CAD S1900 phosphorylation has been captured in prior proteomic profiling studies^53,57–61^ and increases following Kaposi’s sarcoma-associated herpesvirus (KSHV) infection of human cells^62^, its endogenous regulatory mechanisms and functional role in uninfected cells are not clear.

**Figure 6.**
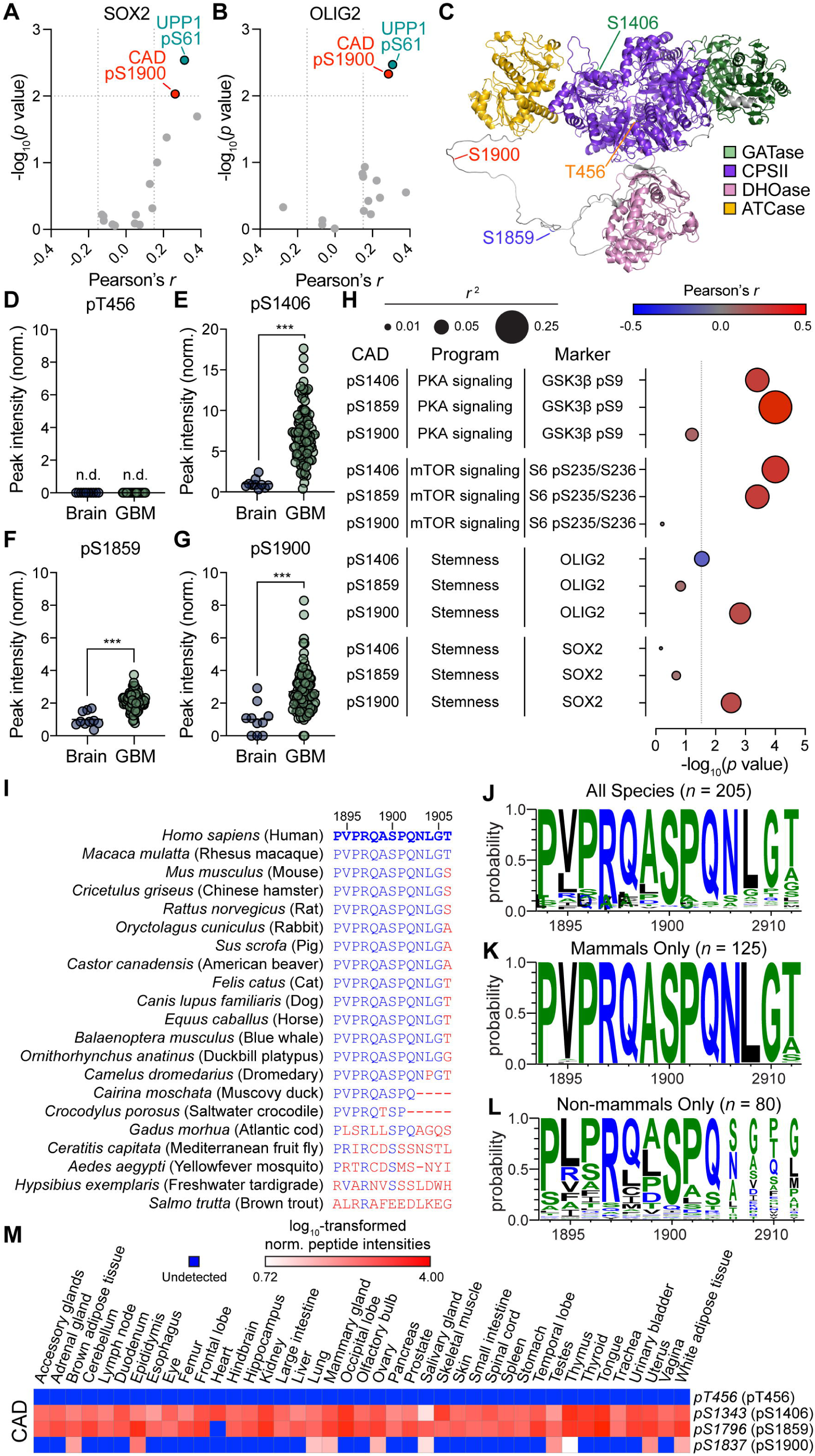
CAD S1900 phosphorylation is enriched in primitive cell populations. **(A-B)** Volcano plot of Pearson’s *r* and *p* values for correlations between phosphorylation modifications of pyrimidine synthesis enzymes and (A) SOX2 or (B) OLIG2 protein expression in human glioblastomas (GBMs) (*n* = 99). **(C)** AlphaFold predicted structure of CAD. GATase = glutamine amidotransferase. CPSII = carbamoyl phosphate synthetase II. ATCase = aspartate carbamoyltransferase. DHOase = dihydroorotase. **(D-G)** Relative abundance of (D) T456, (E) S1406, (F) S1859, and (G) S1900 phosphorylation modifications of CAD in human brain and GBM. Phosphorylation site abundance is normalized to total CAD protein and expressed relative to brain. n.d. = not detected. (*n* = 10 brain, *n* = 99 GBM). **(H)** Correlation between CAD phosphorylation sites and signaling program or stemness markers in human GBM. Phosphorylation site abundance is normalized to total protein. (*n* = 99). Panels (A-B and D-H) are reanalysis of raw data published in Wang et al., 2021^47^. **(I-L)** Sequence alignment relative to human CAD S1900. (I) Sequences from selected species. (J-L) Sequence logos of (J) all species, (K) mammals, or (L) non-mammals. Amino acids in blue are hydrophilic, green are neutral, and black are hydrophobic. **(M)** Heatmaps depicting relative phosphorylation of residues analogous to T456, S1406, S1859, and S1900 sites on human CAD protein in mouse tissues. Data are log_10_-transformed peptide intensities normalized to total CAD protein. Intensities are reanalysis of data published in Giansanti et al., 2022^85^. \*\*\**p* < 0.001. Two-tailed *p* values were determined by (A-B, H) linear regression analysis or (D-G) unpaired *t*-test. For all panels, data are means ± SEM. See also Figure S6.

We next assessed phosphorylation of S1900 and other sites on CAD with established regulatory functions (T456, S1406, and S1859) in human GBM and non-malignant brain. We validated each peptide by examination of MS/MS spectra (Figures S6A-S6C). GBMs displayed elevated CAD phosphorylation at S1406, S1859, and S1900 relative to non-malignant brain, whereas T456 phosphorylation was undetectable in both tissues (Figures 6D-6G). These data revealed a correlation between primitive cell composition (Figures 6A and 6B), CAD S1900 phosphorylation (Figure 6G), and basal rates of de novo pyrimidine synthesis (Figure 4F) in human brain tissues. As specificity controls, we benchmarked relationships between signaling or primitive neural markers and CAD phosphorylation (Figure 6H and S6E-S6V). As expected, CAD S1406 and S1859 phosphopeptide levels correlated with markers of PKA (phosphorylation of GSK3β at S9^63,64^; Figures 6H, S6E, and S6K) and mTORC1^53,54^ (phosphorylation of ribosomal protein S6 at S235/S236; Figures 6H, S6F, and S6L) signaling, respectively. Despite the role of MAPK signaling in modulation of CAD T456 (but not S1406 or S1859) phosphorylation^65^, both S1406 and S1859 phosphopeptide levels positively correlated with ERK phosphorylation^66^ (Figures S6I-S6J and S6O-S6P). These findings suggest either crosstalk or correlation between PKA, mTORC1, and MAPK signaling^67^ in GBM. CAD S1900 phosphorylation, however, positively correlated with both SOX2 and OLIG2 (Figures 6H and S6S-S6T) but not with markers of PKA, mTORC1, or MAPK signaling (Figures 6H, S6Q-S6R, and S6U-S6V). Moreover, CAD phosphorylation at S1406 and S1859 did not positively associate with primitive cell markers (Figures 6H, S6G-S6H, and S6M-S6N). These data suggest two independent regulatory axes converge on CAD in GBM: growth factor signaling-dependent phosphorylation of CAD at S1406 and S1859 and cell state-specific phosphorylation of CAD at S1900.

To assess whether CAD phosphorylation on S1900 is evolutionarily conserved, we evaluated CAD amino acid sequences across all trifunctional CAD orthologs in UniProt^68^. S1900 and surrounding amino acid residues are strongly conserved in CAD across mammalian species, with weaker conservation in non-mammals (Figures 6I-6L). Therefore, CAD S1900 phosphorylation may constitute a molecular link between cell state and pyrimidine synthesis activity that has persisted through mammalian evolution.

To assess the relevance of CAD phosphorylation to physiology, we assessed abundance of T456, S1406, S1859, and S1900 Cad phosphopeptides in adult mouse tissues. Phosphorylation of T456 (mediated by MAPK signaling) was not detected in any tissue, whereas S1406 and S1859 phosphorylation (mediated by PKA and mTORC1 signaling, respectively) was observed in nearly all tissues (Figure 6M). In contrast, Cad S1900 displayed a tissue-specific pattern of phosphorylation. Sex organs (including testes, epididymis, and ovary) were overrepresented among tissues displaying Cad S1900 phosphorylation, suggesting a link with undifferentiated gametes. Although Cad phosphorylation patterns differed significantly across tissue types and residues, expression of pyrimidine synthetic enzymes were largely similar across tissues (Figure S6D).

### CAD phosphorylation at Ser 1900 is sufficient to overcome uridine-dependent repression of de novo pyrimidine synthesis

Because S1859 phosphorylation by mTORC1 signaling increases CAD activity^53,54^, we hypothesized that phosphorylation of the interdomain loop it shares with S1900 may augment CAD catalytic function and contribute to de novo pyrimidine synthesis activation in primitive cells. Therefore, we asked if CAD S1900 phosphorylation is sufficient to overcome the inhibitory effect of uridine availability on de novo pyrimidine synthesis in differentiated cells (Figures 5J-5L). We used G9C, a line derived from Chinese hamster ovary (CHO) cells that lacks endogenous CAD expression^69^, and NHA cells expressing the following constructs: WT CAD, a CAD S1859 phosphomimetic mutant (S1859D), a CAD S1900 phosphomimetic mutant (S1900D), a CAD mutant that is unable to be phosphorylated at either S1859 or S1900 residues on this interdomain loop (S1859A/S1900A), or an empty vector (EV) (Figures S7A and S7B). The CAD S1859D mutant served as an internal positive control to model phosphorylation-dependent activation of CAD. Resulting stable lines were cultured under physiologic uridine and exposed to amide-^15^N-glutamine to assess de novo pyrimidine synthesis. In parallel, WT CAD- or EV-expressing cells were cultured with amide-^15^N-glutamine under uridine-free conditions to benchmark changes in de novo pyrimidine synthesis evoked by uridine deprivation. Under uridine replete conditions, cells expressing WT CAD increased labeling of both carbamoyl aspartate and UMP by amide-^15^N-glutamine relative to EV-expressing cells (Figures 7A-7D). CAD S1900D phosphomimetic-expressing cells displayed similar or higher levels of de novo pyrimidine synthesis relative to WT CAD-expressing cells deprived of uridine. Stimulation of de novo pyrimidine synthesis by CAD S1900D recapitulated effects of the CAD S1859D phosphomimetic, underscoring the role of interdomain loop phosphorylation. Conversely, cells expressing the CAD S1859A/S1900A phosphodeficient mutant displayed similar or lower levels of de novo pyrimidine synthesis compared with WT CAD-expressing cells. To complement tracing, we performed metabolomics analysis of NHA cells engineered to express WT CAD, CAD S1900D, or an EV control. Although ectopically expressing WT CAD or CAD S1900D in NHA cells led to accumulation of carbamoyl aspartate, only CAD S1900D expression increased global pyrimidine pools (Figures 7E and 7F). These data show that CAD S1900 phosphorylation is sufficient to overcome the strict preference for pyrimidine salvage displayed by differentiated cells. Moreover, our findings implicate CAD S1900 phosphorylation as a mechanism contributing to constitutive activation of de novo pyrimidine synthesis in primitive cells.

**Figure 7.**
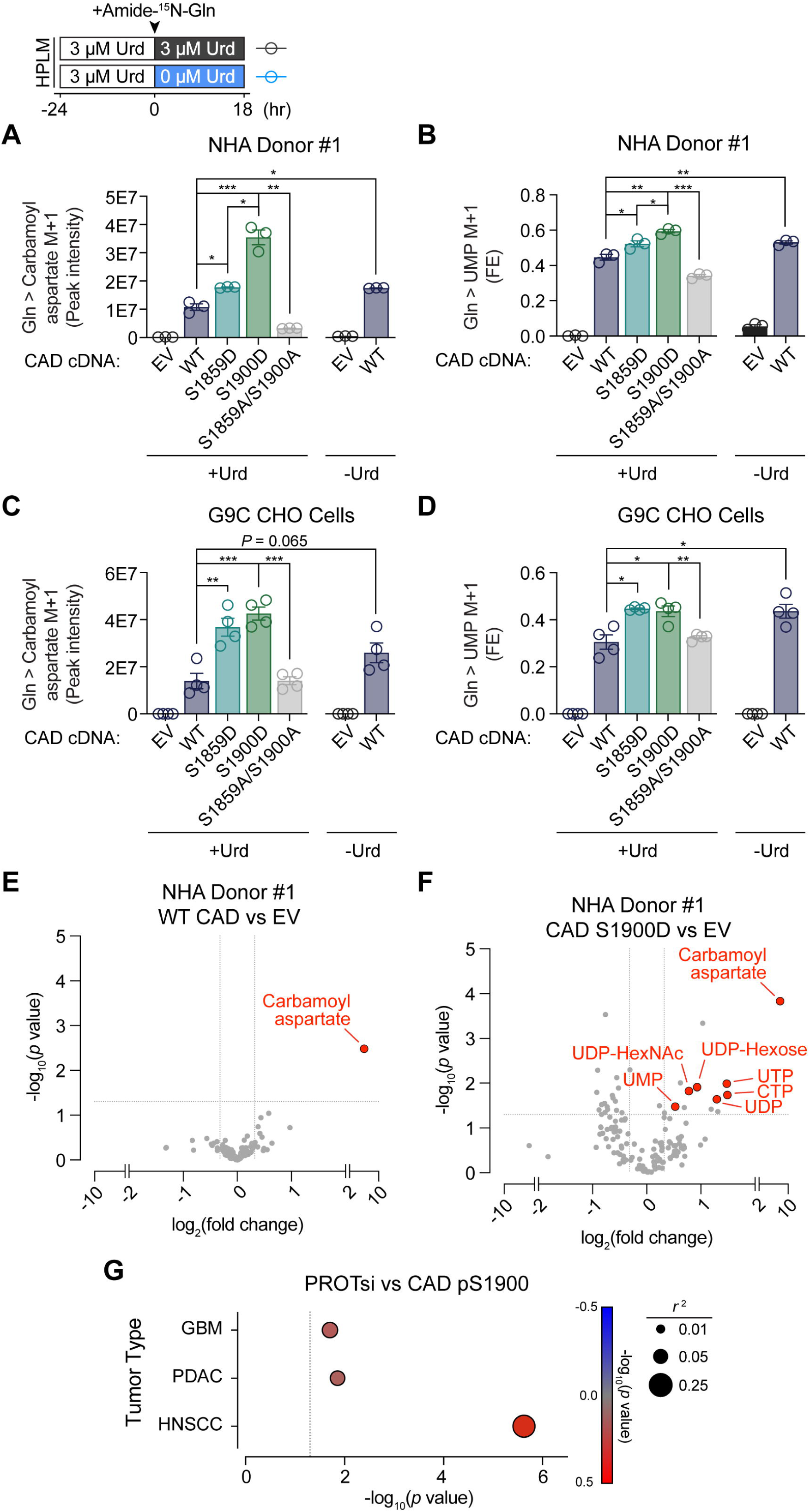
Mimicking CAD S1900 phosphorylation activates de novo pyrimidine synthesis in differentiated cells. **(A-D)** Amide-^15^N-glutamine stable isotope tracing for 18 hours in 0µM or 3µM uridine in (A and B) NHA Donor #1 and (C and D) G9C CHO cells expressing either empty vector (EV), wild-type (WT) CAD, or CAD mutants. Data depict (A and C) carbamoyl aspartate M+1 peak intensity and (B and D) UMP M+1 FE (*n* = 3 per group for NHA Donor #1 cells and *n* = 4 per group for G9C CHO cells). **(E-F)** Volcano plot of metabolites in NHA Donor #1 cells expressing (E) WT CAD versus EV or (F) S1900D CAD versus EV in 3µM uridine. UDP-HexNAc = uridine diphosphate-N-acetylhexosamine. CTP = cytidine triphosphate. (*n* = 3 per group). **(G)** Correlation between protein expression-based stemness index (PROTsi) and CAD S1900 phosphorylation in human GBM, pancreatic ductal adenocarcinoma (PDAC), and head and neck squamous cell carcinoma (HNSCC) issues. Data are from reanalysis of proteomics dataset published in Kołodziejczak-Guglas and Simões et al., 2025^70^. \**p* < 0.05, \*\**p* < 0.01, \*\*\**p* < 0.001. Two-tailed *p* values were determined by unpaired *t*-test. For all panels, data are means ± SEM. See also Figure S7.

Upon establishing CAD S1900 phosphorylation as a determinant of pyrimidine synthesis pathway choice, we asked if the cancer relevance of this modification extends beyond GBM. We assessed a study reporting a proteomic-based stemness index (PROTsi) to quantify primitive cell features in a panel of human tumors^70^. CAD S1900 phosphopeptide was detected in GBM, pancreatic ductal adenocarcinoma (PDAC), and head and neck squamous cell carcinoma (HNSCC). PROTsi positively correlated with CAD S1900 phosphorylation in all three tumor types (Figure 7G), indicating that association between this modification and primitive cell states is generalizable across human tumor subtypes.

## DISCUSSION

Nitrogen metabolic pathways operate in parallel with those of carbon metabolism^10^. Amino acids, nucleotide precursors, ions, and other nitrogenous small molecules serve as substrates for enzymes that sustain macromolecule synthesis, energy production, stress resistance, and cell growth. System level understanding of how nitrogen metabolism pathways operate and intersect with one another in human cells, however, has been limited by a lack of appropriate tools. Although systems biology approaches have revealed global nitrogen metabolism programs in bacteria^71,72^, these studies exploit unique bacterial dependence on ammonium for nitrogen assimilation and are difficult to apply to human cells that use a wide range of nitrogenous nutrients. Here, we develop, validate, and apply an experimental platform that addresses complexities of studying nitrogen metabolism in cells from higher organisms. This platform generates both high-level representations of nitrogen metabolism programs and quantitative measurements of differences in substrate utilization between paired cultures. To facilitate adoption of this platform, we describe how to prepare the HPLM library and provide access to a computational pipeline to aid in processing the rich metabolite labeling datasets generated by the platform.

We used the platform ask how nitrogen metabolism differs between differentiated, non-transformed astrocytes and primitive, malignant glioma cells. One key difference involved pyrimidine synthesis substrate utilization – glioma cells preferentially used glutamine to fuel de novo pyrimidine synthesis whereas immortalized astrocytes nearly exclusively engaged pyrimidine salvage from uridine. Activation of de novo pyrimidine synthesis has been reported in many cancer contexts^14,31,36,39,40,73–75^, but the underlying mechanisms are not fully understood. Data from our platform suggest that upregulation of de novo pyrimidine synthesis in undifferentiated tumor cells is cell-intrinsic, independent of microenvironmental nutrient limitation, and similar to primitive, non-transformed cells. Our data complement prior studies demonstrating increased de novo pyrimidine synthesis enzyme levels as a contributor to activation of this pathway in tumors^35^. Indeed, several oncogenes and tumor suppressors directly regulate transcription of genes encoding pyrimidine synthesis enzymes, as exemplified by c-myc-dependent transactivation of *CAD*^76^. Our findings imply, however, that enzyme expression does not solely explain differences between differentiated and primitive cells. Under basal conditions, non-malignant, adult brain tissues and cultured astrocytes and fibroblasts repress de novo pyrimidine synthesis yet constitutively express pathway enzymes. These findings inform a model in which de novo pyrimidine synthesis machinery is poised for activation in differentiated cells but requires molecular cues to initialize. This regulatory framework may provide an advantage to cells during acute nutrient deprivation by allowing them to engage de novo pyrimidine synthesis on a shorter timescale than would be required to engage transcription and translation.

Our work reveals uridine depletion as an important cue for differentiated cells to activate de novo pyrimidine synthesis. We uncovered a reciprocal relationship between de novo pyrimidine synthesis and extracellular uridine availability in differentiated cell cultures. These data suggest that differentiated cells sense changes in uridine abundance to promote nucleotide homeostasis during nutrient deprivation. Uridine restriction stimulates de novo pyrimidine synthesis through a post-transcriptional process distinct from established regulatory mechanisms. The molecular constituents involved in this pathway are not known but may be characterized through future studies. Nevertheless, our findings demonstrate that cells monitor and respond to changes in pyrimidine abundance in a manner that may cooperate with purine sensing mechanisms^77,78^ to promote adaptation to nucleotide depletion and metabolic stress.

Allosteric regulation and post-translational modification of de novo pyrimidine synthesis enzymes play important roles in determining pathway flux. For example, UTP and uric acid directly bind and inhibit CAD^50–52^ and UMPS^12^, respectively. Negative feedback on CAD by UTP balances rates of pyrimidine metabolism throughout the cell cycle^51^. Signaling from MAPK^65,79^, PKA^64,79^, and mTORC1^53,54^ pathways converge on CAD, inducing activating phosphorylation events at T456, S1406, and S1859, respectively. mTORC1-dependent CAD phosphorylation responds to insulin stimulation and links systemic nutrient availability with cellular nucleotide metabolism^53,54^. Results from our study deepen understanding of the signaling processes that govern de novo pyrimidine synthesis. We propose that differentiation state-dependent phosphorylation of CAD at S1900 represents a key mechanism of de novo pyrimidine synthesis regulation that operates in parallel with inputs from growth factor signaling pathways.

We identified S1900 phosphorylation of CAD based on its enrichment in primitive cell populations and show that this modification is sufficient to overcome uridine-dependent repression of de novo pyrimidine synthesis in differentiated cells. CAD S1900 phosphorylation was reported in one study of insulin-dependent CAD S1859 phosphorylation, although it was not regulated by insulin^53^. CAD S1900 phosphorylation was recently shown to contribute to de novo pyrimidine synthesis activation in KSHV-infected cells^62^, but the relevance of this modification outside the context of viral infection is not clear. We found that human GBM displayed higher CAD S1900 phosphorylation relative to non-malignant brain, and these levels positively correlated with markers of primitive cell populations within GBMs. Supporting the functional relevance of this observation, expressing a CAD S1900D phosphomimetic mutant in differentiated cells phenocopied basal de novo pyrimidine synthesis activation in primitive cells. These findings imply that activation of de novo pyrimidine synthesis in primitive cells can be at least partly explained by constitutive CAD S1900 phosphorylation. Structural predictions demonstrate that S1900 resides nearby S1859 on a large, unstructured, interdomain loop. We propose that signaling from mTORC1 and cell state-specific networks coalesce to regulate CAD activity through phosphorylation of this interdomain loop. Additional work is required to understand how these phosphorylation events may act in concert to control flux through CAD, but prior research indicates that oligomerization-dependent substrate channeling may be relevant^54^.

Selective CAD S1900 phosphorylation and activation of de novo pyrimidine synthesis in discrete cell populations may contribute to therapeutic indices of drugs targeting this pathway. These drugs include dihydroorotate dehydrogenase (DHODH) inhibitors such as teriflunomide and leflunomide, which are approved by the FDA to treat multiple sclerosis and rheumatoid arthritis, respectively^80^. Agents targeting DHODH are being investigated to treat various cancers and autoimmune disorders^34,81–84^. In these studies, CAD S1900 phosphorylation may represent a useful biomarker of basal differences in de novo pyrimidine synthesis activity in healthy and diseased tissues.

## ACKNOWLEDGEMENTS

The authors thank McBrayer, Abdullah, and DeBerardinis laboratory members, J. Engel, C. Menezes, W. Chen, and G. Hoxhaj for helpful discussions; D. Cahill (MGH), H. Kornblum (UCLA), I. Mellinghoff (MSKCC), R. Pieper (UCSF), R. Possemato (NYU), J. Rich (UNC), and S. Weiss (University of Calgary) for sharing cell lines; the UTSW Histo Pathology Core for assistance with H&E staining; the UTSW Animal Resource Center for assistance with mouse studies; and the UTSW Proteomics Core Facility for assistance collecting and interpreting proteomics data.

This study was supported by National Institutes of Health (NIH) grants R01CA258586 and R01CA289260 to S.K.M. and K.G.A., P50CA165962 and U19CA264504 to S.K.M., R35CA220449 to R.J.D., F32HL174084 to T.S.T., K99CA277576 to Y.X., and F30CA271634 to M.R.S. This work was also supported by awards from Oligo Nation to S.K.M. and K.G.A. and by Cancer Prevention and Research Institute of Texas (CPRIT) grants RR190034, RP230344, and RP2400489 to S.K.M. T.P.M., L.G.Z., and the Children’s Research Institute Metabolomics facility are supported by CPRIT Core Facilities Support Award RP240494. T.S.T was supported by an American Heart Association Postdoctoral fellowship 24POST1200352. Y.X. was supported by Human Frontier Science Program postdoctoral fellowship LT0018/2022-L.

The authors wish to extend heartfelt gratitude to the patients who generously donated samples, without whom this study would not have been possible.

## AUTHOR CONTRIBUTIONS

Conceptualization: MRS, KGA, SKM; Data curation: MRS, BCS, YX, GB, SSO, NM, SAS, MSR, WHH, SA; Formal Analysis: MRS, WG, GB, AL, SKM; Funding acquisition: MRS, KGA, SKM; Investigation: MRS, BCS, WG, YX, GB, BL, SSO, NM, LGZ, VTP, SAS, MSR, MML, CKE, WHH, SA, TS, MSMS, RW, JSP, XJ, TST; Methodology: MRS, BCS, WG, YX, SSO, NM, VTP, MML, DDS, POZ, AS, KGA; Project administration: MRS, SSO, NM, KGA, SKM; Resources: MRS, BCS, WG, YX, SSO, NM, VTP, MML, DDS, POZ, AS, KGA; Software: MRS, WG; Supervision: AS, TPM, NYA, RJD, KGA, SKM; Validation: AL, TER; Visualization: MRS, BCS, GB, SAS, MSR, SKM; Writing – original draft: MRS, SKM; Writing – review & editing: MRS, DDS, RJD, KGA, SKM, TER.

## STAR METHODS

### Key Resources Table

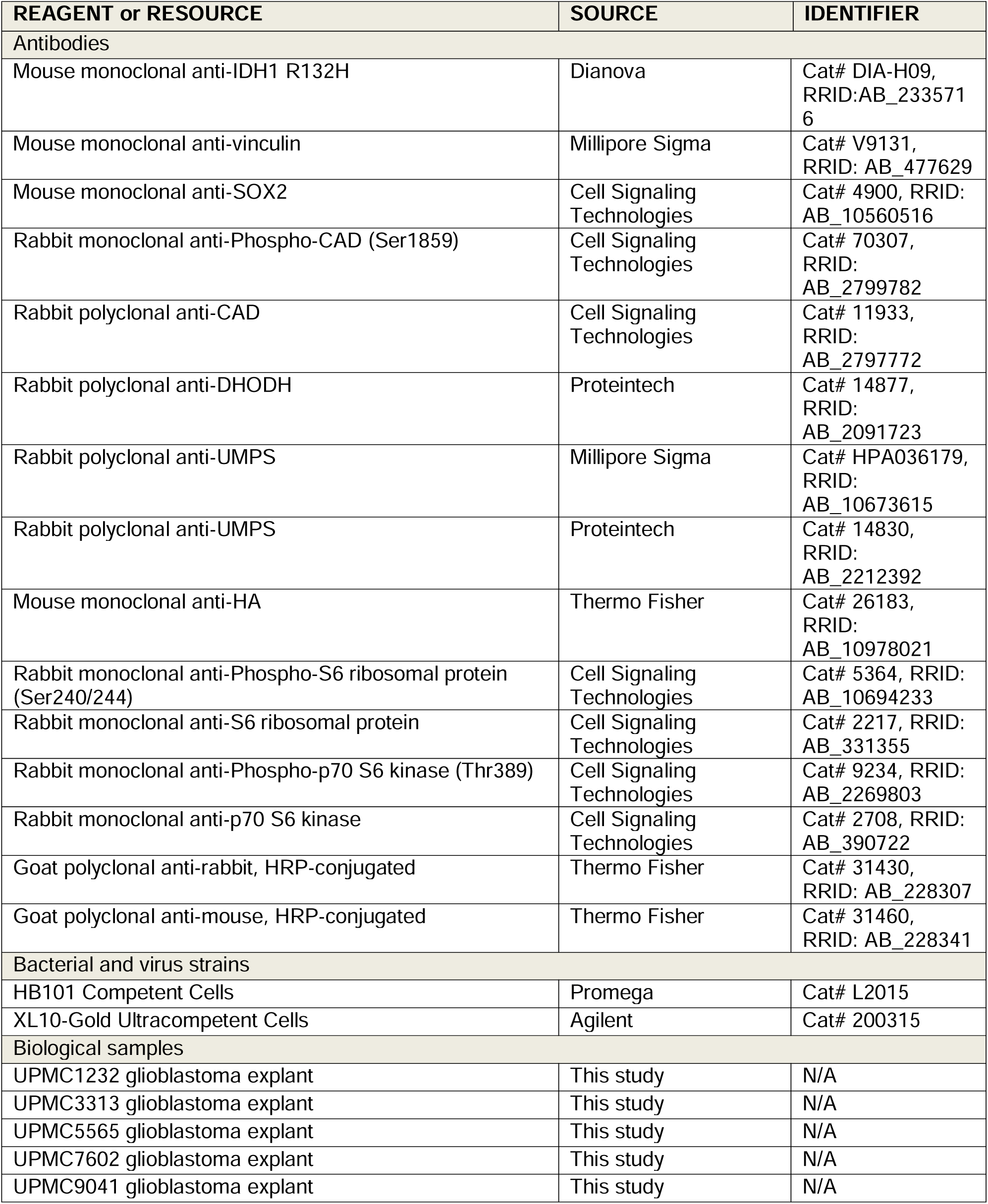
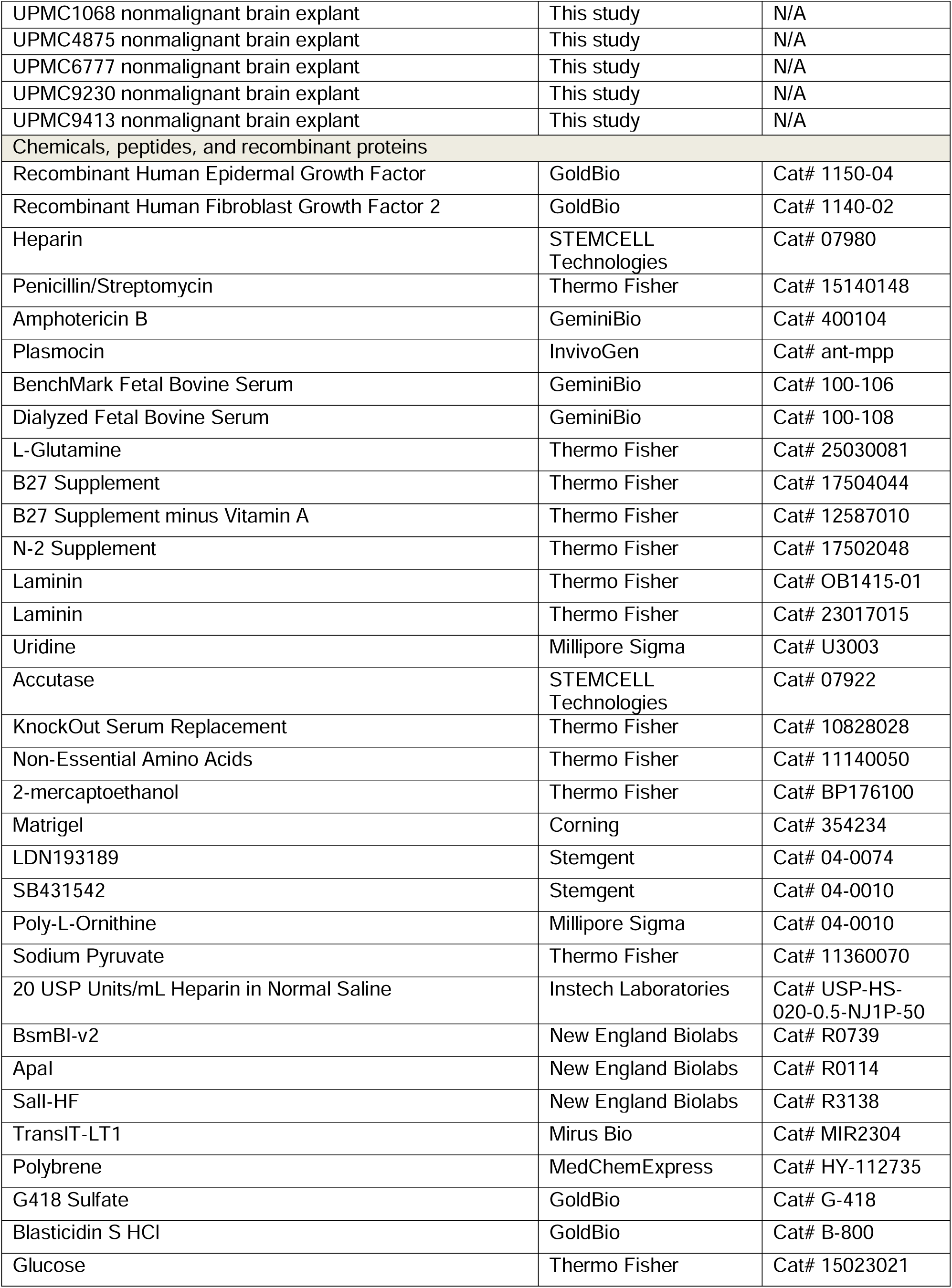
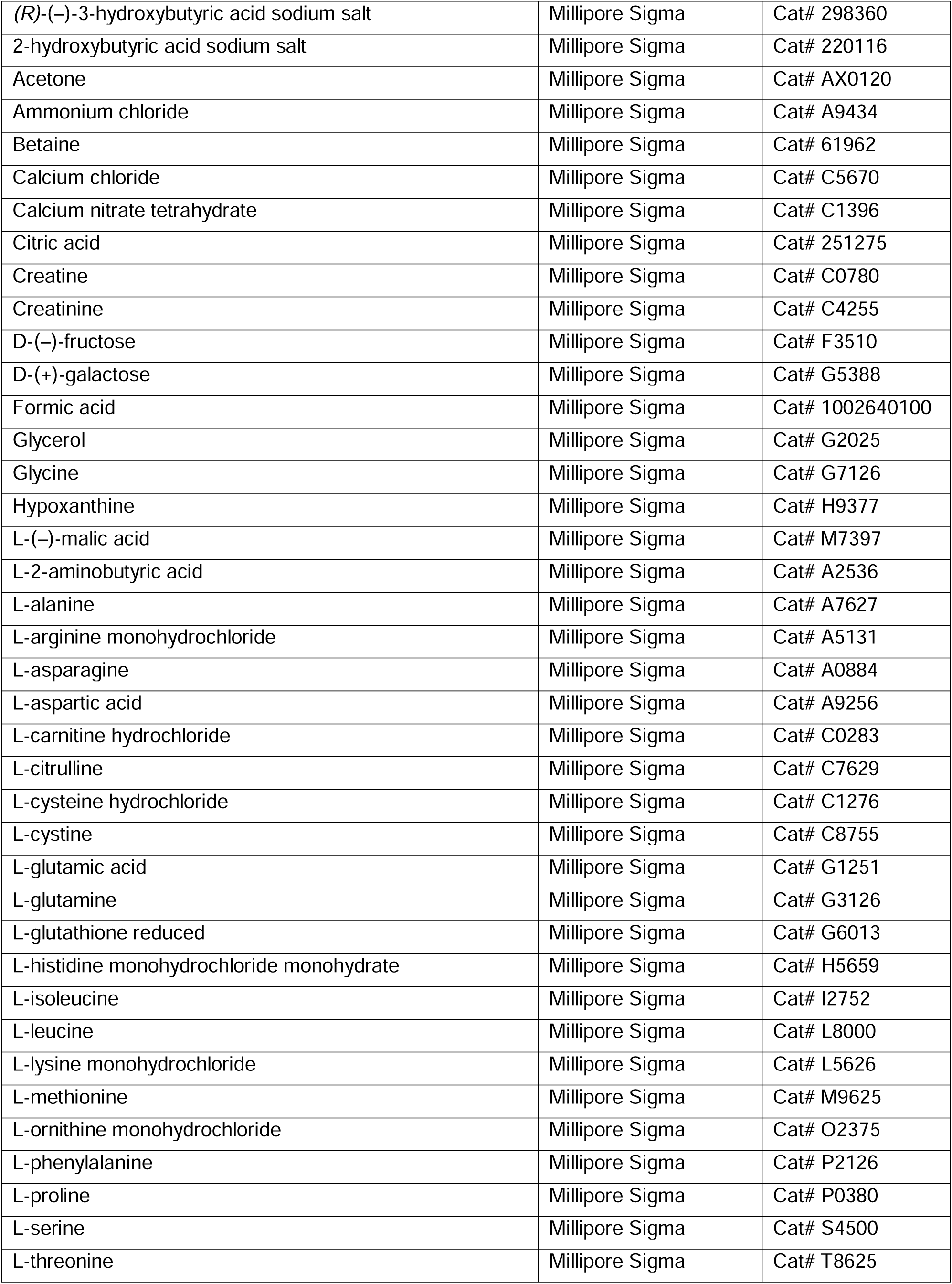
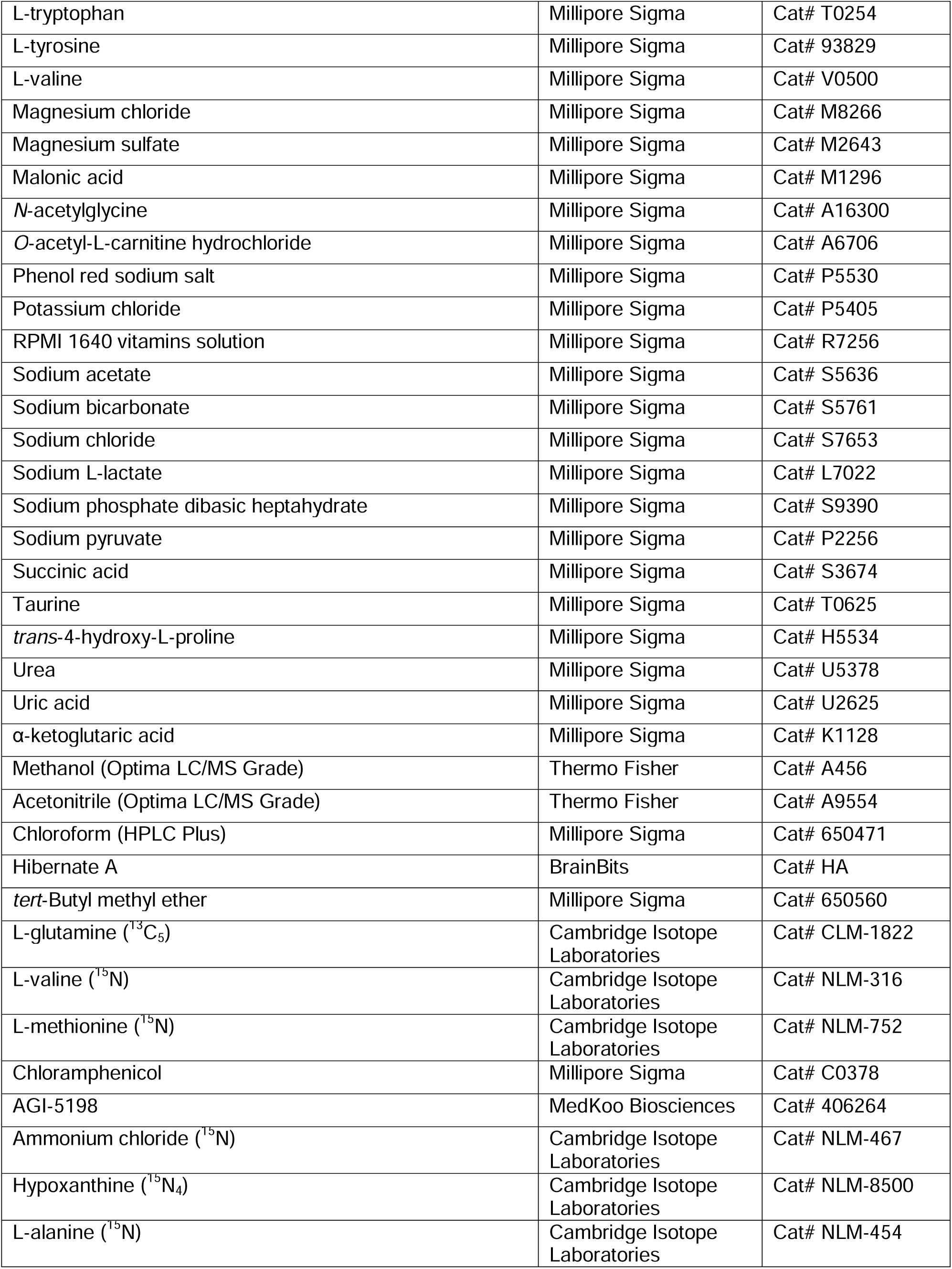
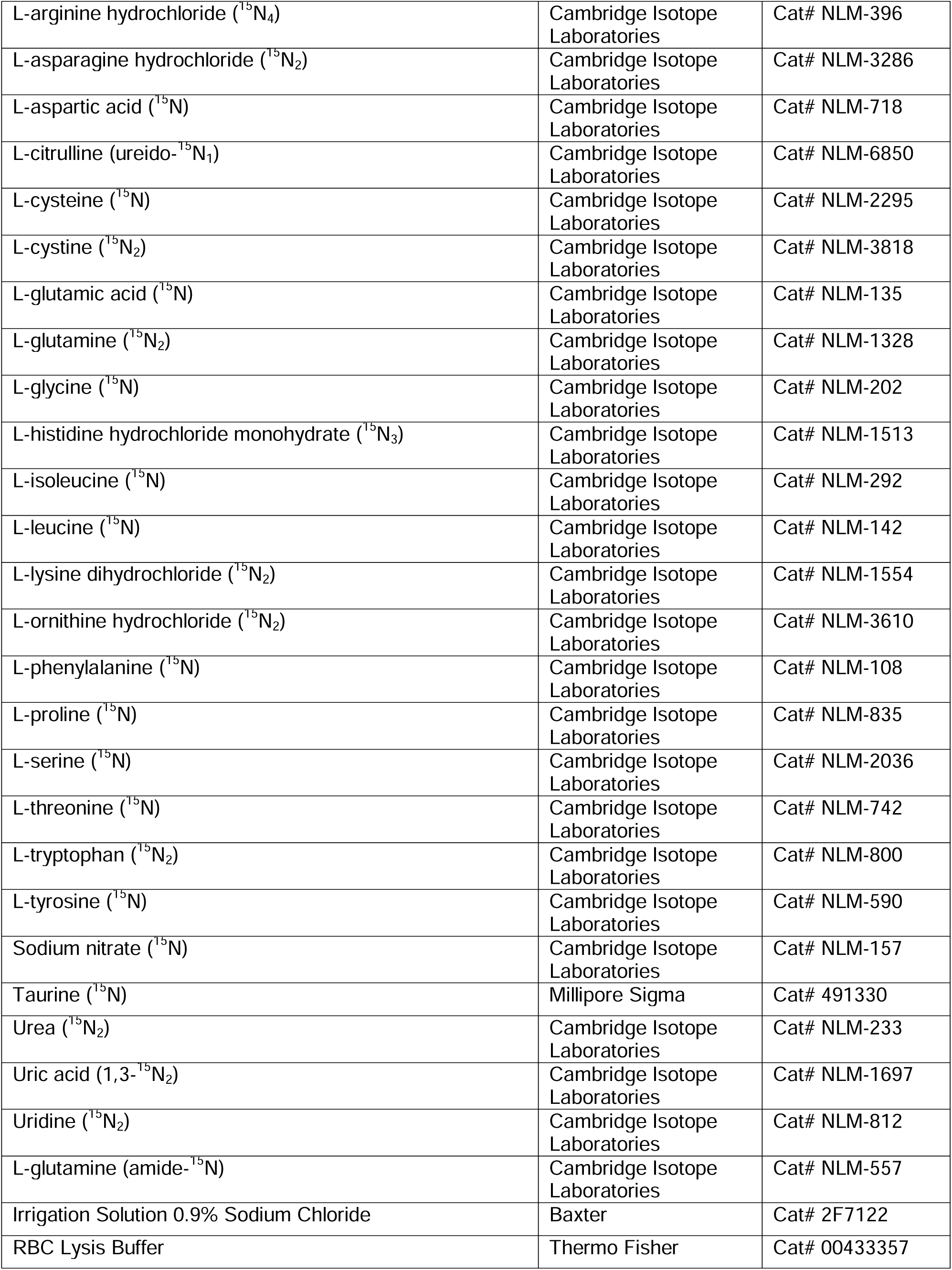
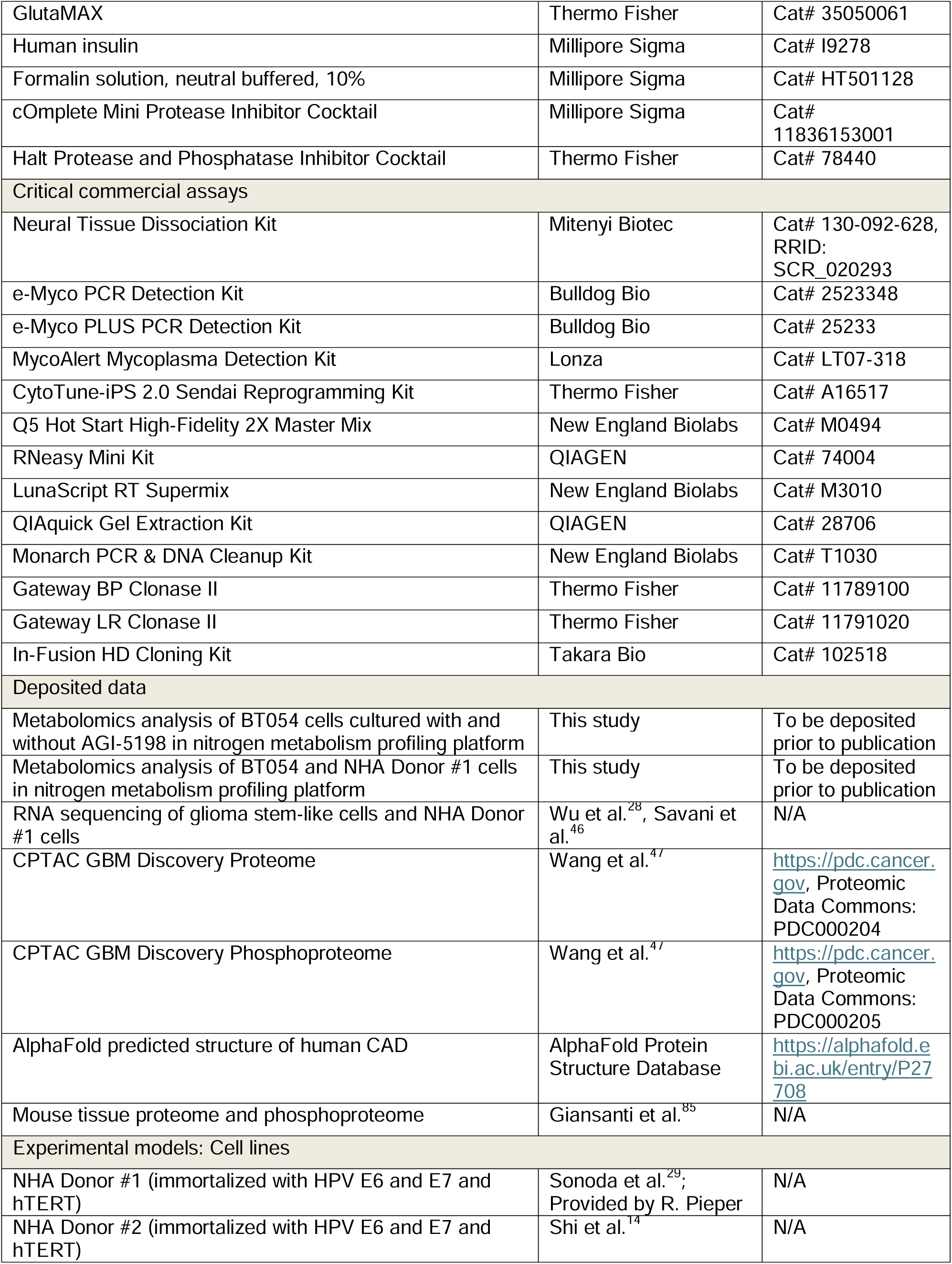
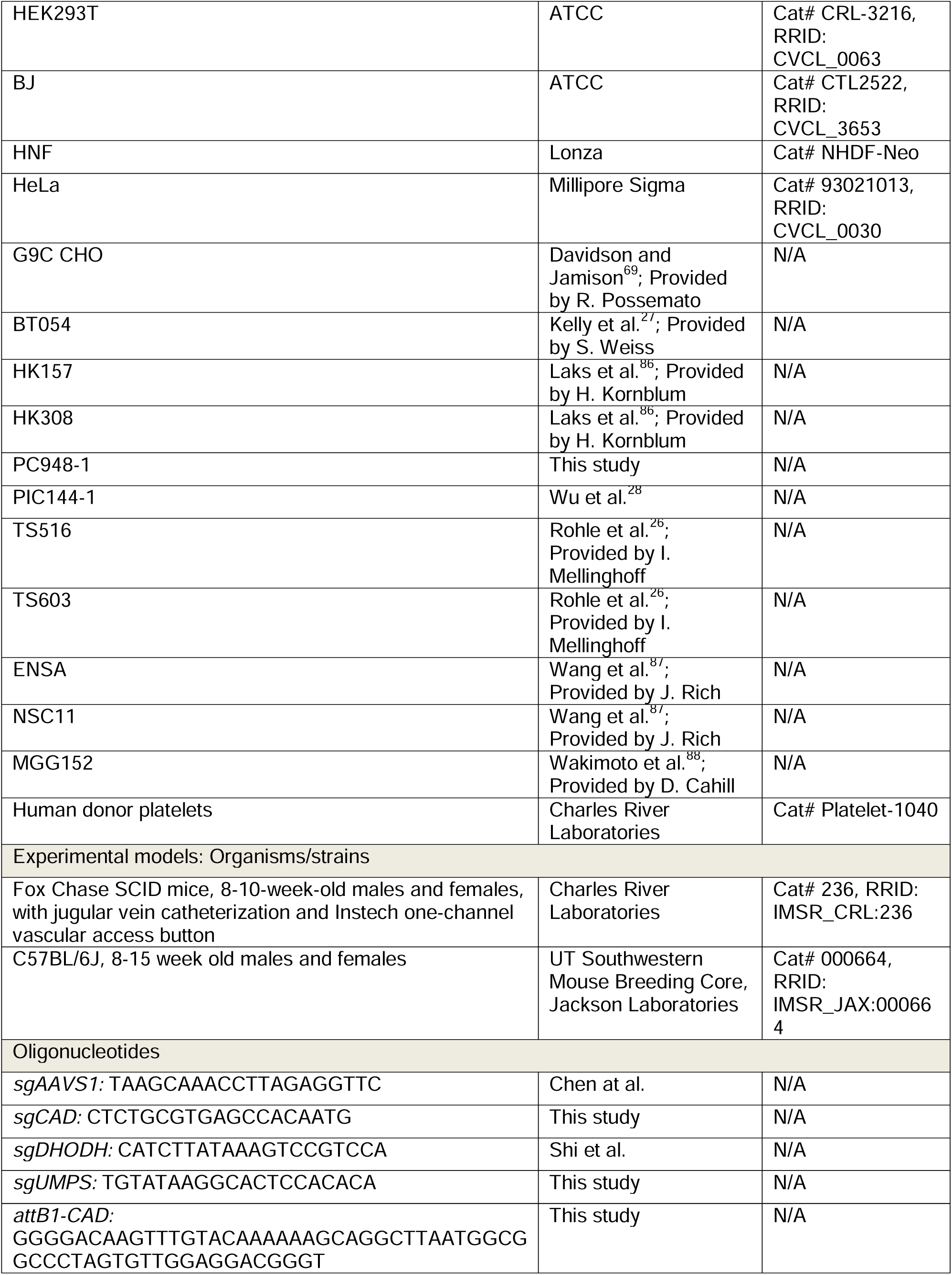
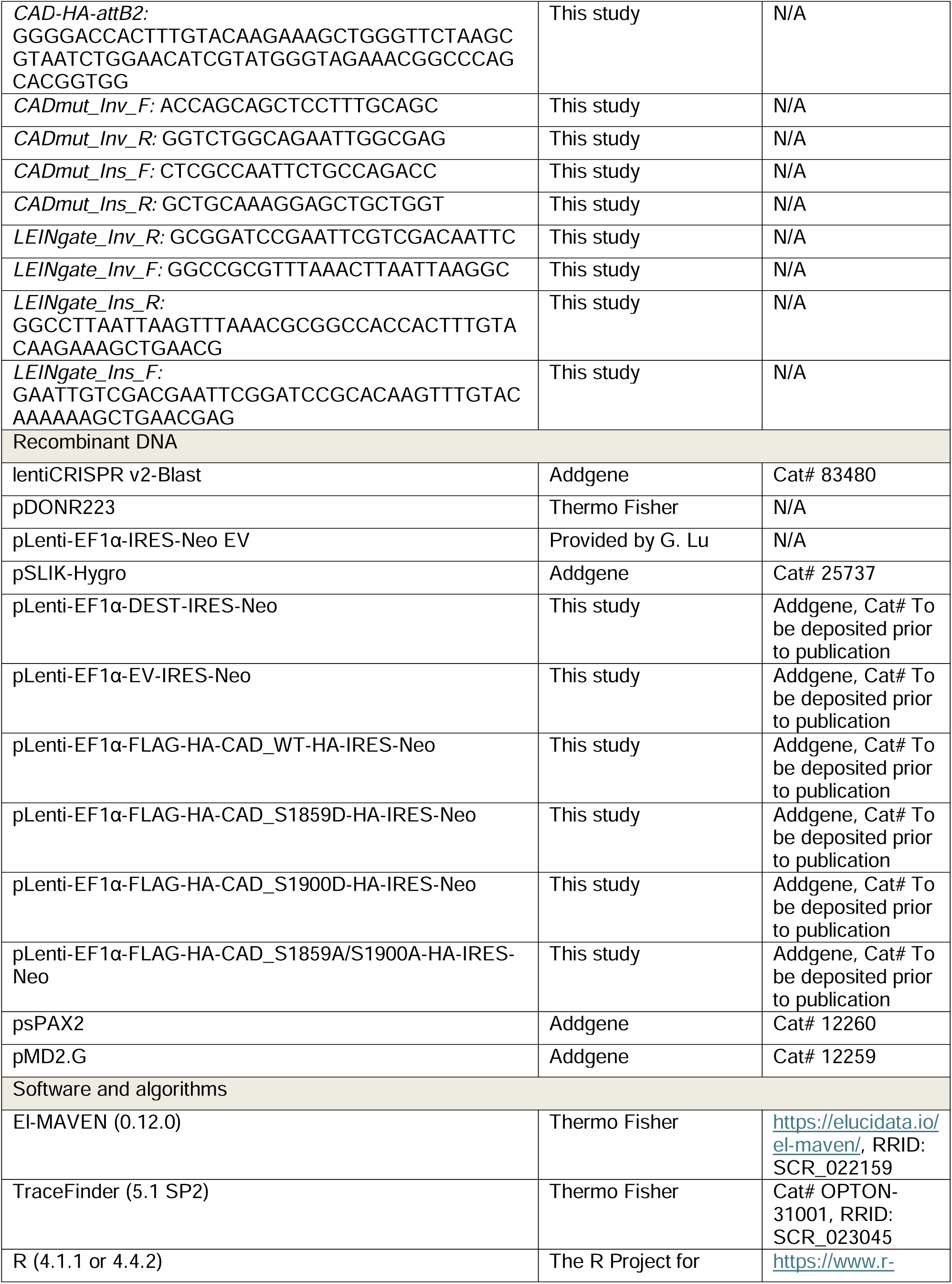
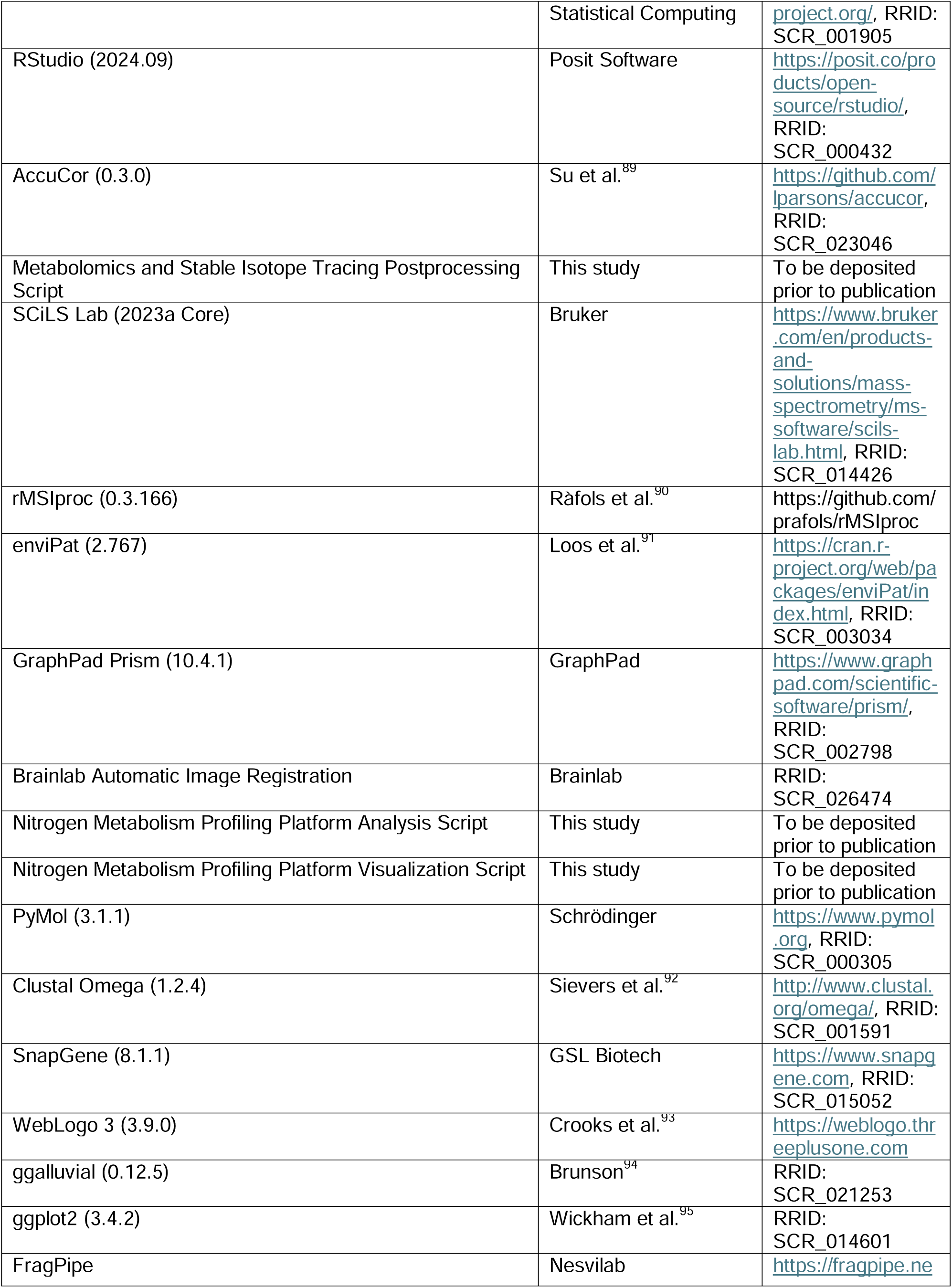
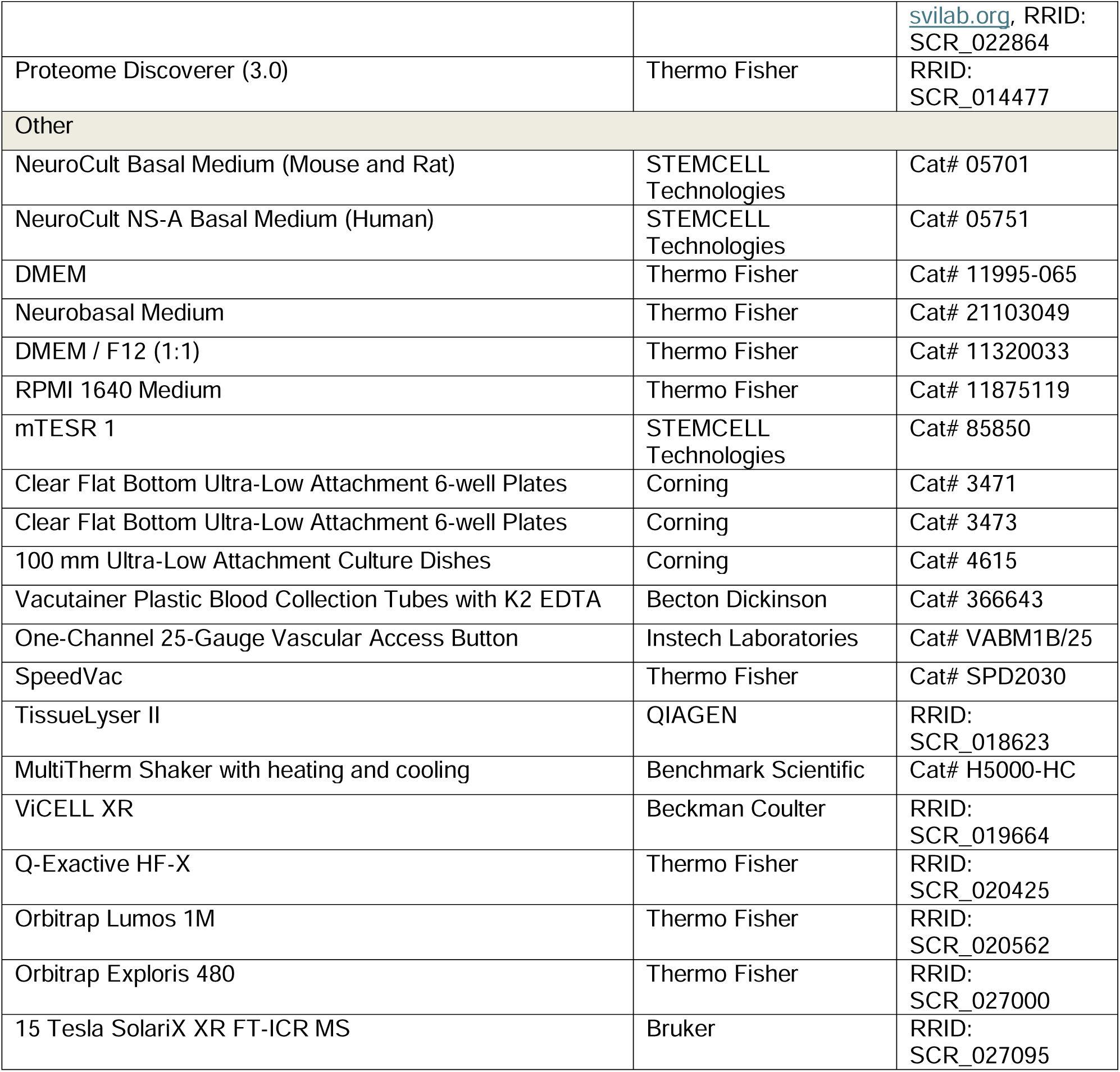

### Experimental Model and Study Participant Details

#### Cell lines

NHA Donor #1 cells^29^ (human astrocytes immortalized with HPV E6 and E7 and hTERT, sex unknown) were obtained from R. Pieper at the University of California San Francisco. NHA Donor #2 cells were generated as previously described^14^ from commercially obtained primary human astrocytes (Lonza CC-3187). HEK293T (female, ATCC CRL-3216, RRID: CVCL_0063), BJ (male, ATCC CRL-2522, RRID: CVCL_3653), HNF primary fibroblast (Lonza NHDF-Neo), and HeLa (female, Millipore Sigma 93021013, RRID: CVCL_0030) cells were obtained commercially. G9C CHO cells^69^ were a gift of R. Possemato at New York University. BT054^27^ (female, RRID: CVCL_N707) cells were obtained from S. Weiss at the University of Calgary. HK157 (female) and HK308^86^ (male) cells were obtained from H. Kornblum at the University of California Los Angeles. TS516 (sex unknown, RRID: CVCL_A5HY) and TS603^26^ (sex unknown, RRID: CVCL_A5HW) cells were obtained from I. Mellinghoff at Memorial Sloan-Kettering Cancer Center. ENSA (sex unknown) and NSC11^87^ (sex unknown) cells were obtained from J. Rich at University of North Carolina-Chapel Hill. MGG152^88^ (male) cells were obtained from D. Cahill at Massachusetts General Hospital.

Murine GSC lines PC948-1 and PIC144-1 were derived from autochthonous astrocytomas that formed in a modified version of a genetically engineered mouse model as previously described^14,28^. Briefly, a portion of murine brain tumor tissue was dissected and dissociated to a single-cell suspension using the Neural Tissue Dissociation Kit (Miltenyi Biotec 130-092-628, RRID: SCR_020293) by following the manufacturer’s instructions and then cultured for selection of neurosphere-forming cells.

All cell lines were routinely evaluated for mycoplasma contamination with the e-Myco Mycoplasma PCR Detection Kit (Bulldog Bio 2523348), e-Myco PLUS Mycoplasma PCR Detection Kit (Bulldog Bio 25233), or MycoAlert Mycoplasma Detection Kit (Lonza LT07-318) and confirmed to be negative. Cell line authentication was not performed because reference short term tandem repeat profiles have not been established for these lines. Sex and source of each line is stated above and listed as unknown if unreported in the original publication describing its derivation.

#### Cell culture

PC348-1 and PIC144-1 cells were cultured in NeuroCult Basal Medium (Mouse and Rat) with Proliferation Supplement (STEMCELL Technologies 05701), supplemented with EGF (20 ng/mL, GoldBio 1150-04), bFGF (20 ng/mL, GoldBio 1140-02), heparin (2 µg/mL, STEMCELL Technologies 07980), penicillin/streptomycin (100 U/mL and 100 μg/mL, respectively, Thermo Fisher 15140148), amphotericin B (250 ng/mL, GeminiBio 400104), and Plasmocin (250 ng/mL, InvivoGen ant-mpp) on ultra-low adherence plates (6-well plates: Corning 3471, 10 cm dishes: Corning 4615) in 5% CO_2_ and 5% oxygen at 37°C.

BT054, HK157, HK308, TS516, and TS603 cells were cultured in NeuroCult NS-A Basal Medium (Human) with Proliferation Supplement (STEMCELL Technologies 05751), supplemented with EGF (20 ng/mL, GoldBio 1150-04), bFGF (20 ng/mL, GoldBio 1140-02), heparin (2 µg/mL, STEMCELL Technologies 07980), penicillin/streptomycin (100 U/mL and 100 μg/mL, respectively, Thermo Fisher 15140148), amphotericin B (250 ng/mL, GeminiBio 400104), and Plasmocin (250 ng/mL, InvivoGen ant-mpp) on ultra-low adherence plates (6-well plates: Corning 3471, 10 cm dishes: Corning 4615) in 5% CO_2_ and at ambient oxygen at 37°C.

NHA Donor #1, NHA Donor #2, HNF, and BJ cells were cultured in DMEM (Thermo Fisher 11995-065) supplemented with fetal bovine serum (FBS, 10%, GeminiBio 100-106) and penicillin/streptomycin (100 U/mL and 100 μg/mL, respectively, Thermo Fisher 15140148) in 5% CO_2_ and at ambient oxygen at 37°C.

ENSA, NSC11, and MGG152 cells were cultured in Neurobasal Medium (Thermo Fisher 21103049) supplemented with glutamine (3mM, Thermo Fisher 25030081), B27 supplement (1×, Thermo Fisher 17504044), N2 supplement (0.25×, Thermo Fisher 17502048), EGF (20 ng/mL, GoldBio 1150-04), bFGF (20 ng/mL, GoldBio 1140-02), heparin (2 µg/mL, STEMCELL Technologies 07980), penicillin/streptomycin (50 U/mL and 50 μg/mL, respectively, Thermo Fisher 15-140-148), amphotericin B (125 ng/mL, GeminiBio 400104), and Plasmocin (0.25 µg/mL, InvivoGen ant-mpp) on ultra-low adherence plates (6-well plates: Corning 3471, 10 cm dishes: Corning 4615) in 5% CO_2_ and at ambient oxygen at 37°C.

HNF-derived NPCs were cultured in DMEM / F12 (1:1) (Thermo Fisher 11320033) supplemented with sodium pyruvate (1 mM, Thermo Fisher 11360070), B27 supplement minus vitamin A (1×, Thermo Fisher 12587010), N2 supplement (1×, Thermo Fisher 17502048), laminin (1 µg/mL, Thermo Fisher OB1415-01), and bFGF (20 ng/mL, GoldBio 1140-02).

G9C CHO cells were cultured in RPMI 1640 medium (Thermo Fisher 11875119) supplemented with fetal bovine serum (FBS, 10%, GeminiBio 100-106), penicillin/streptomycin (100 U/mL and 100 μg/mL, respectively, Thermo Fisher 15140148), and uridine (100 µM, Millipore Sigma U3003) in 5% CO_2_ and at ambient oxygen at 37°C.

All GSC and NSC lines were maintained as neurospheres and dissociated 1-2 times per week with Accutase (STEMCELL Technologies 07922). Only low passage GSC lines were used, and these cells were discarded after 3 months in culture to prevent genetic and/or phenotypic drift.

#### Generation of HNF-derived NPCs

Induced pluripotent stem cells (iPSCs) were generated from HNF fibroblasts using the CytoTune-iPS 2.0 Sendai Reprogramming Kit (Thermo Fisher Scientific A16517) per the manufacturer’s instructions. Briefly, HNF cells were cultured in their native medium until 30-60% confluent, then transduced at a multiplicity of infection of 5:5:3 (CytoTune 2.0 KOS, CytoTune 2.0 hc-Myc, CytoTune 2.0 hKlf4) with non-integrating Sendai virus to deliver the Yamanaka factors KLF4, OCT4, SOX2, and C-MYC. After 24 hours, medium was refreshed, and cells were cultured for 6 more days before being passaged on to irradiated MEFs. After day 7, cells were cultured in DMEM / F12 (1:1) (Thermo Fisher 11320033) supplemented with KnockOut Serum Replacement (20%, Thermo Fisher 10828028), non-essential amino acids (1%, Thermo Fisher 11140050), penicillin/streptomycin (100 U/mL and 100 μg/mL, respectively, Thermo Fisher 15140148), bFGF (10 ng/mL, GoldBio 1140-02), 2-mercaptoethanol (0.1 mM, Thermo Fisher BP176100) with daily media changes. iPSC colonies emerged within 12-21 days. Eight clonal iPSC lines were derived and two were expanded and characterized. Pluripotency was confirmed by alkaline phosphatase staining and immunofluorescence for OCT4, SOX2, NANOG, TRA-1-60, and TRA-1-81. Clearance of Sendai virus was verified by RT-PCR at passage 10. Only virus-free clones were used for subsequent differentiation. To generate neural progenitor cells (NPCs), iPSCs were cultured in mTeSR 1 (STEMCELL Technologies 85850) on plates coated with Matrigel (Corning 354234) diluted 1:100 in phosphate buffered saline. Cells were differentiated to NPCs by transition to DMEM / F12 (1:1) (Thermo Fisher 11320033) supplemented with B27 supplement minus vitamin A (1×, Thermo Fisher 12587010), N2 supplement (1×, Thermo Fisher 17502048), LDN193189 (0.1 µM, Stemgent 04-0074), and SB431542 (10 µM, Stemgent 04-0010). Neural rosettes appeared by days 7-10 and were manually isolated on day 14. Rosettes were transferred to plates coated with 10 µg/mL Poly-L-Ornithine (Millipore Sigma P3655) in water and 5 µg/mL laminin (Thermo Fisher 23017015) in phosphate buffered saline and cultured in DMEM / F12 (1:1) (Thermo Fisher 11320033) supplemented with sodium pyruvate (1 mM, Thermo Fisher 11360070), B27 supplement minus vitamin A (1×, Thermo Fisher 12587010), N2 supplement (1×, Thermo Fisher 17502048), laminin (1 µg/mL, Thermo Fisher OB1415-01), bFGF (20 ng/mL, GoldBio 1140-02). NPC identity was confirmed by immunostaining for NESTIN and SOX2.

#### Animals

All care and treatment of experimental animals were carried out in strict accordance with Good Animal Practice as defined by the US Office of Laboratory Animal Welfare and approved by the UT Southwestern Medical Center (protocols 2017-101840, 2019-102795, and 2022-102897) Institutional Animal Care and Use Committee. Animal welfare assessments were carried out daily during treatment periods. Animals were housed in a pathogen-free environment between 20-26°C and at 30-70% humidity, with a 12 hour:12 hour light:dark cycle. Fox Chase SCID (Charles River 236, RRID: IMSR_CRL:236) mice pre-catheterized in the jugular vein with a one-channel 25-gauge vascular access button (Instech Laboratories VABM1B/25) were obtained from Charles River Laboratories at 8-10 weeks of age. C57BL/6J (RRID: IMSR_JAX:000664) mice were obtained from the UT Southwestern Mouse Breeding Core or Jackson Laboratories. Mice were housed together (2-5 mice of the same sex per cage) and provided free access to chow diet (Teklad 2916) and water. Catheters were flushed with 20 USP units/mL heparin in normal saline (Instech Laboratories USP-HS-020-0.5-NJ1P-50) every 3-5 days prior to infusion experiments.

#### MGG152 orthotopic xenografts

Littermates of the same sex were randomized to receive or not receive orthotopic xenografts prior to implantations. Orthotopic glioma cell implantations were performed 1-2 weeks after receipt of mice. Mice were provided analgesia by subcutaneous injection of buprenorphine SR (0.1 mg/kg) and anesthetized with isoflurane, then immobilized using a stereotactic frame. An incision was made to expose the skull surface, and a hole was drilled into the skull 3 mm anterior and 2 mm lateral to the lambda, 2.5 mm below the surface of the brain. 1 × 10^5^ MGG152 cells suspended in 1-3 μL of culture medium were injected into the brain through the hole using a 5 μL syringe (Hamilton 7634-01). The skin was closed with surgical clips. Mice were provided additional analgesia by subcutaneous injection of carprofen (5 mg/kg) every 24 hours until 72 hours after the procedure. Mice were euthanized after infusion or when they either displayed neurological symptoms or became moribund.

#### Human subjects

The study was conducted according to the principles of the Declaration of Helsinki. Patient tissue and blood were collected following ethical and technical guidelines on the use of human samples for biomedical research at the University of Pittsburgh Medical Center after informed patient consent under a protocol approved by the University of Pittsburgh Medical Center’s Institutional Review Board. All patient samples were de-identified before processing. All patient samples and organoids were diagnosed and graded according to the *2021 WHO Classification of Tumours of the Central Nervous System (CNS)*, 5^th^ edition^96^. Human tumor explants for stable isotope tracing were derived from patients with the following ages, sexes, clinical diagnoses, and resection statuses: UPMC1232 (64; male; Glioblastoma, IDH-wildtype; index resection), UPMC3313 (72; male; Glioblastoma, IDH-wildtype; index resection), UPMC5565 (74; female; Glioblastoma, IDH-wildtype; index resection), UPMC7602 (67; female; Glioblastoma, IDH-wildtype; recurrence), UPMC9041 (62; male; Glioblastoma, IDH-wildtype; index resection). Human non-neoplastic brain explants for stable isotope tracing were derived from patients with the following ages, sexes, and clinical sources: UPMC1068 (65; female; left occipital lobe, tissue remote from metastatic lung adenocarcinoma with neuroendocrine differentiation status post radiosurgery), UPMC4875 (69; female; tissue remote from metastatic colon adenocarcinoma), UPMC6777 (69; male; ventriculoperitoneal shunt placement), UPMC9230 (74; male; ventriculoperitoneal shunt placement), UPMC9413 (68; male; ventriculoperitoneal shunt placement). Further clinical details are available in Table S4. Human platelets (Charles River Platelet-1040) were obtained from Charles River Laboratories from a 28-year-old female donor.

### Method Details

#### Vectors

lentiCRISPR v2-Blast (Addgene 83480, a gift from Mohan Babu) was digested with *BsmBI-v2* (New England Biolabs R0739), and sgRNAs targeting the AAVS1 safe harbor locus (TAAGCAAACCTTAGAGGTTC)^97^, CAD (CTCTGCGTGAGCCACAATG), DHODH (CATCTTATAAAGTCCGTCCA)^14^, or UMPS (TGTATAAGGCACTCCACACA) were ligated into this backbone to generate a construct expressing Cas9 and the desired sgRNA. Reaction mixtures were transformed into XL10-Gold Ultracompetent Cells (Agilent 200315) and confirmed by whole plasmid sequencing (Plasmidsaurus).

CAD WT (RefSeq: NM_004341.5) cDNA was amplified by PCR with Q5 Hot Start High-Fidelity 2X Master Mix polymerase (New England Biolabs M0494). A cDNA library prepared from NHA Donor #1 cells with the RNeasy Mini Kit (QIAGEN 74004) and LunaScript RT Supermix (New England Biolabs M3010) was used to provide template. The following primers were designed to append 5’ *attB1* and 3’ HA-tag and *attB2* sites:

*attB1-CAD:* GGGGACAAGTTTGTACAAAAAAGCAGGCTTAATGGCGGCCCTAGTGTTGGAGGACGGGT
*CAD-HA-attB2:* GGGGACCACTTTGTACAAGAAAGCTGGGTTCTAAGCGTAATCTGGAACATCGTATGGGTAGAAACGGCCCAGCACGGTGG

PCR product was gel-purified with the QIAquick Gel Extraction Kit (QIAGEN 28706), PCR purified with the Monarch PCR & DNA Cleanup Kit (New England Biolabs T1030) and cloned into the Gateway vector pDONR223 by BP reaction (Thermo Fisher 11789100). The resultant product was transformed into HB101 Competent Cells (Promega L2015). N-terminal FLAG and HA tags were added by restriction cloning with *ApaI* (New England Biolabs R0114) and *SalI* (New England Biolabs R3138) with a gBlock synthesized by Twist Biosciences to generate pENTR223-FLAG-HA-CAD-HA plasmid. The resultant product was transformed into XL10-Gold Ultracompetent Cells (Agilent 200315).

CAD S1859D, S1900D, and S1859/S1900A mutant cDNAs were generated by In-Fusion cloning using pENTR223-FLAG-HA-CAD-HA. Inverse PCR with Q5 Hot Start High-Fidelity 2X Master Mix polymerase (New England Biolabs M0494) of pENTR223-FLAG-HA-CAD-HA plasmid used the following primers:

*CADmut_Inv_F:* ACCAGCAGCTCCTTTGCAGC
*CADmut_Inv_R:* GGTCTGGCAGAATTGGCGAG

Insert gBlocks containing CAD S1859D, S1900D, or S1859/S1900A mutations were synthesized by Twist Biosciences and amplified using the following insert primers:

*CADmut_Ins_F:* CTCGCCAATTCTGCCAGACC
*CADmut_Ins_R:* GCTGCAAAGGAGCTGCTGGT

Amplified inserts and inverse PCR products were purified with the Monarch PCR & DNA Cleanup Kit (New England Biolabs T1030) and used in an In-Fusion exchange reaction (Takara 102518), then transformed into XL10-Gold Ultracompetent Cells (Agilent 200315). Final pENTR223-FLAG-HA-CAD-HA product for WT, S1859D, S1900D, and S1859/S1900A mutants were confirmed by whole plasmid sequencing (Plasmidsaurus).

pLenti-EF1α-DEST-IRES-Neo was generated by In-Fusion cloning from pLenti-EF1α-IRES-Neo EV (provided by Dr. Gang Lu). Inverse PCR with Q5 Hot Start High-Fidelity 2X Master Mix polymerase (New England Biolabs M0494) of pLenti-EF1α-IRES-Neo EV used the following primers:

*LEINgate_Inv_R:* GCGGATCCGAATTCGTCGACAATTC
*LEINgate_Inv_F:* GGCCGCGTTTAAACTTAATTAAGGC

The Gateway cassette was amplified from pSLIK-Hygro (Addgene 25737, a gift from Iain Fraser) using the following primers:

*LEINgate_Ins_R:* GGCCTTAATTAAGTTTAAACGCGGCCACCACTTTGTACAAGAAAGCTGAACG
*LEINgate_Ins_F:* GAATTGTCGACGAATTCGGATCCGCACAAGTTTGTACAAAAAAGCTGAACGAG

Amplified inserts and inverse PCR products were purified with the Monarch PCR & DNA Cleanup Kit (New England Biolabs T1030) and used in an In-Fusion exchange reaction (Takara 102518), then transformed into XL10-Gold Ultracompetent Cells (Agilent 200315). Final pLenti-EF1α-DEST-IRES-Neo product was confirmed by whole plasmid sequencing (Plasmidsaurus).

CAD cDNAs were cloned into the lentiviral vector pLenti-EF1α-DEST-IRES-Neo by Gateway cloning. pENTR223-EV, pENTR223-FLAG-HA-CAD_WT-HA, pENTR223-FLAG-HA-CAD_S1859D-HA, pENTR223-FLAG-HA-CAD_S1900D-HA, and pENTR223-FLAG-HA-CAD_S1859A/S1900A-HA were separately mixed with pLenti-EF1α-DEST-IRES-Neo and LR reactions (Thermo Fisher 11791020) performed. Resultant pLenti-EF1α-EV-IRES-Neo, pLenti-EF1α-FLAG-HA-CAD_WT-HA-IRES-Neo, pLenti-EF1α-FLAG-HA-CAD_S1859D-HA-IRES-Neo, pLenti-EF1α-FLAG-HA-CAD_S1900D-HA-IRES-Neo, and pLenti-EF1α-FLAG-HA-CAD_S1859A/S1900A-HA-IRES-Neo lentiviral vectors for mammalian cell expression were transformed into HB101 Competent Cells (Promega L2015) and confirmed by whole plasmid sequencing (Plasmidsaurus).

#### Transfection and viral transduction

Lentiviral particles were produced by cotransfection of HEK293T cells with expression vectors and packaging plasmids psPAX2 (Addgene 12260, a gift from Didier Trono) and pMD2.G (Addgene 12259, a gift from Didier Trono) in a ratio of 4:3:1 using TransIT-LT1 transfection reagent (Mirus Bio MIR2 B-304). Virus-containing media were collected 48 and 72 hours after transfection, passed through a 0.45 μm filter (Corning 431220), divided into 1 mL aliquots, and frozen at −80°C until use.

NHA Donor #1, G9C CHO, or HeLa cells were plated at a density of 0.15 × 10^6^ cells per well in a 6-well plate. The next day, polybrene (8 µg/mL, MedChemExpress HY-112735) was added to each well in addition to 1 mL viral supernatant. Plates were centrifuged at 4,000 × g for 30 minutes at room temperature and incubated overnight. The following day, cells were expanded and replated in a 10 cm dish. Stable cell lines were selected in 1 mg/mL G418 (GoldBio G-418) or 2 μg/mL blasticidin (GoldBio B-800) based on the selection cassette present in each vector. After initial selection, stable cell lines were maintained in 400 µg/mL G418 (GoldBio G-418) or 800 ng/mL blasticidin (GoldBio B-800) based on the selection cassette present in each vector.

#### Preparation of HPLM and tracer HPLM

HPLM was prepared as previously described^12,13^. The compositions and preparation protocols for the 19 constituent pools (both unlabeled and tracer formulations) are provided in the “Pool” and “Preparation” columns of Table S1. For tracer HPLM used in profiling experiments, stock solutions were generated identically to the unlabeled version, substituting the labeled metabolite for its unlabeled counterpart at equimolar concentrations. The tracer HPLM library was assembled by mixing thawed frozen stocks of unlabeled HPLM pools, replacing one pool with its labeled version per tracing condition. Pools designated for fresh preparation were prepared at the time of library assembly. Media was adjusted to pH 7.4 using NaOH or HCl, brought to the final volume, sterile filtered with a 0.22 µm PES filter (Millipore Sigma SCGP00525, Corning 431153, Corning 431097, or Corning 431098) and supplemented as described below.

For experiments in GSCs, NSCs, and NPCs, HPLM was prepared without glutamate, with the exception of conditions in which glutamate was traced, For experiments in GSCs, NSCs, and HNF-derived NPCs HPLM was supplemented with B27 supplement (1×, Thermo Fisher 17504044), N2 supplement (0.25×, Thermo Fisher 17502048), EGF (20 ng/mL, GoldBio 1150-04), bFGF (20 ng/mL, GoldBio 1140-02), heparin (2 µg/mL, STEMCELL Technologies 07980), penicillin/streptomycin (100 U/mL and 100 μg/mL, respectively, Thermo Fisher 15-140-148), amphotericin B (250 ng/mL, GeminiBio 400104), and Plasmocin (250 ng/mL, InvivoGen ant-mpp). For experiments in differentiated cells, HPLM was supplemented with dialyzed FBS (10%, GeminiBio 100-108) and penicillin/streptomycin (100 U/mL and 100 μg/mL, respectively, Thermo Fisher 15140148).

#### Preparation of cultured cells for LC-MS

BT054 cell samples treated with DMSO or AGI-5198 (except those involving low-throughput ^15^N-BCAA tracing) were prepared as previously described^14^. Cells were washed with ice-cold saline and snap frozen in liquid nitrogen. Prior to LC-MS analysis, samples were suspended in ice-cold 80% methanol (Thermo Fisher A456), vortexed for 20 minutes at 4°C, and centrifuged for 10 minutes at 17,000-21,100 × g at 4°C. Samples were then dried using a SpeedVac (Thermo Fisher SPD2030). Dried metabolites were resuspended in 1 µL ice-cold 80% acetonitrile (Thermo Fisher A9554) per 10,000 cells, vortexed for 20 minutes at 4°C, and centrifuged for 10 minutes at 17,000-21,100 × g at 4°C. The resultant supernatant was then analyzed by LC-MS.

Platelets were washed with ice-cold normal saline and snap frozen in liquid nitrogen. Prior to LC-MS analysis, samples were resuspended in 55 µL ice-cold 80% acetonitrile (Thermo Fisher A9554), vortexed for 20 minutes at 4°C, and centrifuged for 10 minutes at 17,000-21,100 × g at 4°C. The supernatant was transferred to a fresh tube and again centrifuged for 10 minutes at 17,000-21,100 × g at 4°C. The resultant supernatant was then analyzed by LC-MS.

All other cell samples were washed with ice-cold normal saline and snap frozen in liquid nitrogen. Prior to LC-MS analysis, samples were suspended in 1 µL ice-cold 80% acetonitrile (Thermo Fisher A9554) per 1,000 cells, vortexed for 20 minutes at 4°C, and centrifuged for 10 minutes at 17,000-21,100 × g at 4°C. The supernatant was transferred to a fresh tube and again centrifuged for 10 minutes at 17,000-21,100 × g at 4°C. The resultant supernatant was then analyzed by LC-MS.

#### Preparation of media for LC-MS

Unconditioned HPLM stocks generated for the nitrogen metabolism profiling platform were collected and snap frozen in liquid nitrogen. Prior to LC-MS analysis, 50 µL of media was added to a 5 mL tube. 1 mL 100% methanol (Thermo Fisher A456), 1 mL of 100% chloroform (Millipore Sigma 650471), and 1 mL of water were added, with brief vortexing after each addition. Samples were centrifuged for 5 minutes at 410 × g at 4°C. 1 mL of the aqueous phase was transferred to a 1.5 mL tube. Samples were dried using a SpeedVac (Thermo Fisher SPD2030). Dried metabolites were resuspended in 100 µL ice-cold 80% acetonitrile (Thermo Fisher A9554), vortexed for 20 minutes at 4°C, and centrifuged for 10 minutes at 17,000-21,100 × g at 4°C. The resultant supernatant was then analyzed by LC-MS.

Conditioned media samples were collected and snap frozen in liquid nitrogen. Prior to LC-MS analysis, media was diluted 1:100 in ice-cold 80% acetonitrile (Thermo Fisher A9554) and vortexed for 20 minutes at 4°C, then centrifuged for 10 minutes at 17,000-21,100 × g at 4°C. The supernatant was transferred to a fresh tube and again centrifuged for 10 minutes at 17,000-21,100 × g at 4°C. The resultant supernatant was then analyzed by LC-MS.

#### Preparation of blood for LC-MS

Blood samples were collected from mice and snap frozen in liquid nitrogen. Prior to LC-MS analysis, blood was diluted 1:100 in ice-cold 80% acetonitrile (Thermo Fisher A9554) and vortexed for 20 minutes at 4°C, then centrifuged for 10 minutes at 17,000-21,100 × g at 4°C. The supernatant was transferred to a fresh tube and again centrifuged for 10 minutes at 17,000-21,100 × g at 4°C. The resultant supernatant was then analyzed by LC-MS.

#### Preparation of tissue for LC-MS

Adult mouse tissues were collected and snap frozen in liquid nitrogen. 50 µL ice-cold 80% methanol (Thermo Fisher A456) per mg of tissue was added to samples, which were then homogenized using a TissueLyser II (Qiagen, RRID: SCR_018623). Samples were vortexed for 20 minutes at 4°C, then centrifuged for 10 minutes at 17,000-21,100 × g at 4°C. The supernatant was transferred to a fresh tube and again centrifuge for 10 minutes at 17,000-21,100 × g at 4°C. 200 µL of supernatant was transferred to a fresh tube, then dried using a SpeedVac (Thermo Fisher SPD2030). Dried metabolites were resuspended in 100 µL ice-cold 80% acetonitrile (Thermo Fisher A9554), vortexed for 20 minutes at 4°C, and centrifuged for 10 minutes at 17,000-21,100 × g at 4°C. The resultant supernatant was then analyzed by LC-MS.

Fetal mouse tissues were collected and snap frozen in liquid nitrogen. Ice-cold 80% acetonitrile (Thermo Fisher A9554) was added to samples, which were then homogenized using a rubber dounce homogenizer. Samples were flash frozen 3 times in liquid nitrogen and then centrifuged for 10 minutes at 17,000 × g at 4°C. Supernatants were analyzed for protein content using a BCA assay, normalized to 70 µg/mL, then analyzed by LC-MS.

Human samples for steady-state metabolomics were collected from the operating room, suspended in ice-cold Hibernate A (BrainBits HA), and snap frozen. Samples were prepared as previously described^98^, with some minor modifications. Briefly, samples were thawed and washed in 1 mL ice-cold normal saline. 50 µL ice-cold 80% methanol (Thermo Fisher A456) per mg of tissue was added to samples, which were then homogenized using a TissueLyser II (Qiagen, RRID: SCR_018623). 520 µL *tert*-Butyl methyl ether (Millipore Sigma 650560) and 380 µL water were added to 500 µL of homogenate in a glass vial and mixture was vortexed at room temperature for 1 hour at 1,000 rpm in a thermal mixer (Benchmark Scientific H5000-HC). Samples were centrifuged for 10 minutes at 1,000 × g at 4°C. 800 µL of the lower aqueous phase and 300 µL of the upper organic phase were combined in a new tube, avoiding interlayer debris. Samples were then dried using a SpeedVac (Thermo Fisher SPD2030). Dried metabolites were resuspended in 100 µL ice-cold 80% acetonitrile (Thermo Fisher A9554) and an internal standard mix of 3 µM ^13^C_5_-glutamine (Cambridge Isotope Laboratories CLM-1822), 3 µM ^15^N-valine (Cambridge Isotope Laboratories NLM-316), 3 µM ^15^N-methionine (Cambridge Isotope Laboratories NLM-752), and 3 µM chloramphenicol (Millipore Sigma C0378) was added. Samples were sonicated in a room temperature water bath for 10 minutes to ensure metabolite dissolution. Samples were then centrifuged for 10 minutes at 21,100 × g at 4°C. The resultant supernatant was then analyzed by LC-MS.

Human tissue explants were snap frozen following stable isotope tracing. 50 µL ice-cold 80% acetonitrile (Thermo Fisher A9554) per mg of tissue was added to samples, which were then homogenized using a rubber dounce homogenizer. Samples were vortexed for 20 minutes at 4°C, then centrifuged for 10 minutes at 17,000-21,100 × g at 4°C. The resultant supernatant was again centrifuged for 10 minutes at 17,000-21,100 × g at 4°C. The resultant supernatant was then analyzed by LC-MS.

#### Metabolite quantification by LC-MS

10-20 µL of acetonitrile-resuspended metabolite extract was injected and analyzed with a Q-Exactive HF-X (RRID: SCR_020425), Orbitrap Fusion Lumos 1M (RRID: SCR_020562), or Orbitrap Exploris 480 (RRID: SCR_027000) hybrid quadrupole-orbitrap mass spectrometer (Thermo Fisher) coupled to a Vanquish Flex UHPLC system (Thermo Fisher), as described previously^21,22^. Chromatographic resolution of metabolites was achieved using a Millipore ZIC-pHILIC column using a linear gradient of 10 mM ammonium formate pH 9.8 and acetonitrile. Spectra were acquired with a resolving power of 60,000, 120,000, or 240,000 full width at half maximum (FWHM), a scan range set to 70-1,050 or 80-1,200 m/z, and polarity switching or a targeted selected ion monitoring (tSIM) scan for low-abundance metabolites. Data-dependent MS/MS data was acquired on unlabeled pooled samples to confirm metabolite IDs when necessary. All solvents used for MS experiments were LC-MS Optima grade (Thermo Fisher).

Peaks were integrated using El-Maven 0.12.0 software (Elucidata, RRID: SCR_022159) or TraceFinder 5.1 SP2 software (Thermo Fisher OPTON-31001, RRID: SCR_023045). Total ion counts were quantified using TraceFinder 5.1 SP2 software (Thermo Fisher OPTON-31001, RRID: SCR_023045). Peaks were normalized using probabilistic quotient normalization^99^ and total ion counts using R (RRID: SCR_001905). For stable isotope tracing studies, correction for natural abundance of metabolite labeling was performed using AccuCor^89^ 0.3.0 (RRID: SCR_023046) in R (RRID: SCR_001905).

#### MALDI-MSI tissue preparation and microscopy

Tissue samples for MALDI-MSI analysis were collected from mice and flash frozen for 3 cycles of 3 seconds each, separated by 2 seconds at room temperature, in liquid nitrogen. Samples were cryo-sectioned in the coronal plane to 10 µm thickness and thaw-mounted onto indium tin oxide slides. Serial sections were collected for H&E staining and imaged using a 10× objective (Zeiss Observer Z.1, Oberkochen, Germany).

#### MALDI matrix preparation

Glutamine and UMP were imaged using 1,5-diaminonaphthalene hydrochloride (4.3 mg/mL) matrix solution prepared in 4.5/5/0.5 HPLC grade water/ethanol/1 M HCl (v/v/v). The matrix was applied using a four-pass cycle with 0.09 mL/min flow rate, spray nozzle velocity (1200 mm/min), spray nozzle temperature (75°C), nitrogen gas pressure (10 psi), and track spacing (2 mm).

#### Glutamine and UMP MALDI-MSI

Glutamine and UMP were imaged from serial tissue sections. Mass spectrometry imaging data was acquired using a 15 Tesla SolariX XR FT-ICR MS (Bruker, RRID: SCR_027095). Instrument parameters were set to negative ion mode, and the mass range was from m/z 46.07 to 3000. The laser raster size was 100 µm and each sampling point consisted of 200 laser shots at a laser power of 21% (arbitrary scale) with a laser repletion rate of 100 Hz. Continuous Accumulation of Selected Ions mode was used with setting Q1 to m/z 150 with an isolation window of 200. Instrument calibration was carried out using a tune mix solution (Agilent) with the electrospray ionization source. A ^15^N glutamate internal standard was used for online calibration during acquisition. MSI data analysis was completed using SCiLS Lab 2024a Pro (Bruker, RRID: SCR_014426) with normalization to ^15^N glutamate. Glutamine and UMP were putatively annotated based on an accurate mass with Δppm < 0.1 and MS/MS measurements.

#### MALDI-MSI isotopic labeling analysis

Acquired MALDI-MSI datasets were preprocessed, visualized, and exported to imZML^100^ format using SCiLS Lab 2023a Core (Bruker, RRID: SCR_014426). Per-pixel natural abundance correction and generation of fractional enrichment images were performed using a pipeline implemented in R 4.1.1 (RRID: SCR_001905). Data were imported with rMSIproc^90^ 0.3.166 and theoretical mass spectra or each isotopologue were calculated using enviPat^91^ 2.767 (RRID: SCR_003034). Natural abundance correction was then performed by solving a system of linear equations incorporating theoretical isotopologue ratios alongside the experimental spectra at each pixel^101^.

#### Nitrogen metabolism profiling of AGI-5198- or DMSO-treated BT054 cells

BT054 cells were plated in 2 mL 6-well ultra-low adherence plates (0.5 × 10^6^ cells per well) in NeuroCult NS-A Basal Medium (Human) prepared as described above. After 24 hours, media was changed to 2 mL of 50% NeuroCult (Human) and 50% unlabeled HPLM prepared as described above. 24 hours later, media was changed to 100% unlabeled HPLM. Either DMSO or 3 µM AGI-5198 (MedKoo Biosciences 406264) was added. 24 hours later, media was again changed to fresh 100% unlabeled HPLM with either DMSO or 3µM AGI-5198. 24 hours later, was changed to fresh 100% HPLM with one nitrogen-containing metabolite per sample exchanged for an equimolar amount of its ^15^N-labeled counterpart (see Table S1) and either DMSO or 3µM AGI-5198. After 18 hours, samples were harvested and prepared for LC-MS analysis as described above.

#### Nitrogen metabolism profiling of NHA and BT054 cells

NHA Donor #1 cells were plated in 6-well plates (0.025 × 10^6^ cells per well) in 2 mL DMEM prepared as described above. BT054 cells were plated in 6-well ultra-low adherence plates (0.2 × 10^6^ cells per well) in 2 mL NeuroCult NS-A Basal Medium (Human) prepared as described above. After 24 hours, 2 mL unlabeled HPLM prepared as described above was added to produce a mixture of 50% native medium and 50% HPLM. 24 hours later, media was changed to 100% unlabeled HPLM (10 mL for NHA Donor #1 cells and 4.5 mL for BT054 cells). 24 hours later, was changed to fresh 100% HPLM with one nitrogen-containing metabolite per sample exchanged for an equimolar amount of its ^15^N-labeled counterpart (see Table S1, 10 mL for NHA Donor #1 cells and 4.5 mL for BT054 cells). After 18 hours, samples were harvested and prepared for LC-MS analysis as described above.

#### Pyrimidine synthesis tracing in cells

Differentiated cells were plated in 6-well plates (0.025 × 10^6^ cells per well for NHA Donor #1, 0.15 × 10^6^ cells per well for NHA Donor #2 and G9C CHO, 0.05 × 10^6^ cells for BJ, and 0.125 × 10^6^ cells per well for HNF) in 2 mL RPMI 1640 (G9C CHO cells) or DMEM (all other differentiated cells) prepared as described above. GSCs were plated in 6-well ultra-low adherence plates (0.2 × 10^6^ cells per well) in 2mL NeuroCult NS-A Basal Medium (Human) prepared as described above. NSCs were plated in 6-well ultra-low adherence plates (0.2 × 10^6^ cells per well) in 2 mL Neurobasal Medium prepared as described above. HNF-derived NPCs were plated in 6-well plates (0.15 × 10^6^ cells per well) in 2 mL DMEM / F12 (1:1) prepared as described above. After 24 hours, 2 mL unlabeled HPLM prepared as described above was added to produce a mixture of 50% native medium and 50% HPLM. 24 hours later, media was changed to 100% unlabeled HPLM (10 mL for differentiated cells, 4.5 mL for GSCs, NSCs, and HNF-derived NPCs). 24 hours later, media was changed to fresh 100% HPLM (10 mL for differentiated cells, 4.5 mL for GSCs, NSCs, and HNF-derived NPCs) modified as follows: glutamine exchanged for an equimolar amount of amide-^15^N-glutamine (Cambridge Isotope Laboratories NLM-557), glutamine exchanged for an equimolar amount of amide-^15^N-glutamine (Cambridge Isotope Laboratories NLM-557) and uridine removed, or uridine exchanged for an equimolar amount of ^15^N_2_ uridine (Cambridge Isotope Laboratories NLM812). After incubation with tracer for indicated timepoints, samples were harvested and prepared for LC-MS analysis as described above.

For stable isotope tracing experiments with altered growth factor sources, 24 hours after plating in their native media NHA Donor #1 cells were cultured in conditions described for GSCs, NSCs, and HNF-derived NPCs and BT054 cells were cultured in conditions described differentiated cells. For stable isotope tracing experiments with altered adherence conditions, ENSA and NSC11 cells were plated on 6-well plates coated with Matrigel (Corning 354234) diluted 1:100 in phosphate buffered saline. For stable isotope tracing experiments with uridine deprivation preconditioning, during the final 24 hours prior to culture in amide-^15^N-glutamine HPLM, NHA Donor #1 cells were instead cultured in 10 mL 100% HPLM with uridine removed.

#### HPLM metabolite depletion analysis

NHA Donor #1 cells were plated in 6-well plates (Figure S2A: 0.025 × 10^6^ cells per well, Figure 5A: 0.2 × 10^6^ cells per well) in 2 mL DMEM prepared as described above. BT054 cells were plated in 6-well ultra-low adherence plates (0.2 × 10^6^ cells per well) in 2 mL NeuroCult NS-A Basal Medium (Human) prepared as described above. After 24 hours, 2 mL unlabeled HPLM prepared as described above was added to produce a mixture of 50% native medium and 50% HPLM. 24 hours later, media was changed to 100% unlabeled HPLM (10 mL for NHA Donor #1 in Figure S2A, 4.5 mL for NHA Donor #1 in Figure 5C and BT054). 24 hours later, media was changed to fresh 100% unlabeled HPLM (10 mL for NHA Donor #1 in Figure S2A, 4.5 mL for NHA Donor #1 in Figure 5C and BT054). After 18 hours, media was harvested and prepared for LC-MS analysis as described above.

#### Doubling time quantification

NHA Donor #1 cells were plated in 6-well plates (0.05 × 10^6^ cells per well) in 2 mL DMEM prepared as described above. 24, 48, and 72 hours later, cells were counted using a Vi-CELL XR (Beckman Coulter, RRID: SCR_019664) cell viability analyzer. Doubling time was calculated using nonlinear regression to fit an exponential growth curve in GraphPad Prism (RRID: SCR_002798).

#### Adult mouse infusions

Infusions and tissue harvests were performed by researchers who were not blinded to the treatment arms or genotypes of the mice. No successfully infused mice were excluded from analysis. Mice that did not undergo orthotopic xenograft surgery were infused 1 to 7 weeks after receipt. Mice that did undergo orthotopic xenograft surgery were infused 34 days after surgery. Littermates of the same sex and xenograft status were randomized to tracer groups at the time of infusion. Amide-^15^N-glutamine (Cambridge Isotope Laboratories NLM-557) was prepared at 30 mg/mL in sterile normal saline and infused at a rate of 3 mg/kg/min. ^15^N_2_-uridine (Cambridge Isotope Laboratories NLM-812) was prepared at 3 mg/mL in sterile normal saline and infused at a rate of 0.3 mg/kg/min. 3 females and 1 male were infused per group of tracer and xenograft status. For saline infusions, 1 female and 1 male mouse per group of xenograft status were infused at a rate of 1 µL/g/min. The infusion apparatus (Instech Laboratories) included a swivel and tether to allow free movement around the cage. Mice were monitored constantly during infusion. During infusion, blood was collected by tail snip (∼5 µL) in blood collection tubes with K_2_EDTA (Becton Dickinson 366643). Mice were infused for 18 hours, after which mice were anesthetized with isoflurane and euthanized by decapitation. Tissues were quickly dissected and divided into two halves: one for MALDI MSI analysis, which was immediately frozen, and one for LC-MS analysis, which was dissected manually to isolate tumor region, washed in ice-cold normal saline, then frozen. Tissue and blood were stored at −80°C until analysis.

#### Fetal mouse infusions

Fetal mouse infusions were performed as described previously^102^. Healthy 12-week-old naïve pregnant female C57BL/6J mice were set up for mating between 05:00 and 07:00 with proven studs of the same genotype. The following morning, females displaying vaginal plugs were identified as pregnant and moved to a new cage until gestational day 11. Infusion took place between 09:00 and 11:00 with no prior fasting of pregnant dams. Mice were initially anaesthetized using intraperitoneal injection of ketamine and xylazine (120 mg/kg and 16 mg/kg, respectively) and maintained under anesthesia using subsequent intraperitoneal doses of ketamine (20 mg/kg) as needed. 25-gauge catheters were inserted into the tail vein and infusion begun immediately after an initial retro-orbital blood collection. Amide-^15^N-glutamine (Cambridge Isotope Laboratories NLM-557) was prepared as a total dose of 1.73 g/kg in 1,500 µL sterile normal saline infused at a rate of 147 μL/minute for 1 minute followed by 3 μL/minute for 5 hours. Retro-orbital blood was collected throughout the infusion to monitor tracer enrichment in maternal blood. Mice were euthanized at the end of the infusion, then the uterus was removed, and embryonic brain was dissected in cold sodium chloride irrigating solution (Baxter 2F7122) then snap frozen in liquid nitrogen. Tissue was stored at −80°C until analysis.

#### Tracing in human explants

Tumor tissue was collected using Brainlab Stereotactic Frameless Navigation Suite (Brainlab, RRID: SCR_026474). Tumor or brain tissue was collected from the operating room and immediately suspended in ice-cold Hibernate A (BrainBits HA). Within 30 minutes of explantation, tissue was transferred to RBC Lysis Buffer (Thermo Fisher 00433357) for 10 minutes with rocking. Next, tissue was washed three times with Hibernate A containing GlutaMAX (2mM, Thermo Fisher 35050061), penicillin/streptomycin (100 U/mL and 100 μg/mL, respectively, Thermo Fisher 15140148), and amphotericin B (250 ng/mL, GeminiBio 400104). Tissues were cut using dissection scissors or scalpels into 1-2 mm^3^ pieces and plated in a 24-well ultra-low adherence plate (Corning 3473) in 1 mL HPLM supplemented with B27 supplement minus vitamin A (1×, Thermo Fisher 12587010), N2 supplement (1×, Thermo Fisher 17502048), penicillin/streptomycin (100 U/mL and 100 μg/mL, respectively, Thermo Fisher 15140148), Plasmocin (250 ng/mL, InvivoGen ant-mpp), 2-mercaptoethanol (55µM, Thermo Fisher BP176100), and insulin (2.375-2.875 µg/mL, Millipore Sigma I9278). Tissues were randomized to groups of 1 to 4 technical replicates per tracer from each biologic sample, depending on available tissue. Plated tissues were placed in a 37°C incubator at ambient oxygen and 5% CO_2_ with shaking. Remaining tissue was used for histopathology as described below. After 30 minutes, media was exchanged for 1 mL HPLM supplemented as above with either glutamine exchanged for an equimolar amount of amide-^15^N-glutamine (Cambridge Isotope Laboratories NLM-557) or uridine exchanged for an equimolar amount of ^15^N_2_ uridine (Cambridge Isotope Laboratories NLM-812). Tissues were incubated for 18 hours in a 37°C incubator at ambient oxygen and 5% CO_2_ with shaking. After 18 hours, tissues were harvested, washed with ice-cold normal saline, and stored at −80°C until analysis. Tissues were then prepared for LC-MS analysis as described above.

#### Histopathology of human explants

Residual human tumor and brain tissues from parcellation for tracing experiments were fixed for 1 hour in neutral buffered 10% formalin solution (Millipore Sigma HT501128). Following fixation, tissues were washed and stored in 70% ethanol. Tissues were embedded in paraffin, sectioned, and stained with hematoxylin and eosin (H&E) by the UTSW Histo Pathology Core. H&E sections were reviewed by a board-certified neuropathologist (T.E.R).

#### Pyrimidine synthesis analysis in continuous cell culture with media change

NHA Donor #1 cells were plated on 6 cm dishes (0.8 × 10^6^ cells per dish) in 2 mL DMEM prepared as described above. 18 hours later, and every subsequent 6 hours for an additional 24 hours, media was changed to 2mL fresh DMEM. At indicated timepoints after media changes, media and cells were harvested and prepared for LC-MS analysis as described above. At the same time, a paired plate was counted with a Vi-CELL XR (Beckman Coulter, RRID: SCR_019664) cell viability analyzer.

#### Immunoblot analysis of protein expression

Cells were lysed in EBC lysis buffer containing either a protease inhibitor cocktail (Millipore Sigma 11836153001) or a protease and phosphatase inhibitor cocktail (Thermo Fisher 78440). Lysates were resolved by SDS-PAGE and transferred to nitrocellulose membranes (Bio-Rad 1620112). Primary antibodies used included: anti-IDH1 R132H (Dianova DIA-H09, 1:500, Mouse monoclonal, RRID: AB_2335716), anti-vinculin (Sigma V9131, 1:100,000, mouse monoclonal, RRID:AB_477629), anti-SOX2 (Cell Signaling Technologies 4900, 1:1000, mouse monoclonal, RRID: AB_10560516), anti-Phospho-CAD (Ser1859) (Cell Signaling Technologies 70307, 1:1,000, rabbit monoclonal, RRID: AB_2799782), anti-CAD (Cell Signaling Technologies 11933, 1:1,000, rabbit polyclonal, RRID: AB_2797772), anti-DHODH (Proteintech 14877, 1:1,000, rabbit polyclonal, RRID: AB_2091723), anti-UMPS (Millipore Sigma HPA036179, 1:1,000, rabbit polyclonal, RRID: AB_10673615), anti-UMPS (Proteintech 14830, 1:1,000, rabbit polyclonal, RRID: AB_2212392), and anti-HA (Thermo Fisher 26183, 1:1,000, mouse monoclonal, RRID: AB_10978021), anti-Phospho-S6 ribosomal protein (Ser240/244) (Cell Signaling Technologies 5364, 1:1,000, rabbit monoclonal, RRID: AB_10694233), anti-S6 ribosomal protein (Cell Signaling Technologies 2217, 1:1,000, rabbit monoclonal, RRID: AB_331355), anti-Phospho-p70 S6 kinase (Thr389) (Cell Signaling Technologies 9234, 1:1,000, rabbit monoclonal, RRID: AB_2269803), and anti-p70 S6 kinase (Cell Signaling Technologies 2708, 1:1,000, rabbit monoclonal, RRID: AB_390722). HRP-conjugated secondary antibodies used included: anti-Mouse IgG (Thermo Fisher 31430, 1:2,000, goat polyclonal, RRID: AB_228307) and anti-Rabbit IgG (Thermo Fisher 31460, 1:2,000, goat polyclonal, RRID: AB_228341).

#### Pyrimidine synthesis tracing in platelets

Human donor platelets (Charles River Platelet-1040) were received on dry ice and immediately thawed at 37°C with manual agitation. 1 mL of platelet suspension was washed in 9 mL HPLM supplemented with dialyzed FBS (10%, GeminiBio 100-108) and penicillin/streptomycin (100 U/mL and 100 μg/mL, respectively, Thermo Fisher 15140148), then centrifuged for 10 minutes at 300 × g at room temperature. Supernatant was aspirated, then platelets were resuspended in 5 mL unlabeled HPLM. Suspension was centrifuged for 15 minutes at 600 × g at room temperature. Supernatant was aspirated, and pellet was resuspended in 100 mL unlabeled HPLM. Platelets were counted using a Vi-CELL XR (Beckman Coulter, RRID: SCR_019664) cell viability analyzer and plated (2 × 10^6^ platelets per well) in 6-well plates in 10 mL HPLM. 24 hours later, each well was collected and centrifuged for 10 minutes at 600 × g at room temperature, then resuspended in 10 mL HPLM supplemented as above and modified as follows: glutamine exchanged for an equimolar amount of amide-^15^N-glutamine (Cambridge Isotope Laboratories NLM-557), glutamine exchanged for an equimolar amount of amide-^15^N-glutamine (Cambridge Isotope Laboratories NLM-557) and uridine removed, or uridine exchanged for an equimolar amount of ^15^N_2_ uridine (Cambridge Isotope Laboratories NLM-812). After incubation with tracer for indicated timepoints, samples were harvested and prepared for LC-MS analysis as described above.

#### Structure of human CAD

AlphaFold-predicted model^56,103^ of full-length human CAD protein (AF-P27708-F1-v4) was downloaded from the AlphaFold Protein Structure Database (https://alphafold.ebi.ac.uk, RRID: SCR_023662) and visualized using PyMol (3.1.1, Schrödinger, RRID: SCR_000305).

#### S1900 conservation analysis

The sequence of all proteins with >50% identity to the entry for human CAD (P27708) were downloaded from UniProt^68^. Of the 571 retrieved sequences, the longest isoform sequence available per species was retained to represent ortholog sequences in 289 unique species. 84 sequences that were shorter than 1,000 amino acids were removed from the analysis to exclude CAD fragments and orthologs that do not contain trifunctional CAD domains. The remaining full 205 sequences, 125 sequences containing class Mammalia in the taxonomy field, or 80 sequences not containing class Mammalia in the taxonomy field were aligned with Clustal Omega^92^ 1.2.4 (RRID: SCR_001591) implemented in SnapGene 8.1.1 (GSL Biotech, RRID: SCR_015052). Sequence logos were generated from alignment files with WebLogo^93,104^ 3.9.0 (https://weblogo.threeplusone.com) using probability units, a ± 6 amino acid window from human S1900, and the hydrophobicity color scheme.

### Quantification and Statistical Analysis

#### Analysis of nitrogen metabolism profiling platform data

Integrated peaks from LC-MS analysis of nitrogen metabolism profiling platform data were analyzed using R (RRID: SCR_001905). Metabolites which were not quantified in more than one sample, had a mean total pool size below an experiment-defined threshold (for AGI-5198 versus DMSO experiments: 2 × 10^6^, for NHA versus BT054 experiments: 1 × 10^5^), had calculated total labeling across all conditions of less than 1%, had quantified labeling of >3% in any unlabeled sample, had calculated total labeling across all conditions of greater than 500% were filtered and removed from subsequent analysis. Differential labeling scores were calculated using the formula: 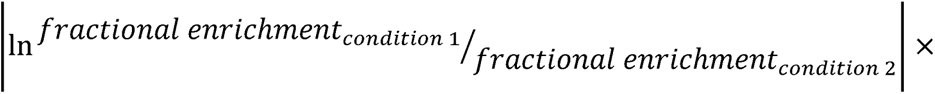 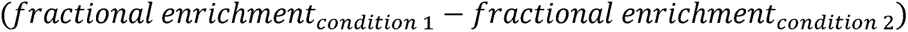. For visualization, metabolites with less than 5% total labeling were filtered and removed from Sankey diagrams. Sankey diagrams were generated using the R packages ggalluvial^94^ (RRID: SCR_021253) and ggplot^95^ (RRID: SCR_014601). Metabolites plotted as representative of individual pathways are available in Table S1.

#### CPTAC proteomic data analysis

For proteomic analysis of human brain and GBM tissues, raw phosphoproteome and proteome mass spectra converted to .mzML open standard format from the Clinical Proteomic Tumor Analysis Consortium (CPTAC) GBM discovery cohort were downloaded from the Proteomic Data Commons (https://pdc.cancer.gov, Study IDs: PDC000204, PDC000205, RRID: SCR_018273). FragPipe (RRID: SCR_022864) was used to analyze mass spectra. Peptides were identified using MSFragger^105–107^ using a 10 ppm parent ion tolerance, allowing for isotopic error in precursor ion selection, and searched against the UniProt human reference proteome UP000005640 retrieved on December 3, 2024, with reviewed sequences only, with 50% of the database representing decoys, and common contaminants added. Peptide search identified peptide cleavages for Lys-C and strict trypsin digestion, with a maximum of 2 missed cleavages. For the proteome dataset, carbamidomethylation on cysteine residues, TMT modification on the peptide N-terminus and lysine residues, and oxidation on methionine residues were considered. For the phosphoproteome dataset, the prior modifications as well as phosphorylation on serine, threonine, and tyrosine were considered. Peptide-to-spectrum matches were scored and validated using MSBooster^108^ and Percolator^109^. Proteins were inferred using ProteinProphet^110^ and false discovery rate filtering and reporting was performed using Philosopher^111^. A maximum false discovery rate at the peptide level of 1% was used. Protein groups covering 13,007 genes were identified in the proteome dataset and protein groups covering 9,124 genes were identified in the phosphoproteome dataset. Isobaric quantification was performed using TMT Integrator^112^, with comparison of each sample to the pooled reference sample labeled with TMT 126 reagent. Phosphopeptide abundances were normalized to the total abundance of each protein.

For LC-MS/MS tandem spectrum generation, files were analyzed using Proteome Discoverer (3.0, Thermo Fisher, RRID: SCR_014477), with peptide identification performed using Sequest HT searching against the human reviewed protein database from UniProt (downloaded January 4, 2025, 20354 entries). Fragment and precursor tolerances of 10 ppm and 0.6 Da were specified, and three missed cleavages were allowed. Carbamidomethylation of Cys was set as a fixed modification, with oxidation of Met and phosphorylation of Ser, Thr, and Tyr set as variable modifications. The false-discovery rate (FDR) cutoff was 1% for all peptides. Peptide peak intensities were used as peptide abundance values.

#### Mouse proteome data analysis

For proteomic analysis of mouse tissues, C57BL/6N mouse tissue-specific proteome and phosphoproteome data were obtained from Giansanti et al.^85^ log_10_-transformed abundance of tissue-specific total Cad and Cad phosphopeptide levels were transformed to raw peptide intensities. Phosphopeptide levels were normalized to total Cad abundance within each tissue, then log_10_ transformed. Total Cad, Umps, Dhodh, Uck1, Uck2, and Uprt tissue-specific abundance were plotted as log_10_-transformed values.

#### Other statistical analyses

Information related to data presentation and statistical analysis for individual experiments can be found in the corresponding figure legends. Statistical analyses were carried out using GraphPad Prism (RRID: SCR_002798). For all experiments, *n* represents biological replicates rather than technical replicates. All experiments involving primary tissues (mouse infusions, human explants) are derived from at least three independent experiments. Student’s unpaired, two-tailed *t*-tests were used for pairwise comparisons. For each *t*-test, an F test was performed to compare variances between groups. If the variances differed significantly (F test *p* < 0.05), Welch’s correction was applied. One-way ANOVA tests were performed for significance comparison of three or more groups. Linear regression analysis was used to test statistical significance of correlations. To model metabolite labeling over time, curves were fit to labeling data by nonlinear regression analysis using a sigmoidal dose-response (variable slope) model. Statistical significance is indicated as follows: n.s. = not significant, ∗*p* < 0.05, ∗∗*p* < 0.01, ∗∗∗*p* < 0.001. Error bars represent the standard error of the mean (SEM).

## RESOURCE AVAILABILITY

### Lead Contact

- Further information and requests for resources and reagents should be directed to and will be fulfilled by the Lead Contact, Samuel K. McBrayer.

### Materials Availability

- Plasmids generated in this study will be deposited to Addgene prior to publication.

### Data and Code Availability

- Metabolomics data will be deposited to the National Metabolomics Data Repository (NMDR) and will be publicly available as of the date of publication. Accession and identification numbers will be listed in the key resources table.
- Original code will be deposited on GitHub and will be publicly available as of the date of publication. Accession number will be listed in the key resources table.
- All data and additional information required to reanalyze the data reported in this paper is available from the lead contact upon request.

## SUPPLEMENTAL INFORMATION

### SUPPLEMENTAL FIGURE LEGENDS

**Figure S1. Development of a nitrogen metabolism profiling platform and differential labeling score metric, related to Figure 1. (A)** Heatmap depicting ^15^N fractional enrichment (FE) (1-M+0) in multiple metabolite pools (y-axis) in each ^15^N-labeled HPLM stocks (x-axis) (*n* = 1 per tracer). **(B)** Total metabolite levels in HPLM stocks (*n* = 30). **(C)** FE of cognate intracellular metabolite pool for each tracer in BT054s treated with 3µM AGI-5198 or DMSO for 72 hours, followed by tracing for 18 hours (*n* = 1 per tracer per treatment). **(D)** Schema depicting nitrogen metabolism profiling platform analytical pipeline and derivation of differential labeling score. **(E-G)** Peak intensity of (E) 2-hydroxyglutarate (2HG), (F) glutamate, or (G) oxidized glutathione from nitrogen metabolism profiling of BT054s treated with 3µM AGI-5198 or DMSO for 72 hours, followed by tracing for 18 hours (*n* = 30 per treatment). **(H-I)** ^15^N FE from indicated tracers in (H) glutamate or (I) reduced glutathione (GSH) pools in BT054s treated with 3µM AGI-5198 or DMSO for 72 hours, followed by tracing for 18 hours (*n* = 3). n.s. = not significant, \*\**p* < 0.01, \*\*\**p* < 0.001. Two-tailed *p* values were determined by unpaired *t*-test. For all panels, data are means ± SEM.

**Figure S2. Nitrogen metabolism profiling and evaluation of pyrimidine synthesis pathways in neural cell cultures, related to Figure 2. (A)** Metabolite levels in conditioned HPLM samples after BT054 or NHA Donor #1 cell culture for 18 hours. Data are normalized to unconditioned HPLM (*n* = 3 per cell line). **(B)** FE of cognate intracellular metabolite pool for each tracer in BT054 or NHA Donor #1 cells at 18 hours (*n* = 1 per tracer per cell line). **(C)** Volcano plot of metabolites in BT054 or NHA Donor #1 cells (*n* = 30). **(D)** Doubling times of NHA Donor #1 and BT054 cells (*n* = 3 per cell line). Doubling times for BT054 are from Shi et al., 2022^14^. **(E-K)** Peak intensity of indicated metabolites from nitrogen metabolism profiling of BT054 or NHA Donor #1 cells (*n* = 30 per cell line). **(L)** Labeling from indicated tracer to intracellular metabolite pool in GSCs, differentiated cells, and NSCs/NPCs at 18 hours (*n* = 3 for all lines except HK308, for which *n* = 2). **(M)** Amide-^15^N-glutamine and ^15^N_2_-uridine tracing to UMP in BT054 cells cultured in HPLM with dialyzed FBS (*n* = 3). **(N)** Amide-^15^N-glutamine and ^15^N_2_-uridine tracing to UMP in NHA Donor #1 cells cultured in HPLM without dialyzed FBS and with epidermal growth factor (EGF) and basic fibroblast growth factor (bFGF) (*n* = 3). **(O-P)** Amide-^15^N-glutamine and ^15^N_2_-uridine tracing to UMP in (O) ENSA and (P) NSC11 NSC lines grown in adherent culture conditions. In (M-P) All amide-^15^N-glutamine to UMP labeling data represent UMP M+1 FE normalized to glutamine M+1 FE. All ^15^N_2_-uridine to UMP labeling data represent UMP M+2 FE. n.s. = not significant, \*\**p* < 0.01, \*\*\**p* < 0.001. Two-tailed *p* values were determined by unpaired *t*-test for all panels except (L), for which two-tailed *p* values were determined by ordinary one-way ANOVA. For all panels, data are means ± SEM.

**Figure S3. Validation of in vivo labeling patterns, related to Figure 3. (A)** Glutamine levels in mouse blood (*n* = 4 for saline infusion, *n* = 8 for amide-^15^N-glutamine and ^15^N_2_-uridine infusions). **(B)** Glutamine levels in mouse brain after 18 hour infusions (*n* = 2 for saline infusion, *n* = 3 for amide-^15^N-glutamine infusion, *n* = 4 for ^15^N_2_-uridine infusion). **(C)** Uridine levels in mouse blood (*n* = 4 for saline infusion, *n* = 8 for amide-^15^N-glutamine and ^15^N_2_-uridine infusions). **(D)** Uridine levels in mouse brain after 18 hour infusions (*n* = 2 for saline infusion, *n* = 4 for amide-^15^N-glutamine and ^15^N_2_-uridine infusions). **(E)** Schema of cytidine triphosphate synthetase (CTPS1)-dependent nitrogen transfer in amide-^15^N-glutamine stable isotope tracing assays. ATP = adenosine triphosphate. ADP = adenosine diphosphate. CMPK1 = cytidine/uridine monophosphate kinase 1. NDPK = nucleoside-diphosphate kinase. CTP = cytidine triphosphate. PCYT2 = phosphate cytidylyltransferase 2. CDP-ethanolamine = cytidine diphosphate ethanolamine. **(F)** M+0 and **(G)** M+1 FE in CDP-ethanolamine in mouse brain at 18 hours (*n* = 1 for saline infusion, *n* = 3 for amide-^15^N-glutamine infusion). **(H)** M+0 and **(I)** M+1 FE in phosphoethanolamine in mouse brain at 18 hours (*n* = 1 for saline infusion, *n* = 4 for amide-^15^N-glutamine infusion). n.d. = not detected. n.s. = not significant. Two-tailed *p* values were determined by ordinary one-way ANOVA. For all panels, data are means ± SEM.

**Figure S4. Validation of explant labeling patterns, related to Figure 4. (A)** Glutamine M+1 FE in explants incubated in unlabeled (*n* = 3 for brain cohort, *n* = 4 for glioma cohort), amide-^15^N-glutamine (*n* = 5 per cohort), or ^15^N_2_-uridine (*n* = 5 per cohort) HPLM for 18 hours. **(B)** Uridine M+2 FE in explants incubated in unlabeled (*n* = 3 for brain cohort, *n* = 4 for glioma cohort), amide-^15^N-glutamine (*n* = 5 per cohort), or ^15^N_2_-uridine (*n* = 5 per cohort) HPLM for 18 hours. n.s. = not significant, \*\**p* < 0.01. Two-tailed *p* values were determined by unpaired *t*-test. For all panels, data are means ± SEM.

**Figure S5. Uridine deprivation acutely activates de novo pyrimidine synthesis in differentiated cells through a post-transcriptional mechanism, related to Figure 5. (A-C)** Quantification of intracellular (A) UMP, (B) UDP-hexose, and (C) UDP-HexNAc levels in NHA Donor #1 cells cultured in DMEM with 3µM uridine. Media was changed at timepoints indicated by arrows. Curves are fit by nonlinear regression for full 24-hour experiment (*n* = 3). **(D)** Heatmap of metabolite levels in NHA Donor #1 cells cultured in 0µM or 3µM uridine for 0.25 to 9 hours after 24 hours of culture in 3µM uridine (*n* = 6 per group). Data are log_2_-transformed peak intensities normalized to 0-hour timepoint for each metabolite. **(E-F)** Ratio of the area under the curve of (E) UMP M+1 or (F) carbamoyl aspartate M+1 after 18-hour amide-^15^N-glutamine tracing in 0µM versus 3µM uridine. Cells were preconditioned in 3µM uridine for 24 hours. **(G-L)** Protein expression of (G) CAD, (H) DHODH, (I) UMPS, (J) UCK1, (K) UCK2, and (L) UPRT1 in human brain and glioblastoma (GBM) samples (*n* = 10 brain, *n* = 99 GBM). **(M)** Heatmap depicting relative abundance of *CAD*, *DHODH*, *UMPS*, *UCK1*, *UCK2*, and *UPRT* mRNA in a panel of glioma stem-like cells (GSCs) and NHA Donor #1 (NHA) cells. Data are log_2_-transformed, median-normalized fragments per kilobase of transcript per million mapped reads (FPKM) from RNA sequencing. Abundances are results of reanalysis of raw data published in Wu et al., 2024^28^ and Savani et al., 2025^46^. **(N)** Carbamoyl aspartate M+1 peak intensity after amide-^15^N-glutamine tracing in NHA Donor #1 cells cultured in 0µM (blue) or 3µM (gray) uridine. Cells were preconditioned in 3µM uridine for 24 hours (*n* = 6 per group). * compares 3-hour timepoint. **(O)** Representative immunoblots for CAD, DHODH, and UMPS in HeLa cells engineered to express Cas9 and *AAVS1* safe harbor locus control, *CAD*, *DHODH*, or *UMPS* sgRNAs. **(P-R)** Representative immunoblots for CAD, DHODH, and UMPS in (P) NHA Donor #1, (Q) NHA Donor #2, or (R) BJ cells cultured in 0µM or 3µM uridine for indicated times after 24 hours of culture in 3µM uridine. **(S-U)** Normalized peak intensities of (S) carbamoyl aspartate M+1, (T) UTP, or (U) uric acid in NHA Donor #1 cells cultured in 0µM (blue) or 3µM (gray) uridine for 1 hour after 24 hours of culture in 3µM uridine (*n* = 6). **(V-X)** Normalized peak intensities of (V) carbamoyl aspartate M+1, (W) UTP, or (X) uric acid in NHA Donor #2 cells cultured in 0µM (blue) or 3µM (gray) uridine for 6 hours after 24 hours of culture in 3µM uridine (*n* = 3). **(Y)** Representative immunoblots for S1859 phosphorylation site on CAD, total CAD, the S240/S244 phosphorylation site of S6 ribosomal protein, total S6 ribosomal protein, the T389 phosphorylation site on S6 kinase (S6K), and total S6K in NHA Donor #1 cells treated for 1 hour with DMSO or 100µM torin. **(Z-CC)** Representative immunoblots for S240/S244 phosphorylation site of S6 ribosomal protein, total S6 ribosomal protein, the T389 phosphorylation site on S6K, and total S6K in (Z-AA) NHA Donor #1, (BB) NHA Donor #2, or (CC) BJ cells cultured in 0µM or 3µM uridine for indicated times after 24 hours of culture in 3µM uridine. **(DD-GG)** Representative immunoblots for S1859 phosphorylation site on CAD and total CAD in (DD-EE) NHA Donor #1, (FF) NHA Donor #2, or (GG) BJ cells cultured in 0µM or 3µM uridine for indicated times after 24 hours of culture in 3µM uridine. n.s. = not significant, \**p* < 0.05, \*\**p* < 0.01, \*\*\**p* < 0.001. In (E-F), two-tailed *p* values were determined by ordinary one-way ANOVA. For all other panels, two-tailed *p* values were determined by unpaired *t*-test. For all panels, data are means ± SEM.

**Figure S6. CAD S1900 phosphorylation is enriched in primitive cell populations, related to Figure 6. (A-C)** LC-MS/MS tandem mass spectrum for the peptides (A) RLpSSFVTK, corresponding to phosphorylation at S1406, (B) IHRApSDPGLPAEEPK, corresponding to phosphorylation at S1859, or (C) QApSPQNLGTPGLLHPQTSPLLHSLVGQHILSVQQFTK, corresponding to phosphorylation at S1900, of full-length CAD protein from human brain and glioblastoma (GBM) samples. **(D)** Heatmaps depicting protein expression of Cad, Umps, Dhodh, Uck1, Uck2, and Uprt in mouse tissues. Data are log_10_-transformed protein abundance in each tissue. Abundances are results of reanalysis of data published in Giansanti et al., 2022^85^. **(E-V)** Correlation between (E-J) S1406, (K-P) S1859, and (Q-V) S1900 phosphorylation of CAD and (E, K, Q) GSK3β phosphorylation at S9, (F, L, R) S6 ribosomal protein phosphorylation at S235/S236, (G, M, S) OLIG2 protein levels, (H, N, T) SOX2 protein levels, (I, O, U) ERK1 phosphorylation at T202/Y204, or (J, P, V) ERK2 phosphorylation at T185/Y187. For (E-V), phosphorylation data are normalized to total protein and all data are mean normalized. Correlation coefficients and *p* values in red indicate statistically significant positive correlation, blue indicate statistically significant negative correlation, and black indicate lack of significant correlation (*n* = 99). *r* = Pearson’s correlation coefficient. *p* values were determined by simple linear regression analysis. Panels (A-C and E-V) are results of reanalysis of raw data published in Wang et al., 2021^47^.

**Figure S7. Validation of CAD cDNA expression, related to Figure 7. (A)** Representative immunoblots of total CAD and HA epitope on exogenous CAD proteins in NHA Donor #1 cells engineered to express either EV, WT CAD, or indicated CAD mutants. **(B)** Representative immunoblots of total CAD and HA epitope on exogenous CAD proteins in G9C CHO cells engineered to express either EV, WT CAD, or indicated CAD mutants. Endogenous CAD expression in parental NHA Donor #1 cells is shown for comparison.

