## Supplementary figures and images for "Nitrogen metabolism profiling reveals cell state-specific pyrimidine synthesis pathway choice"

### Figure S1

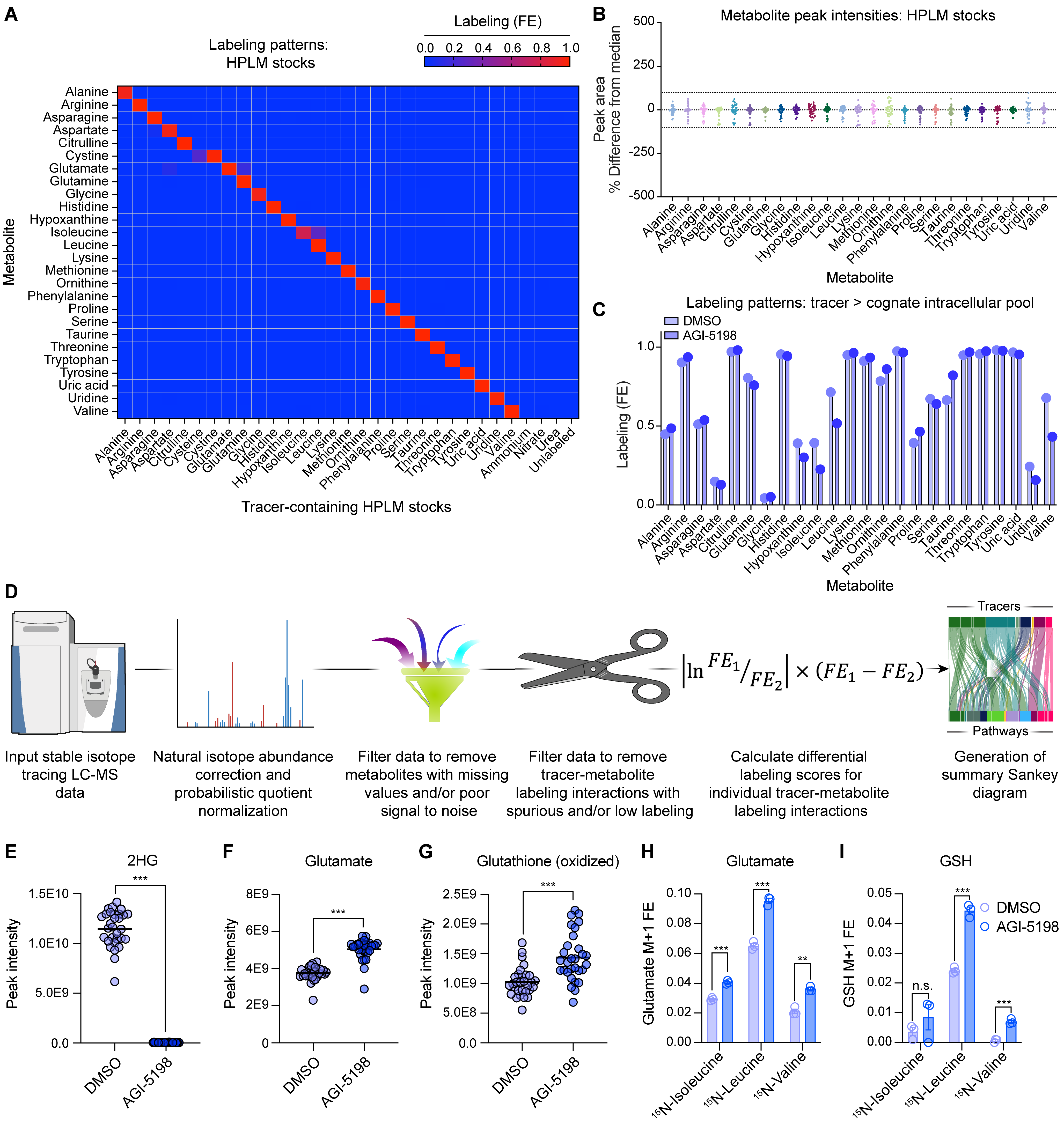

### Figure S2

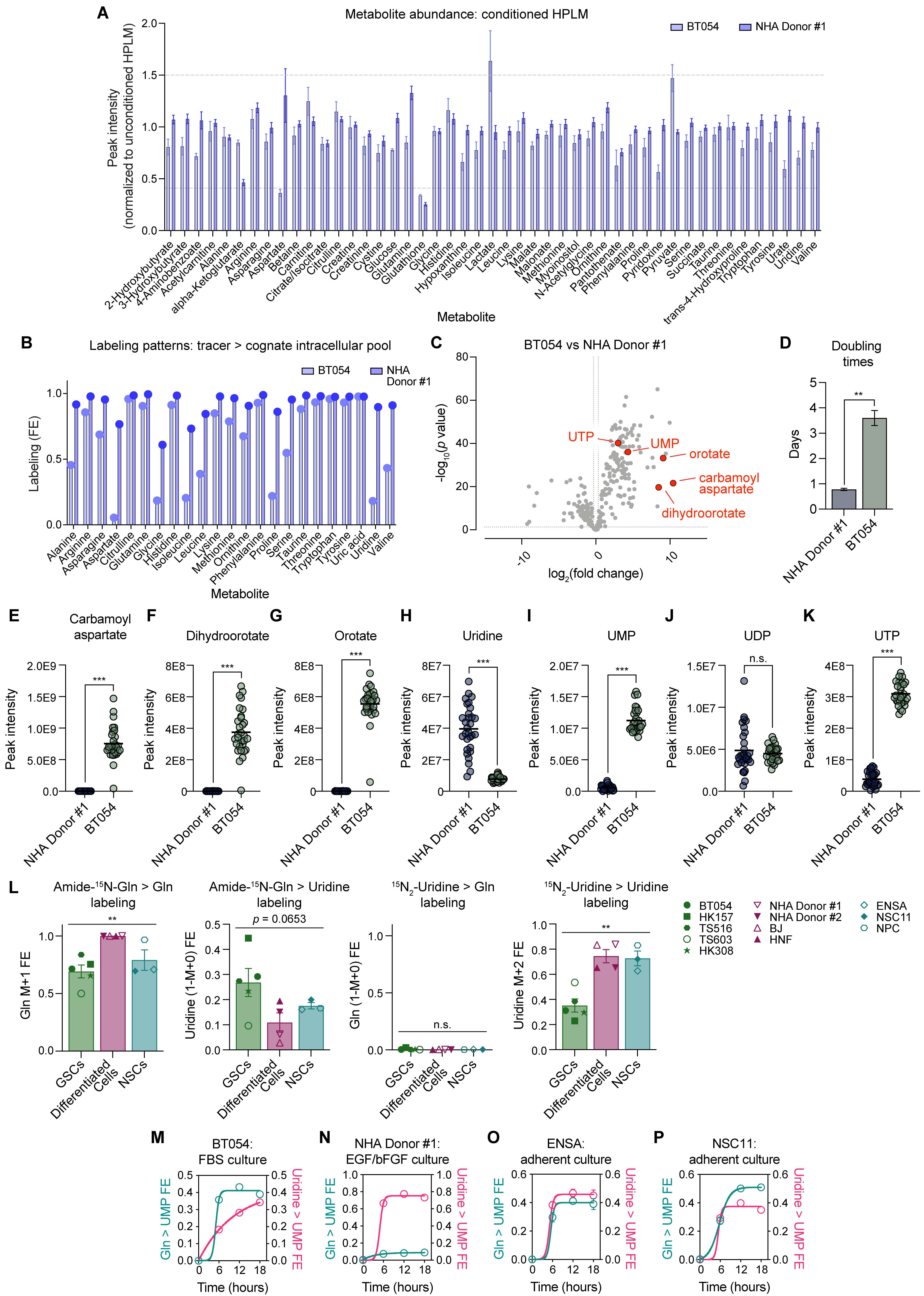

### Figure S3

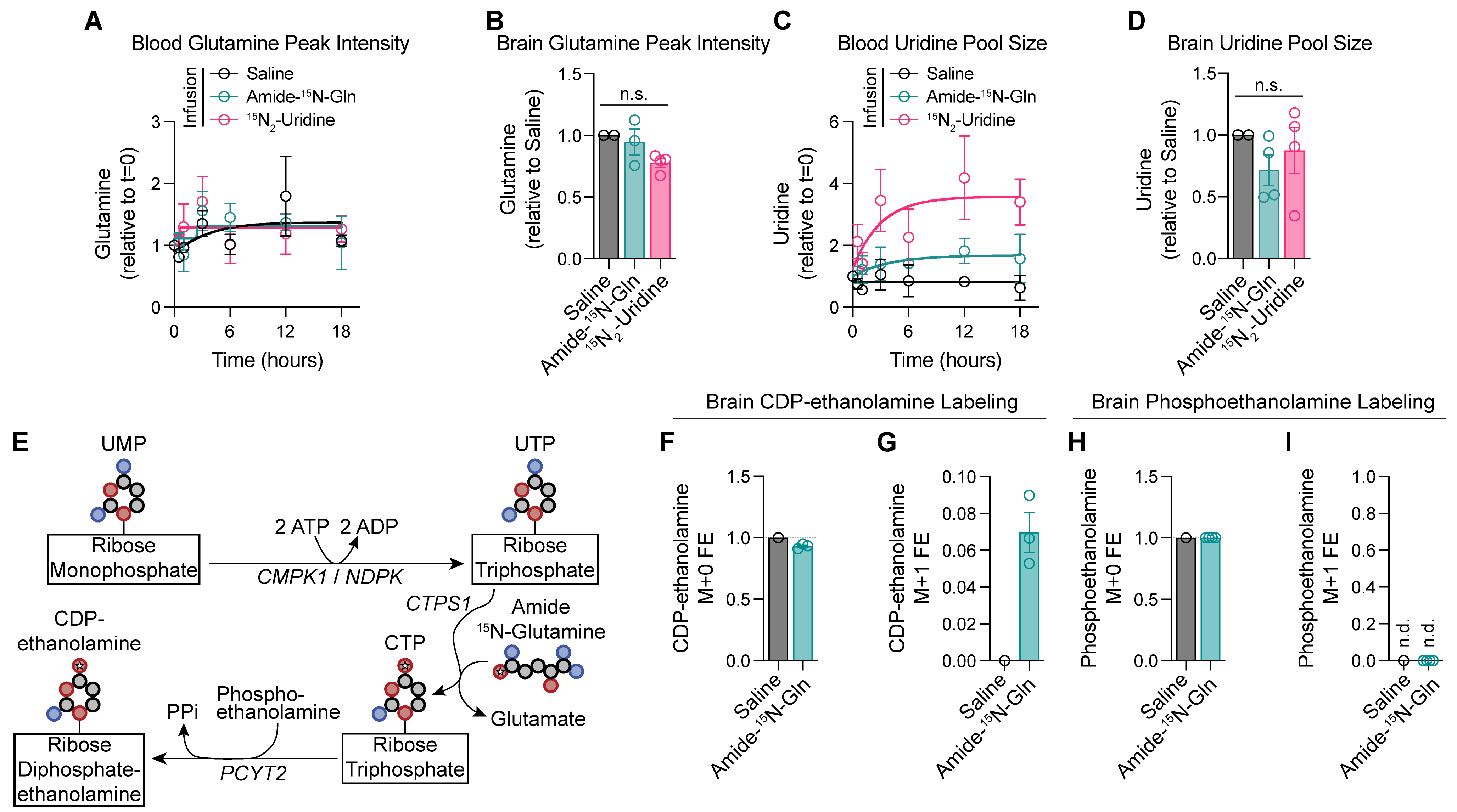

### Figure S4

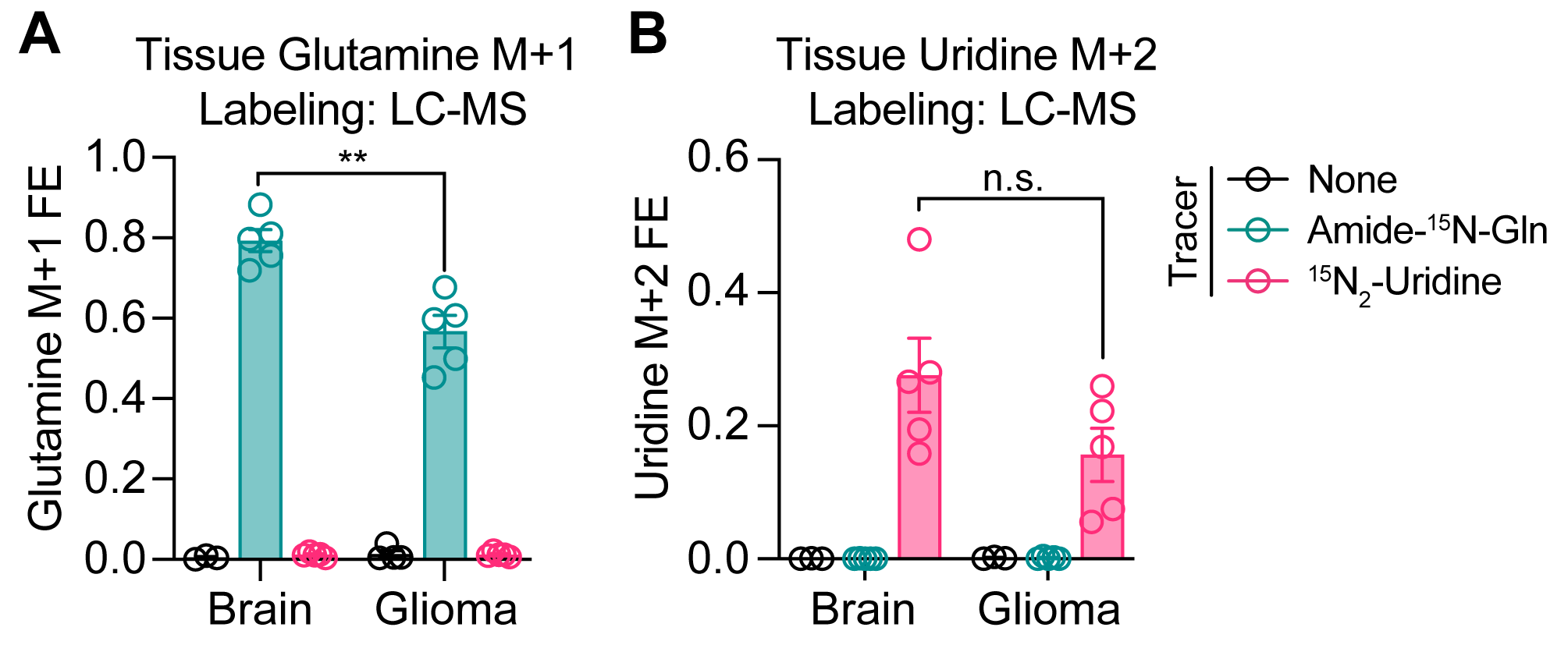

### Figure S5

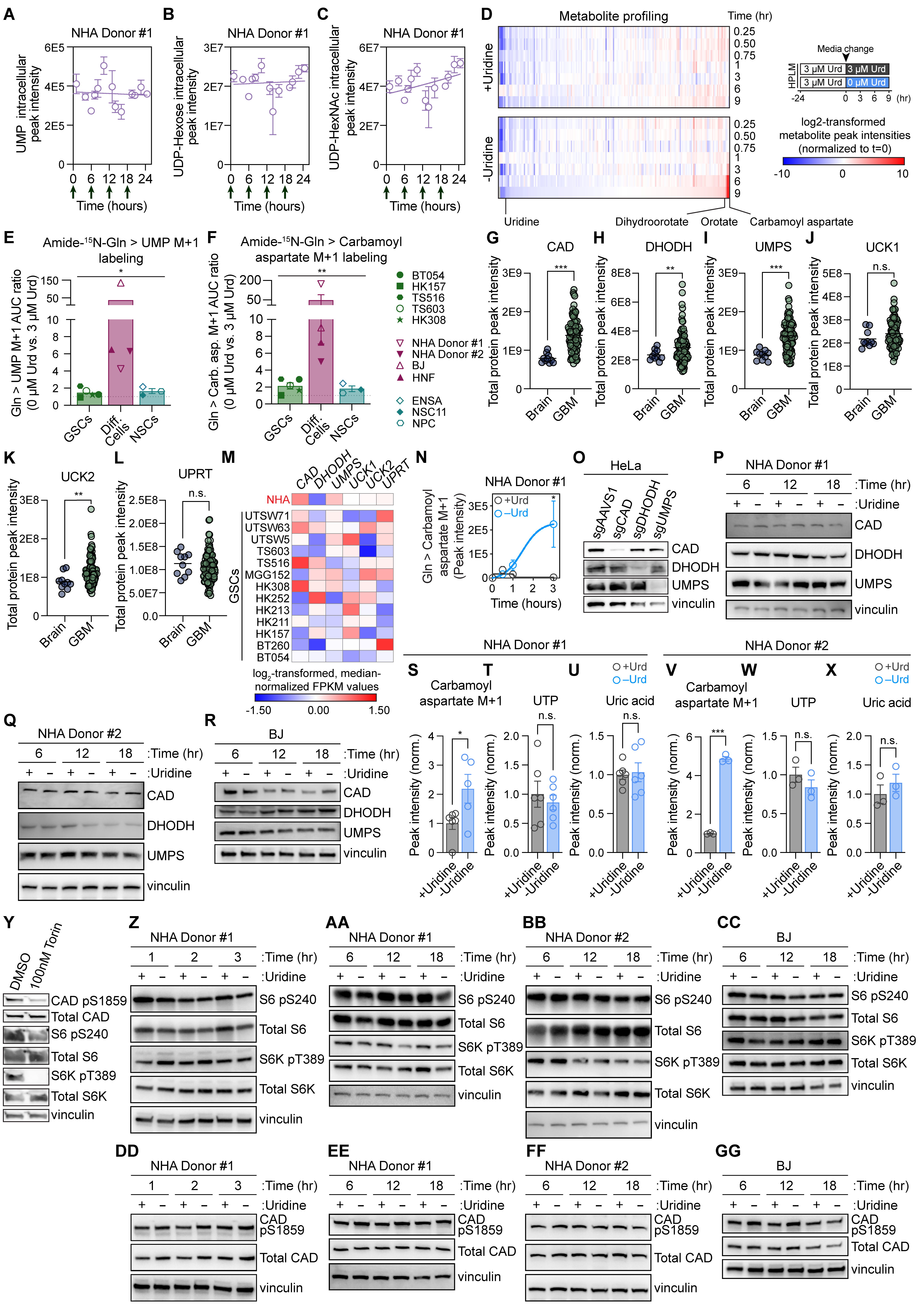

### Figure S6

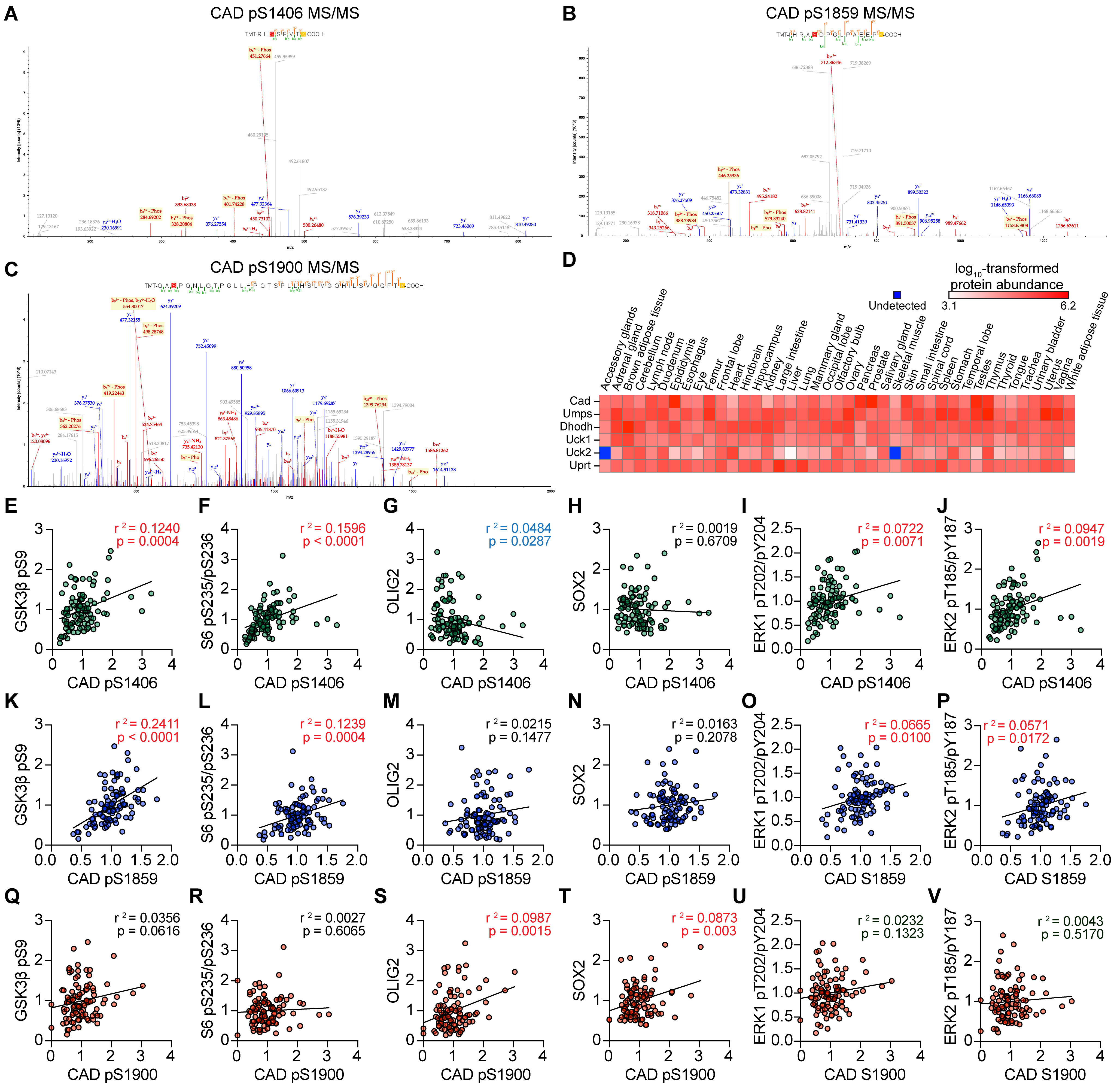

### Figure S7

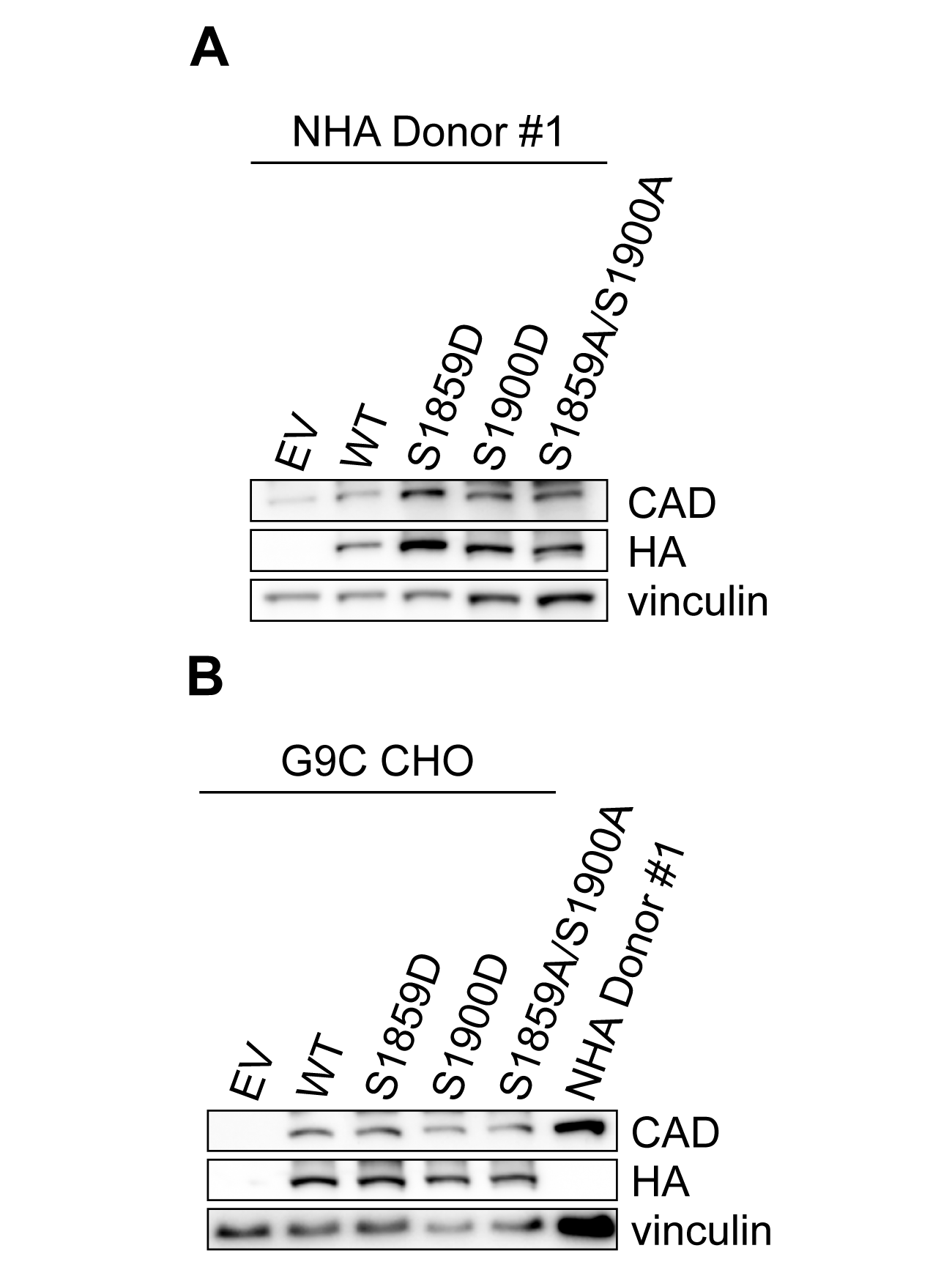
